# Evolution-Aware Deep Reinforcement Learning for Single-Cell DNA Copy Number Calling

**DOI:** 10.1101/2024.03.08.583988

**Authors:** Stefan Ivanovic, Mohammed El-Kebir

## Abstract

Recent high-throughput single cell DNA sequencing technologies enable one to detect copy number aberrations (CNAs) on thousands of individual cells within a tumor. Several new methods enable haplotype-specific copy number calling on such data. However, these algorithms do not utilize evolutionary constraints, leading to the inference of spurious CNAs due to measurement noise that are inconsistent with a realistic evolutionary model. To address this gap, we introduce DeepCopy, an evolution-aware deep reinforcement learning algorithm for haplotype-specific copy number calling on single-cell DNA sequencing data. On simulated data, DeepCopy’s predictions better fit the ground truth than existing methods. As shown on 10x Genomics Single Cell CNV sequencing of several breast cancers and DLP+ sequencing of an ovarian cancer, DeepCopy’s joint estimation of copy number profiles and evolutionary model parameters results in larger, more realistic clones of cells with identical copy number profiles that lead to more parsimonious CNA phylogenies than existing methods, while retaining agreement with truncal single-nucleotide variants. Finally, on a breast cancer patient sequenced with both ACT and 10x technologies, we demonstrate the consistency of DeepCopy’s predictions across sequencing technologies.

## 1 Introduction

Cancer results from an evolutionary process during which somatic mutations accumulate in a population of cells, resulting in intra-tumor heterogeneity, i.e., the presence of multiple cellular subpopulations, or *clones*, with distinct mutations [1]. Copy number aberrations (CNAs) are a very common form of mutation in cancer, on average affecting 44% of the genome in solid tumors [2–4]. Unlike single-nucleotide variants (SNVs), individual CNAs can simultaneously affect thousands of genes by amplifying or deleting large regions of the genome. Consequently, the identification of CNAs is vital for understanding cancer evolution. Additionally, identifying CNAs is of great clinical utility, since there exist certain drug therapies that target vulnerabilities in the cancer that arise as a result of specific CNAs [5–7]. However, the combination of intra-tumor heterogeneity and limitations of current sequencing technologies render copy number calling a challenging task.

Several works have focused on copy number calling from bulk DNA sequencing data [8–14]. Two key signals are used for copy number calling in these data. First, the number of reads, or *read depth*, of a genomic region is proportional to its total number of copies. Second, the *B-allele frequency* (BAF) of heterozygous germline SNPs is indicative of allelic imbalance, where a BAF of 0 (1) indicates the absence of B (A) alleles and a BAF of 0.5 indicates the same number of A and B alleles. A key limitation of these data is that we do not measure read depth and BAF in individual clones but rather obtain composite measurements of millions of cells in the bulk sample. This composite signal obscures intra-tumor heterogeneity and requires deconvolution, resulting in an inevitable loss of signal. Unlike bulk sequencing, single cell sequencing technologies potentially enable the precise characterization of copy number in individual cells. Due to the popularity of single cell RNA sequencing, a wide variety of algorithms exist for determining CNAs on these data such as Numbat [15], CopyKAT [16], HoneyBadger [17] and InferCNV [18]. However, as read depth in these data is additionally confounded by expression, it is extremely difficult to precisely determine integer copy numbers, and, instead, only the general presence of deletions and amplifications is predicted.

By contrast, single-cell DNA sequencing technologies have enabled the sequencing of hundreds of cells at relatively uniform coverage, suitable for copy number calling [23]. Algorithms such as HMMcopy [24], Ginkgo [25], SCICoNE [26], SCNA [27] and SCOPE [28] have been utilized to determine total copy numbers of regions on the genome on this single cell data. More recent technologies, including 10x Genomics CNV solution [19], DLP+ [20, 21] and ACT [22], have improved upon first-generation technologies to enable high-throughput single-cell DNA sequencing of thousands of cells with lower error rates. Three recent methods, CHISEL [29] SIGNALS [30], and Alleloscope [31], have enabled the determination of *haplotype-specific copy numbers* on this newer type of single cell data, indicating the number of copies of both parental haplotypes phased within each chromosome. CHISEL works by clustering bins by read depth and BAF, and then jointly inferring the copy number of these clustered bins. SIGNALS first applies HMMcopy to obtain total copy number estimates and then utilizes the BAF measurement to determine the haplotype-specific state. Alleloscope first performs segmentation on bulk or pseudobulk data and then performs phasing and copy number calling. These algorithms have detected additional patterns on single-cell sequencing data not observable from total copy numbers such as copy-neutral loss of heterozygosity and mirrored-subclonal CNAs. However, these algorithms have not taken advantage of biological constraints imposed by the evolutionary nature of cancer to jointly estimate copy number profiles across cells. The temporal structure of these biological constraints can act as a regularizer, reducing the number of spurious CNAs predicted due to noise in the data, as discussed in [26, 32, 33].

To address this gap, we introduce DeepCopy, an evolution-aware deep reinforcement learning algorithm for haplotype-specific copy number calling. DeepCopy models the formation of copy number profiles with an evolutionary process where CNAs sequentially occur on copy number profiles starting with a normal cell (Fig. 1). We use deep reinforcement learning to generate copy number profiles according to this evolutionary process and guide this evolutionary process toward producing copy number profiles that maximize the probability of the observed read depth and BAF data. DeepCopy does not rely on the presence of normal cells, enabling the analysis of data where only tumor cells have been sequenced through FACS sorting. We compare DeepCopy with SIGNALS and CHISEL on simulated data, finding DeepCopy’s predictions better fit ground truth. On breast and ovarian cancers [20, 22], DeepCopy produces larger clones, fewer unique copy number profiles, and ultimately enables the inference of more parsimonious phylogenies than competing methods, while maintaining agreement with orthogonal SNV data. Additionally, running DeepCopy on a breast cancer patient sequenced with both 10x and ACT technologies demonstrates the consistency of DeepCopy’s predictions independent of sequencing technology. We expect that DeepCopy’s more accurate copy number analyses will enable more precise analyses of intra-tumor heterogeneity.

**Figure 1.**
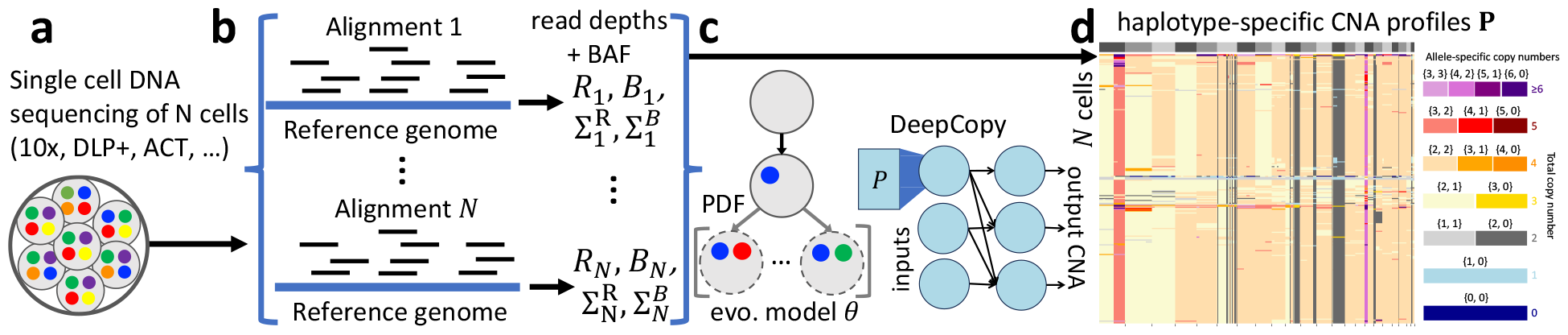
Overview of DeepCopy. **a**, Technologies such as 10x Genomics CNV solution [19], DLP+ [20, 21] and ACT [22] enable high-throughput single-cell DNA sequencing suitable for CNA calling. **b**, Reads are aligned and processed to obtain DeepCopy inputs. **c**, DeepCopy uses a generative model of CNA evolution parameterized by a convolutional neural network and optimized via reinforcement learning. **d**, From the evolutionary model *θ*, DeepCopy produces haplotype-specific copy number profiles **P**.

## 2 Results

### 2.1 DeepCopy algorithm

The objective of DeepCopy is to estimate the parameters of an evolutionary model of CNAs that maximize the data probability of observed read counts. Then, given the model, we predict a copy number profile for each individual cell with maximum joint probability according to said evolutionary model and observed read counts (Section 4.1). To accomplish this, DeepCopy’s pipeline follows three stages: (i) data processing, (ii) initial copy number estimation, and (iii) final copy number estimation via deep reinforcement learning.

First, in the data processing stage, the input is a BAM file with *N* read groups indicating cell barcodes and the outputs are read depths 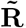, haplotype-specific read counts **A**, and variance estimates **Σ**^*R*^ and **Σ**^*B*^ for the read depth and B-allele frequency, respectively, of *L* segments in the genome (Section 4.2). To do so, we perform phasing and initially split the genome into *K* bins of length 100kb in order to perform bias correction and segmentation. Second, our NaiveCopy algorithm performs initial copy number estimation separately for each cell, yielding approximate copy number numbers profiles 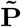 (Section 4.2). To accomplish this for each cell, NaiveCopy uses low-noise regions to estimate the cell-specific scaling factor and then determines the copy number profile that maximizes the probability of that cell’s read counts. Third, our evolution-aware deep reinforcement learning system inputs 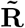, **A, Σ**^*R*^, **Σ**^*B*^, and 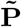 in order to jointly estimate copy number profiles **P** for all cells in order to maximize the probability of the observed read count data (Section 4.3). In contrast to existing methods, this joint estimation is evolution-aware, and based on a generative model for copy number profiles that sequentially adds CNAs to clones starting from the normal clone. The reinforcement learning system views the addition of mutations to clones as actions and utilizes a reward that maximizes the probability of the observed read counts given the model.

### 2.2 Evaluation on simulated scDNA-seq data

To benchmark our method on simulated data, we developed a fitness-aware simulator that generates copy number profiles **P** in addition to read depths 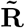 and haplotype-specific count measurements **A**. Based on the noise levels of breast cancer patient S0 (Fig. S1 and Section 2.4), we generated 20 simulation instances with *n* = 1000 cells and *K* = 27,283 bins of 100kb length that had varying levels of fitness rate increases and half of which included a truncal whole-genome duplication (WGD). Briefly, the simulator generates evolutionary trees starting with either a normal clone or a whole genome duplicated clone, and then sequentially adds CNAs to existing clones, with each CNA potentially increasing the corresponding clone’s fitness and thus the number of cells in the sample that originate from that clone or its descendants. As our simulations generate intermediate data rather than BAM or FASTQ files, we were not able to include Alleloscope in our benchmarking and only compared DeepCopy to SIGNALS and CHISEL. Further details are provided in Supplementary Section A.4.4, and runtimes are provided in Fig. S2.

Visually inspecting the copy number profiles of a single simulation instance with WGD (Fig. 2a), we find that DeepCopy’s predictions better match the ground truth compared to SIGNALS and CHISEL. To better quantify performance, we developed four different metrics. First, we defined *accuracy* as the average proportion of bins across all cells for which the predicted allele-specific copy number matches the true (un-ordered) allele-specific copy number (defined precisely in Section 4.4). For the shown simulation instance, DeepCopy’s accuracy was 0.948 compared to 0.704 for SIGNALS and 0.816 for CHISEL (Fig. 2a). We see similar trends across all simulation instances, with DeepCopy achieving a median accuracy of 0.956, followed by 0.882 for SIGNALS and 0.857 for CHISEL (Fig. 2b). Second, to better assess the magnitude of errors, we defined *error* as the mean absolute difference between predicted and ground-truth copy numbers across all bins and cells (again allele-specific as precisely defined in Section 4.4). DeepCopy achieved the smallest error of 0.055 for the shown simulation instance (Fig. 2a) and a median error of 0.064 across all simulation instances (Fig. 2c), followed by CHISEL (0.233 and 0.191, respectively) and then SIGNALS (0.470 and 0.315, respectively). In Supplementary Section B.2 and Fig. S3, we additionally analyzed sensitivity to focal CNAs (including CNAs smaller than 5Mb) and error on haplotype-specific copy numbers, finding DeepCopy to have lower errors than SIGNALS or CHISEL in each circumstance.

**Figure 2.**
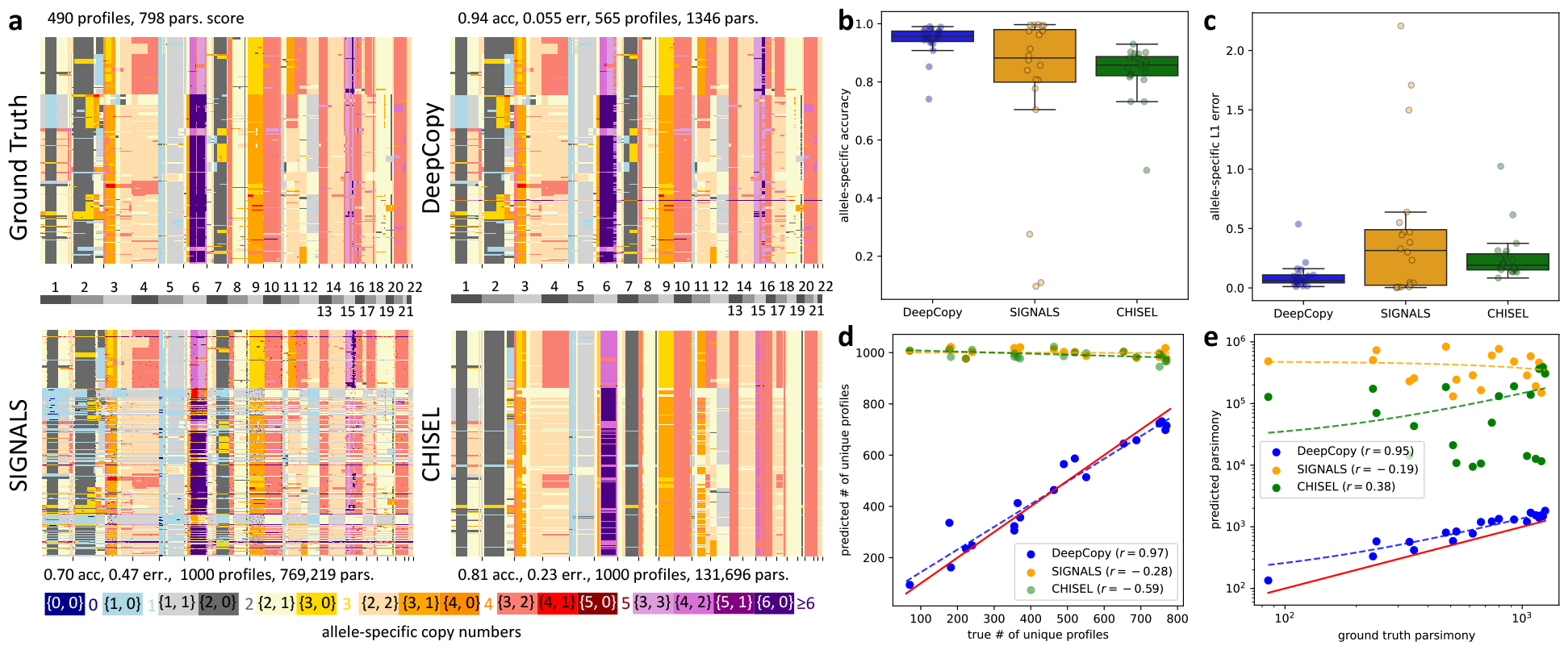
Results on simulated data. **a**, Allele-specific copy number profiles from the ground truth as well as predictions by DeepCopy, SIGNALS and CHISEL for a single simulation instance with a whole genome duplication. **b**, Accuracy values of unordered allele-specific copy numbers. **c**, L1 errors of unordered allele-specific copy numbers. **d**, The number of predicted unique copy number profiles (*y*-axis) vs. ground-truth number of unique profiles (*x*-axis), with the *y* = *x* line in red and Pearson correlations indicated in the key and dashed lines. A small amount of jitter was added to predictions with exactly 1,000 unique copy number profiles for clarity. **e**, Parsimony scores for phylogenetic trees inferred from ground-truth (*x*-axis) and predicted copy-number profiles (*y*-axis), with the ground-truth *y* = *x* line shown in red, and best-fit lines shown with dashes as well as Pearson correlations indicated in the key. The best-fit lines can appear slightly curved as a consequence of the plot’s log scale.

Third, we compared the number of *unique copy number profiles* inferred by each method to that in the ground truth. The 1000 cells of the simulation instance shown in Fig. 2a correspond to 490 unique copy number profiles in the ground truth. While DeepCopy slightly overfit the data with 565 inferred unique copy number profiles, both SIGNALS and CHISEL inferred a unique copy number profile for each cell (Fig. 2a). Looking across all simulation instances (Fig. 2d), we find that DeepCopy better matched the true number of unique profiles with a Pearson correlation of *r* = 0.98 and a median percentage error of 6.92% compared to SIGNALS (*r* = *−*0.28 and 110%) and CHISEL (*r* = *−*0.59 and 110%). Fourth, we utilized the copy number profiles inferred by each method to construct a phylogenetic tree as described in Section 4.4, defining the *parsimony score* as the number of CNA events on this tree using the zero-agnostic copy number transformation (ZCNT) distance [34]. For the shown simulation instance in Fig. 2a DeepCopy predicted a far more parsimonious solution (with parsimony score 1,346) than SIGNALS (769,219) or CHISEL (131,696), despite being less parsimonious than the ground truth (798). Note that high parsimony values may be reflective of homoplasy, i.e., identical CNAs occurring independently at many locations on the tree, due to a lack of evolutionary structure in the predictions even if the predictions do not look very visually noisy as is the case for CHISEL’s prediction. We find similar trends across all 20 simulation instances (Fig. 2e), with DeepCopy’s parsimony scores both lower and better correlated with the ground-truth values (Pearson correlation *r* = 0.95 and median value 1201) compared to SIGNALS (*r* = *−*0.19 and median value 348,770) and CHISEL (*r* = 0.38 and median value 59,560). However, the median ground truth parsimony of 708 is even lower than DeepCopy’s parsimony, showing there still exists room for improvement.

In summary, DeepCopy infers more accurate copy-number profiles that better recapitulate ground-truth clonal structure and lead to more parsimonious phylogenetic trees compared to SIGNALS and CHISEL. These findings are not unexpected, as unlike DeepCopy, neither SIGNALS nor CHISEL uses an evolutionary model to constrain inferred copy number profiles.

### 2.3 Evaluation on an ovarian cancer dataset sequenced with DLP+ technology

Next, we benchmarked DeepCopy on 890 direct library preparation (DLP+) sequenced cells from three clonally-related cancer cell lines sourced from the same high-grade serous ovarian cancer OV2295 [20], with coverages ranging from 0.1*×* to 0.4*×* (Fig. S4). We compared DeepCopy to SIGNALS, CHISEL, and Alleloscope. As Alleloscope and SIGNALS applied additional filtering criteria, we restricted our analysis to the *n* = 617 cells for which all methods were run and produced predictions.

We find that the difference in predicted copy number profiles between Alleloscope and all other methods is very stark (L1 distances at least 1.35 compared to a distance of 0.14 between DeepCopy and SIGNALS shown in Fig. S5), as a result of Alleloscope predicting no WGD while all other methods predicted WGDs on many cells (Fig. 3a). For the other methods, the differences are more subtle with the zoomed in plots of chromosome 1 showing a lower level of noise in DeepCopy and that DeepCopy’s and SIGNAL’s predictions are most similar, while CHISEL’s predictions have more substantial differences, especially towards telomeres of chromosomes (Fig. 3b). Looking at all *n* = 617 cells and the complete genome, we find that SIGNALS predicted a unique copy number profile for each cell, while the largest clones for DeepCopy, CHISEL, and Alleloscope contained 41, 9, and 2 cells, respectively (Fig. 3c). In total, DeepCopy, CHISEL, and Alleloscope predicted 468, 469, and 610 unique profiles, respectively. Our findings of extensive heterogeneity in CNA profiles are in line with previous analyses of high-grade serous ovarian cancer [35–38].

**Figure 3.**
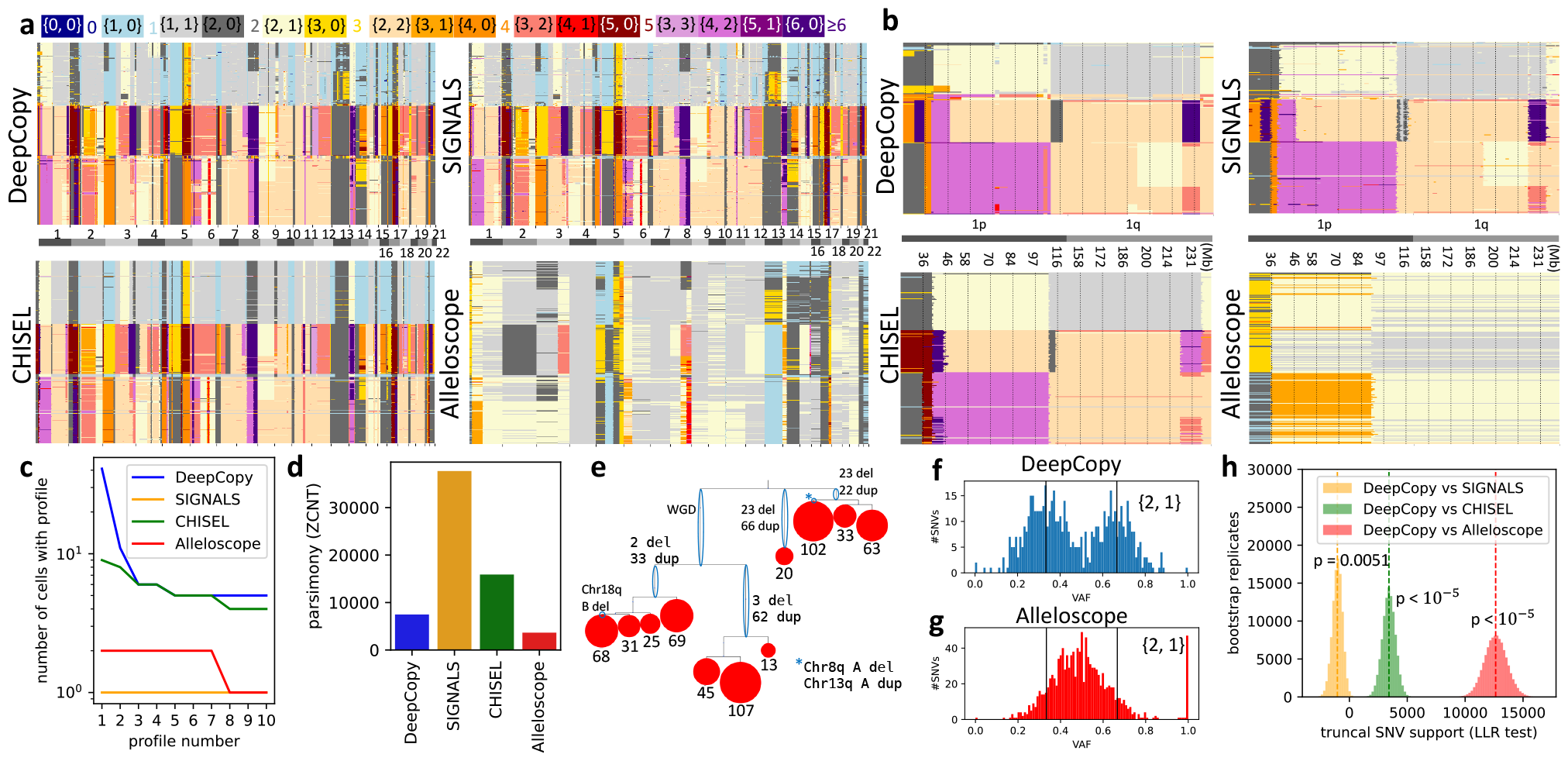
Results on high-grade serous ovarian cancer OV2295 [20]. **a**, Allele-specific copy number profiles for DeepCopy, SIGNALS, CHISEL, and Alleloscope on *n* = 617 cells. Unlike other methods, Alleloscope did not predict WGD for any cell. **b**, Allele-specific copy number profiles of chromosome 1, showing that while DeepCopy retains the ability to detect small CNAs also detected by SIGNALS, the other methods’ predictions are far noisier. **c**, The size of the 10 largest clones for each method, showing larger clones for DeepCopy and CHISEL than SIGNALS and Alleloscope. **d**, DeepCopy’s predictions resulted in a more parsimonious phylogeny than SIGNALS or CHISEL, while Alleloscope resulted in the lowest parsimony score. **e**, A simplified drawing of DeepCopy’s tree with edges circled in blue labeled by CNAs (edges with many CNAs are labeled by the number of duplications/deletions), and leaves indicate clades labeled by the number of comprising cells. **f-g**, VAF values on the copy number 1, 2 for DeepCopy and Alleloscope. For DeepCopy but not Alleloscope, VAFs are concentrated around 1*/*3 and 2*/*3 (indicated with black vertical lines) as expected. **h**, Log likelihood ratios (LLR) of truncal SNV support on bootstrapped replicates between our method and the alternative methods SIGNALS, CHISEL, and Alleloscope. DeepCopy outperforms CHISEL (LLR 3,418.66 and *p <* 10^*−*5^) and especially Alleloscope (LLR 12,635.36 and *p <* 10^*−*5^) while being slightly outperformed by SIGNALS (LLR *−*1,033.42 and *p* = 0.0051).

To assess the downstream effects of overfitting to noise, we constructed phylogenies from each method’s copy number profiles and evaluated the resulting parsimony scores via a similar procedure described in Section 2.2. We find that DeepCopy’s tree had a parsimony score of 7,878 (Fig. 4e), which is far more parsimonious than the trees produced from SIGNALS’ and CHISEL’s copy number profiles, achieving parsimony scores of 37,692 and 15,890, respectively. This difference in parsimony is visually clear in images of the full trees shown in Fig. S6, showing long branches near the leaves indicative of many CNAs occurring on individual cells rather than on larger clones. We note that Alleloscope achieved an even lower parsimony score of 3,651, however, we believe this is due to Alleloscope incorrectly predicting a lack of WGD. To test this hypothesis and orthogonally validate our DeepCopy’s predictions, we evaluated consistency with orthogonal SNV data reported by Laks et al. [20]

**Figure 4.**
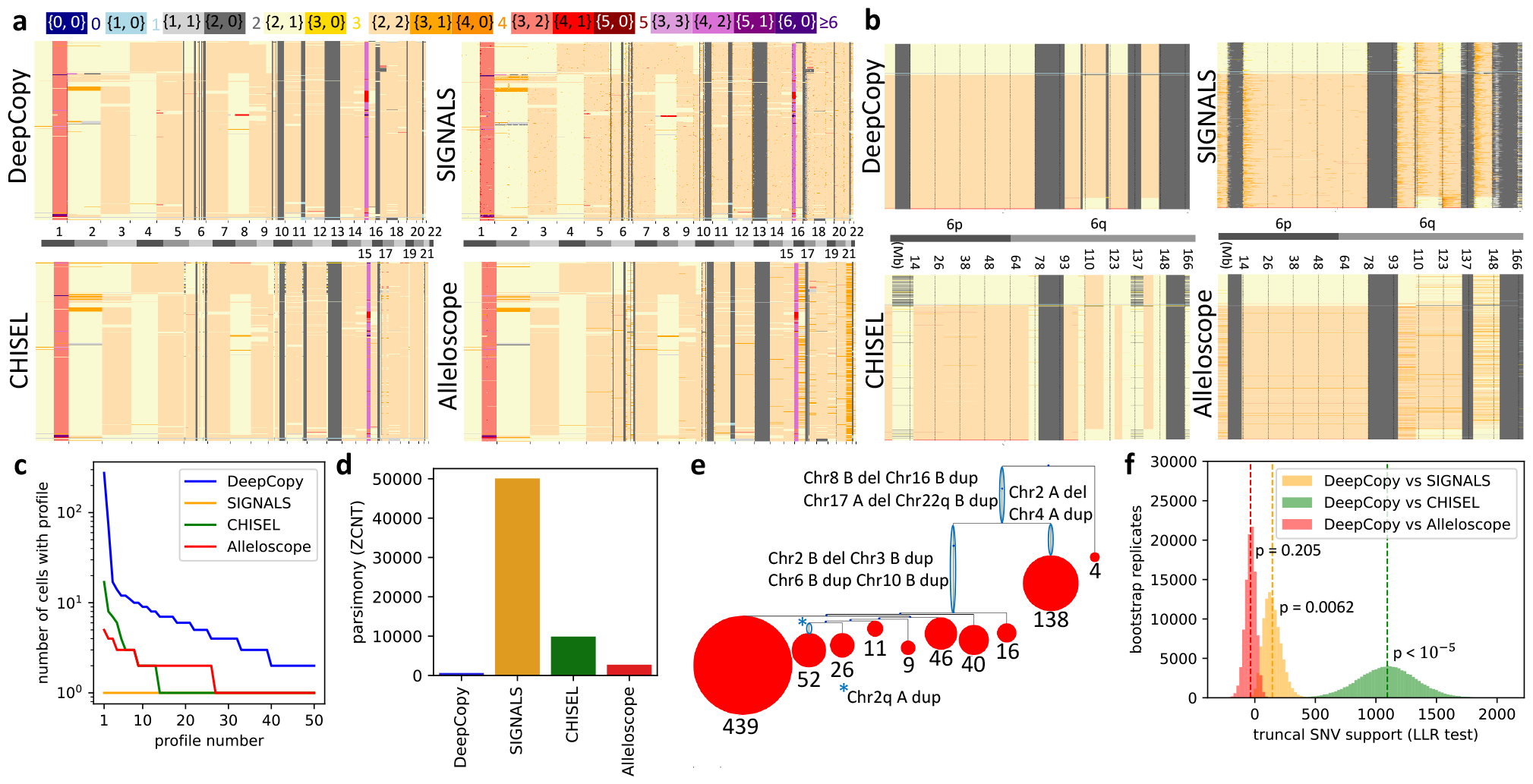
Results on breast cancer patient S0. **a**, Allele-specific copy number profiles for DeepCopy, SIGNALS, CHISEL, and Alleloscope on *n* = 785 cells. **b**, Allele-specific copy number profiles of chromosome 6, showing that while DeepCopy retains the ability to detect small CNAs also detected by SIGNALS, the other methods’ predictions are far noisier. **c**, The size of the 50 largest clones for each method, showing much larger clones for DeepCopy than SIGNALS, CHISEL SIGNALS, and Alleloscope. **d**, DeepCopy’s predictions resulted in a tree with a far lower parsimony score (630) than SIGNALS (50,085), CHISEL (9,877), or Alleloscope (2,733). **e**, A simplified drawing of DeepCopy’s tree with edges circled in blue labeled by CNAs (edges with many CNAs are labeled by the number of duplications/deletions), and leaves indicate clades labeled by the number of comprising cells. **f**, Log likelihood ratios (LLR) of SNVs support on bootstrapped replicates between our method and the alternative methods SIGNALS, CHISEL, and Alleloscope. DeepCopy outperforms CHISEL (LLR 1,093.83 and *p <* 10^*−*5^) and somewhat outper-forms SIGNALS (LLR 145.26 and *p* = 0.0062) while being slightly outperformed by Alleloscope (LLR *−*35.24 and *p* = 0.205).

Specifically, we focused on a subset of 2,801 out of 14,068 SNVs that are likely *truncal*, i.e., present in the most recent common ancestor of all tumor cells (details provided in Section 4.4). Truncal SNVs that occur on a segment with copy number {*X*^*A*^, *X*^*B*^} must have a variant allele frequency (VAF) on those cells equal to some integer multiple of 1*/*(*X*^*A*^ +*X*^*B*^). We find that this is the case for allele-specific copy number {2, 1} for DeepCopy (Fig. 3f) as well as SIGNALS and CHISEL (Fig. S7), showing bimodal distributions with peaks around 1*/*3 and 2*/*3, but not Alleloscope, showing a unimodal distribution with a single peak around 1*/*2 (Fig. 3g). Moreover, allelic mirroring for the CNA {2, 1} (demonstrated in Fig. S8) is predicted by all methods and reflected in the VAFs of truncal SNVs for DeepCopy, SIGNALS, CHISEL but not Alleloscope, finding that SNVs with a VAF near 1*/*3 for (2, 1) have a VAF near 2*/*3 for (1, 2) and vice versa (Fig. S9). We see similar trends for other CNAs (Fig. S7 and Fig. S10), finding several cases where SNVs show evidence for LOH whereas Alleloscope inferred heterozygous CNAs. To better quantify differences between methods, we developed a statistical test for goodness of fit with the SNV data, exploiting the fact that the vast majority of truncal SNVs seem to precede their coinciding CNAs (Supplementary Section B.1). In such cases, we expect a VAF of either *X*^*B*^*/*(*X*^*A*^ + *X*^*B*^) or *X*^*A*^*/*(*X*^*A*^ + *X*^*B*^) depending on which allele the SNV occurred. We defined a log-likelihood ratio (LLR), measuring the relative probability of each truncal SNV’s observed reference and variant reads given the copy numbers predicted by DeepCopy when compared to each alternative method, with positive values supporting DeepCopy’s predictions and negative values supporting the method being compared against (Supplementary Section A.4.3). Alleloscope has by far the worst statistical support of truncal SNVs (LLR: 12,635.36, *p <* 10^*−*5^), followed by CHISEL (LLR: 3,418.66, *p <* 10^*−*5^) whereas SIGNALS achieved slightly better SNV support than DeepCopy (LLR: *−*1,033.42, *p* = 0.0051). The comparatively small log-likelihood ratio between SIGNALS and DeepCopy is reflective of general agreement.

In summary, we find that while Alleloscope produced the most parsimonious solution, it did not properly account for the presence of a subclonal WGD event. On the other hand, similarly to our simulation results, we find that DeepCopy avoids spurious CNAs producing a more parsimonious solution than SIGNALS or CHISEL while retaining a high accuracy towards legitimate CNAs as reflected by accurately fitting the orthogonal truncal SNV data.

### 2.4 Evaluation on a breast cancer dataset sequenced with 10x Chromium CNV technology

For further benchmarking, we considered single-cell DNA sequencing data from breast cancer patient S0 sequenced with Chromium Single Cell Copy Number Variation (CNV) Solution from 10x Genomics. A total of 10,202 cells, distributed across five spatial sections, were sequenced with coverage ranging from 0.01*×* to 0.05*×* (Fig. S11). While both CHISEL and DeepCopy produced output for all cells, the SIGNALS output consisted of only 3,540 cells due to additional filtering criteria, including removal of normal cells as well as high noise cells suspected to be actively replicating, doublets, or otherwise problematic. Additionally, we utilized publicly available predictions from Alleloscope on cells from one particular section of the tumor, amounting to a total of *n* = 785 cells for which all methods produced predictions.

In contrast to the DLP+ data, we find a general agreement between DeepCopy, SIGNALS, and Alleloscope other than additional small CNAs in SIGNALS predictions likely due to noise (Fig. 4a and Fig. S12) and CHISEL predicting a smaller quantity of LOH, as can be seen in chromosome 6 (Fig. 4b). We find that DeepCopy identified large sets of cells with identical copy number profiles (see Supplementary Section A.4.1 for details) — the largest such set identified by DeepCopy consists of 276 cells (Fig. 4c). On the other hand, the largest set of cells with identical profiles identified by SIGNALS, CHISEL, and Alleloscope consist of only 1, 17, and 5 cells, respectively. Additionally, DeepCopy determined fewer unique copy number profiles (206) compared to SIGNALS (785), CHISEL (737) and Alleloscope (747). This lower number is due to DeepCopy’s use of a shared evolutionary model; indeed, NaiveCopy, which has the same data processing pipeline and statistic model for the data as DeepCopy but lacks an evolutionary model, produced 628 unique copy number profiles (Fig. S13). Similar comparisons on the set of 3,540 cells for which DeepCopy, SIGNALS, and CHISEL were run are provided in Fig. S14 and Fig. S15.

Additionally, we calculated phylogenies from the copy number profiles and evaluated their parsimony scores. We find that DeepCopy’s phylogeny is far more parsimonious (score: 582) than SIGNALS (50,085), CHISEL (9,877), and Alleloscope (2,733). This difference in parsimony is visually clear in images of the full trees shown in Fig. S16, showing long branches near the leaves indicative of many CNAs occurring on individual cells rather than on larger clones. DeepCopy’s tree is also simplified to nodes representing major clades as illustrated in Fig. 4e, showing the presence of CNAs common to large clades. To assess whether DeepCopy’s increased robustness to spurious CNAs came at the expense of substantially decreased sensitivity to legitimate CNAs, we validated on orthogonal SNV data as described in Section 2.3. Specifically, we analyzed a subset of 755 likely truncal SNVs, finding that DeepCopy provided a somewhat better fit than SIGNALS (LLR 145.26 and *p* = 0.0062), a substantially better fit than CHISEL (LLR 1,093.83 and *p <* 10^*−*5^), and a slightly worse but not significant fit than Alleloscope (LLR *−*35.24 and *p* = 0.205). VAF values are plotted in Fig. S17, showing only minor differences between methods. Additionally in Fig. S18, we show that DeepCopy, SIGNALS, and CHISEL predict allelic mirroring on copy number {2, 1}, but this prediction is supported by orthogonal SNV data only for DeepCopy and SIGNALS, finding SNVs with a VAF near 1*/*3 on copy number (2, 1) have a VAF near 2*/*3 on copy number (1, 2) and vice versa.

In summary, we have shown that DeepCopy avoids spurious CNAs by producing a far more parsimonious solution with fewer unique copy number profiles than SIGNALS, CHISEL, and Alleloscope, which is in line with the clonal structure commonly observed in breast cancer that results from punctuated evolution [39].

### 2.5 Consistency of DeepCopy on matched 10x and ACT sequencing samples

In our final analysis, we assessed whether DeepCopy is able to produce consistent predictions across multiple sequencing technologies applied to the same tumors. To that end, we analyzed breast cancer patient TN3, which was sequenced using both ACT [22] and 10x sequencing technologies. The coverage for the *n* = 1,101 cells sequenced with ACT technology ranged from 0.005*×* to 0.015*×*, whereas the coverage for the *n* = 1,070 cells sequenced with 10x technology ranged from 0.01*×* to 0.07*×* (Fig. S19). DeepCopy inferred 119 unique copy number profiles for the ACT cells and 90 profiles for the 10x cells. Strikingly, when projecting the profiles to two dimensions using UMAP and clustering using DBSCAN, we find that the cells cluster into four common groups (Fig. 5a). Moreover, each cluster has a similar proportion of cells for the two technologies (Fig. 5b), with 448 (41%), 451 (41%), 67 (6%), and 135 (12%) cells respectively for the four clusters for ACT sequenced cells and 434 (40%), 404 (38%), 85 (8%), and 147 (14%) cells respectively for 10x sequenced cells. Phylogenies for both sequencing technologies with cells colored by cluster are shown in Fig. S20, demonstrating general agreement. Copy number profiles for each cluster for the two sequenced technologies are shown in Fig. 5c, showing the similarity between the two technologies for each cluster. Thus, we find that DeepCopy successfully uncovers similar ground truth copy number profiles with similar proportions despite the use of two different sequencing technologies.

**Figure 5.**
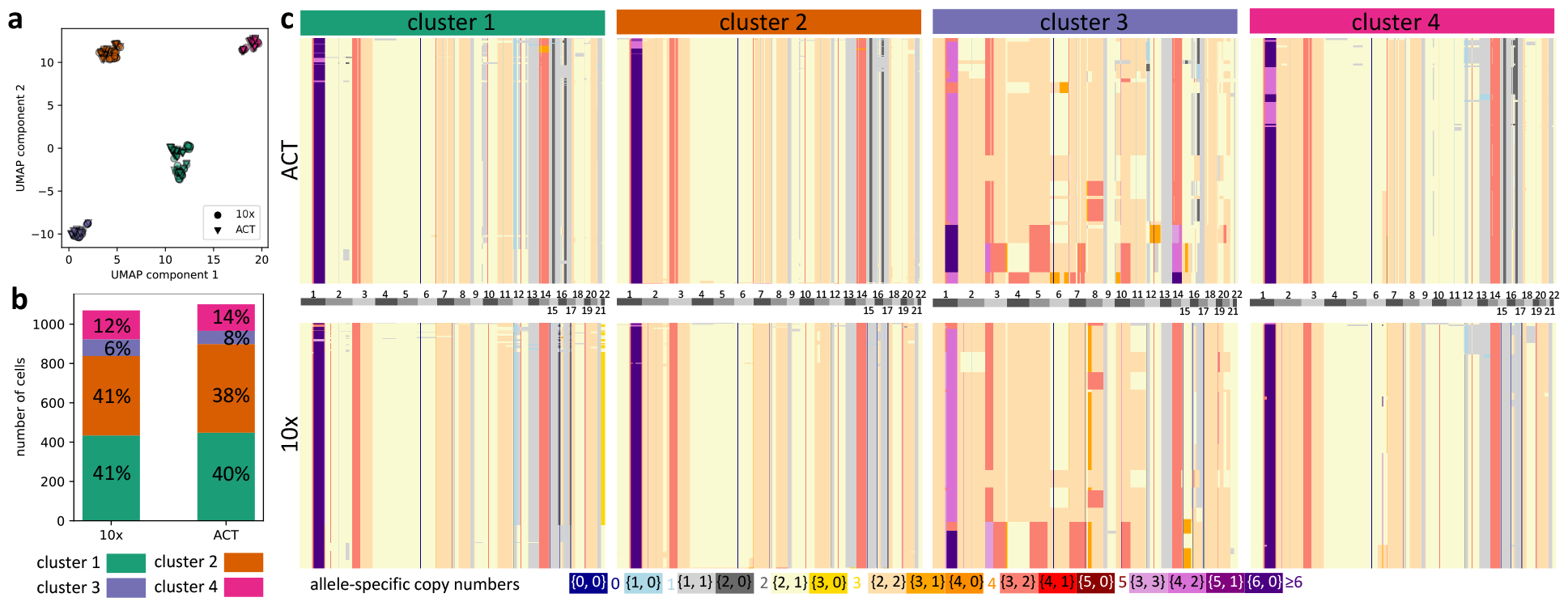
Analysis of breast cancer patient TN3 [22] for both ACT and 10x Chromium technologies. **a**, A UMAP of copy number profiles shows 4 clear clusters on pooled copy number profiles for both technologies. **b**, Similar proportions for each cluster exist for both technologies. For ACT sequenced cells there are 448, 451, 67, and 135 cells respectively in the four clusters, meanwhile for 10x sequenced cells there are 434, 404, 85, and 147 cells respectively. **c**, Copy number profiles in all four clusters are shown for both technologies (with cells in each cluster sorted by average ploidy).

## 3 Discussion

We introduced the method DeepCopy, a deep reinforcement learning based method for haplotype-specific copy number calling. This method utilizes a generative model of CNA evolution to model the probability of different copy number profiles. We note that our use of reinforcement learning for modeling a generative evolutionary process is conceptually similar to our previous work CloMu [40], an algorithm for modeling SNV evolutionary trees. While CloMu directly models evolutionary tree probabilities, DeepCopy models the probabilities of trajectories starting on the normal cell and ending on some copy number profile, allowing for a more flexible search while avoiding local minima. On simulated data, DeepCopy predicts copy number profiles and numbers of unique copy number profiles that better fit ground truth values than SIGNALS or CHISEL. On real data of breast and ovarian cancer, DeepCopy produced copy number profiles that fit more parsimonious trees and had more realistic clone sizes than existing methods. Additionally, we validated DeepCopy on orthogonal SNV data, showing a better fit than CHISEL on both datasets, a much better fit than Alleloscope on ovarian cancer, and similar fits as SIGNALS overall. Finally, we found that DeepCopy inferred identical clonal structure of a breast cancer patient when run separately on the cells sequenced with 10x and ACT technologies.

Currently, DeepCopy’s optimization is guided by copy number profiles predicted by NaiveCopy. Consequently, another useful addition to DeepCopy would be allowing any copy number profiles to be provided for guiding DeepCopy, such as the copy number profiles predicted by SIGNALS. Although DeepCopy’s predictions yield more parsimonious trees than competitor methods, simulations show these parsimony values are still meaningfully higher than the ground truth. Consequently, further improving DeepCopy in order to produce a maximally parsimonious tree is an important direction for future work. One approach for this could be a post-processing local search to modify DeepCopy’s CNA tree and obtain improved copy number profiles. For instance, an MCMC optimization approach on CNA trees as performed in SCICoNE [26] could be applied. Additionally, our simulations only generate read depth and haplotype-specific read depth data but not BAM files. A clear direction for future work is generating simulated BAM files in order to enable comparisons against methods that do not utilize out intermediate inputs such as Alleloscope. One other natural extension of DeepCopy would be including the sex chromosomes currently excluded from our analysis. Future work could also include the ability to detect doublets and S-phase cells. Currently, DeepCopy produces copy number profiles for such cells, but they are not specifically identified as doublets or S-phase. We also plan on updating our visualization tool [41] to support the interpretation of DeepCopy results. Utilizing ideas used in the orthogonal validation of CNAs using SNVs [42], we plan on performing integrative inference of SNVs and CNAs, improving upon previous work [43]. Finally, a broader more complex future direction would be allowing DeepCopy to share a subset of parameters across patients and analyzing a large cohort of patients. One essential advantage of deep learning is its ability to learn complex subtle patterns across large datasets. Consequently, we believe the advantages of DeepCopy’s deep reinforcement learning could be greatly extended if training parameters across a large cohort of patients were enabled. Such a system could potentially learn general patterns of CNA evolution while also learning the particular evolutionary trajectories of individual patients, akin to methods for identifying evolutionary trajectories of SNVs [40, 44–48].

## 4 Methods

### 4.1 Problem statement

Given a set of aligned sequencing reads from single-cell DNA sequencing data, we wish to identify a copy number profile *P* = [*P* ^(1)^, *P* ^(2)^]^*T*^ ∈ ℕ^2*×L*^ for each cell across *L* genomic regions. More precisely, the copy numbers that we seek are both allele specific and haplotype specific. That is, each copy number 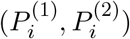 indicates the number of copies of genetic region *i* of each of the two parental alleles. Additionally, the copy numbers are phased across each chromosome such that copy numbers 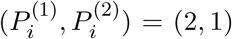 and 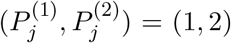 of two different positions *i* and *j* on the same chromosome indicate two distinct copy numbers in terms of whether the maternal or paternal haplotype is the one amplified.

Although the raw input is a set of aligned sequencing reads for each cell, this needs to be further processed to produce the appropriate inputs for copy-number calling. Specifically, we must split the genome into *L* genomic regions, or *segments* such that the copy number is constant in each segment. Then, we determine read counts for each cell and segment, which are corrected for several biases, to produce read depths **R** = [*R*_1_, …, *R*_*N*_ ]^*T*^ ∈ *ℝ*^*N ×L*^ such that the read depth of a segment of a cell is proportional to the total copy number, i.e., 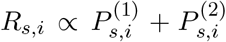 Additionally, SNPs must be detected, counted, phased into haplotype blocks, and phased across haplotype blocks in order to produce the number of reads matched to each haplotype for each segment. This then produces the B-allele frequencies (BAFs) **B** = [*B*_1_, …, *B*_*N*_ ]^*T*^ ∈ [0, 1]^*N×L*^, which measure the proportion of reads coming from the second haplotype (typically set to be the minor haplotype), i.e., 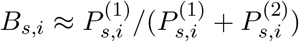. We provide more details on determining the *L* segments and inputs **R** and **B** in Supplementary Section A.1.3.

Our measurements **R** and **B** not only depend on the copy number but include additional uncertainty specific to the cell and segment being considered. For instance, GC bias as well the number of SNPs in a segment can affect the variance in read depth and BAF, respectively, across segments (Fig. S1). Additionally, differing coverage across cells can affect the variance across cells. Consequently, we define 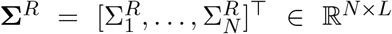 as the cell and segment-specific variances for the read depths **R** = [*R*_1_, …, *R*_*n*_]^*T*^, and 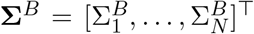 as the cell and segment-specific variances for the BAFs **B** = [*B*_1_, …, *B*_*N*_ ]^*T*^. In Supplementary Section A.1.3, we describe how we estimate these values. Given these mean and variance values, we define 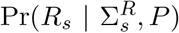 as the probability of the observed read depths *R*_*s*_ given variances 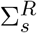 and copy number profile *P* of a cell *s*. Similarly, we define 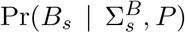 as the probability of the observed BAFs *B*_*s*_ given the standard deviations 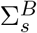 and the copy number profile *P*. Precise definitions of the probability distributions used are given in Supplementary Section A.2.1. This gives the below equation for the observed data given a copy number profile *P* .

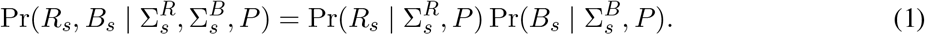

While previous works infer a copy number profile *P*_*s*_ of each cell *s* in isolation, here we specifically account for the joint evolution of all copy number profiles **P** = [*P*_1_, …, *P*_*N*_ ]. As such, we define Pr(*P* | *θ*) as the probability of the copy number profile *P* according to a model with tumor-specific parameters *θ*. For our method DeepCopy, we utilize a deep neural network in combination with an evolutionary model to calculate Pr(*P* | *θ*). We define *𝒫* as the set of all possible haplotype-specific copy number profiles with *L* segments. In practice, *𝒫* will be bounded by setting a maximum per-haplotype copy number of *C*_max_ (set to a default value of 19). The probability of observed read depths *R*_*s*_ and BAFs *B*_*s*_ is

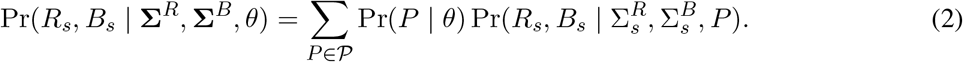

The probability of the read depths **R** = [*R*_1_, …, *R*_*N*_ ]^*T*^ and the BAFs **B** = [*B*_1_, …, *B*_*N*_ ]^*T*^ for *N* cells is then

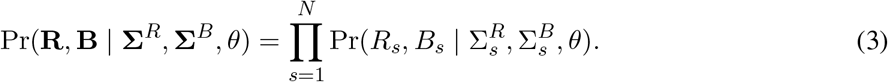

This leads to the following problem.

#### Problem 1

(Copy Number Profile Distribution). *Given read depths* **R** = [*R*_1_, …, *R*_*N*_ ]^*T*^ *and B-allele frequencies* **B** = [*B*_1_, …, *B*_*N*_ ]^*T*^ *and their variances* **Σ**^*R*^ *and* **Σ**^*B*^ *for N cells and L segments find the model parameters θ that maximize* Pr(**R, B** | **Σ**^*R*^, **Σ**^*B*^, *θ*).

Once we identify the parameters *θ* that maximize the probability of the observed measurement data, it is relatively straightforward to find a predicted profile *P*_*s*_ for each cell *s* such that

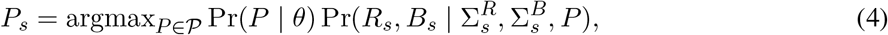

provided one can sample copy number profiles *P* from Pr(*·* | *θ*).

### 4.2 Data processing and NaiveCopy

The input to NaiveCopy’s pipeline is a BAM file with read groups for each sequenced cell of a tumor. This is then processed into total read counts as well as haplotype-specific counts for *K* bins of size *l*_bin_ in the genome (in practice we set *l*_bin_ = 100kb). The processing of calculating the set of bins and their GC-bias corrected read counts 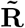 is described in Supplementary Section A.1.1. The process of calculating haplotype-specific read counts **A** is described in Supplementary Section A.1.2 where SNPs are found using bcftools [49], haplotype blocks are determined using SHAPE-IT [50], and haplotype blocks are phased using our phasing algorithm. The genome is then split into *L* segments consisting of many 100kb bins, as described in Supplementary Section A.1.3. In each of these segments, the average read depth **R** is calculated as well as the B-allele frequency **B** from the haplotype-specific counts **A**. Additionally, the variance **Σ**^*R*^ in the read depth and the variance **Σ**^*B*^ in the BAF are estimated using a heuristic described in Supplementary Section A.1.3.

NaiveCopy produces copy number profiles 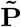 by maximizing 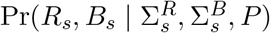 (defined in Supplementary Section A.2.1). This is done by first determining a cell-specific scaling factor (Supplementary Section A.2.2), followed by the determination of integer copy numbers (Supplementary Section A.2.3). Note that NaiveCopy solves Problem 1 by taking a flat prior for Pr(*P* | *θ*) (with the exception of restricting cell specific scaling factor). The set 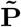 of copy number profiles returned by NaiveCopy is used to help guide DeepCopy’s optimization. Fig. 6a-b shows a diagram of this pipeline.

**Figure 6.**
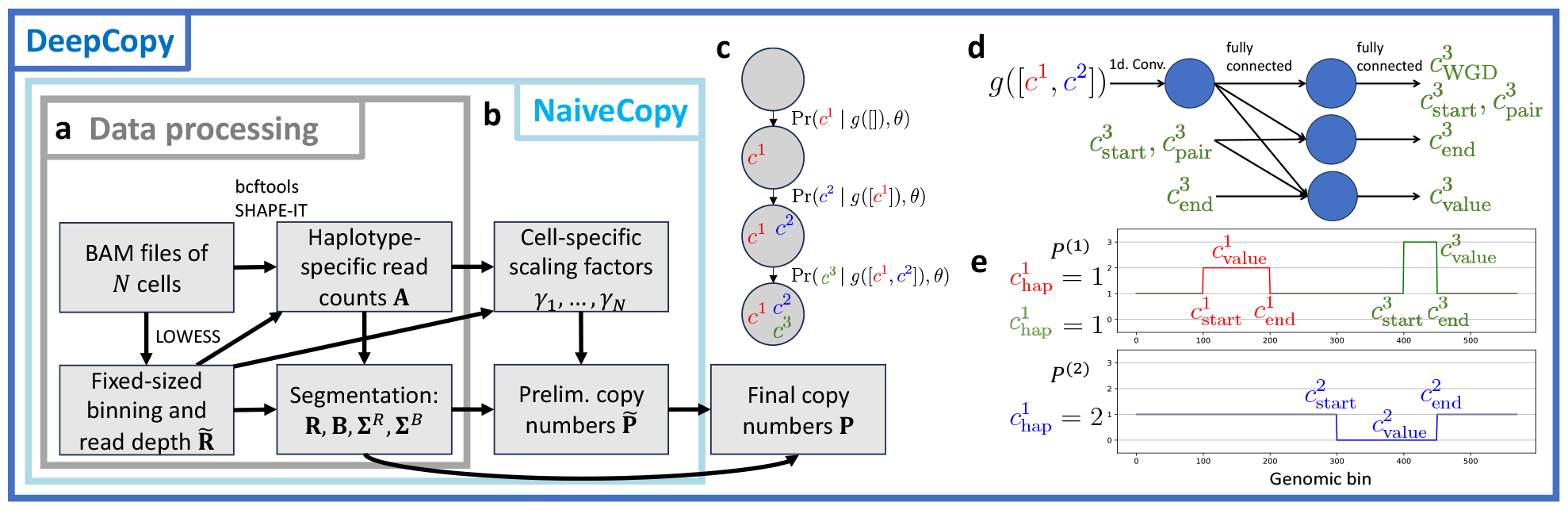
Methods overview. **a**, Data processing steps to obtain measurements **R, B, Σ**^*R*^, **Σ**^*B*^ for *L* segments from BAM files of *N* cells. **b**, Given these measurements, NaiveCopy produces preliminary copy number profiles 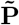 by identifying a cell-specific scaling factor *γ*_*s*_ for each cell *s*, without using an evolutionary model. **c**, In DeepCopy’s evolutionary model, a copy number profile is generated by the application of a series of CNAs such that the probability of each new CNA depends on the previous copy number profile. **d**, These probabilities are determined by a neural network. Here, *g*([*c*^1^, *c*^2^]) is the copy number profile generated by applying the two CNA tuples *c*^1^ and *c*^2^ to the normal cell. Then, the probability of the components of CNA tuple *c*^3^ is generated. Ultimately, Pr(*c*^3^ | *g*([*c*^1^, *c*^2^]), *θ*) is the probability of the next CNA *c*^3^ given that CNAs *c*^1^ and *c*^2^ have already been applied. **e**, A potential copy number profile *g*([*c*^1^, *c*^2^, *c*^3^]) generated by panel c is shown. Specifically, the copy numbers for haplotype 1 and haplotype 2 are plotted, demonstrating the CNAs *c*^1^, *c*^2^, and *c*^3^. For each CNA tuple, the haplotype number, the start position, the end position, and the value are shown in the copy number profile.

### 4.3 DeepCopy

#### 4.3.1 Evolutionary model

To solve the Copy Number Profile Distribution problem (Problem 1), we define an evolutionary model for probabilistically generating copy number profiles. Each cancer is trained on independently, and so trained model parameters *θ* describe the evolution of a single cancer and capture the probability of different copy number profiles within that tumor. The copy number profile of a normal cell is 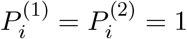 for all segments *i* ∈ [*L*] = {1, …, *L*}. In our model, a tumor first starts with a single clone with the copy number profile of a normal cell. Then, a CNA occurs producing a new clone. The new clone has a copy number profile in which the values in either *P* ^(1)^ or *P* ^(2)^ are modified to new values for some interval of segments within a chromosome. Alternatively, a whole genome duplication can occur in which all copy numbers are doubled. In either case, this produces a new copy number profile which can then be modified by additional CNAs. This process repeats unit terminating on some final CNA. As an example, if segments 30 through 60 are within some chromosome, a CNA could be amplifying *P* ^(1)^ in segments 35 to 50 to the value 2, i.e.,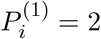 for segments 35 *≤ i ≤* 50.

To model the evolution of CNAs, we define a *CNA tuple c* = (*c*_WGD_, *c*_hap_, *c*_start_, *c*_end_, *c*_value_). The value of *c*_WGD_ is 1 if the mutation is a whole genome duplication and 0 otherwise. Assuming *c*_WGD_ = 0, then *c*_hap_ ∈ {1, 2} is the haplotype number of the CNA, *c*_start_ ∈ [*L*] is the starting segment, *c*_end_ ∈ [*L*] is the ending segment, which must be on the same chromosome as the starting segment, and *c*_value_ ∈ *ℤ* indicates the change in the copy number. Let *C* be the set of all such possible CNA tuples (with the restriction *−*5 *≤ c*_value_ *≤* 5). We define a *generating sequence G* as any list of CNA tuples [*c*^1^, …, *c*^*k*^] ⊆ 𝒞. We define *g*(*G*) as the copy number profile generated by applying the CNAs in *G* to the copy number profile of a normal clone. An example of a copy number profile being generated by a sequence of CNA tuples is shown in Fig. 6c-e. A more mathematically precise definition of *g* is given in Supplementary Section A.3.1.

Given an existing copy number profile, a model can assign a probability to each new CNA tuple as well as a probability that no new CNAs will occur. We define this probability for a new CNA tuple as Pr(*c* | *P, θ*) where *c* ∈ *𝒞* is a possible CNA tuple. Additionally, we define Pr(stop | *P, θ*) as the probability that no new CNAs will occur in the cell. In practice, since 𝒞 can be extremely large, we first assign a probability to each starting position *c*_start_ given the existing profile *P*, then assign a probability to each ending position *c*_end_ given the starting position *c*_start_ and *P*, and finally assign a probability to the copy number value *c*_value_ given the start position *c*_start_, ending position *c*_end_, and *P*. This is done using a deep convolutional neural network with an architecture shown in Fig. 6c, and described in Supplementary Section A.3.2. Using this, we assign a probability to any generating sequence *G* = [*c*^1^, …, *c*^*k*^] of CNA tuples as

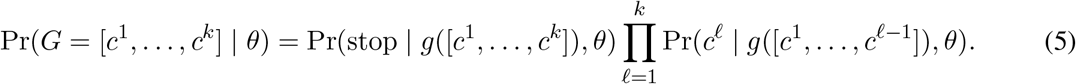

In the above equation *g* applied to the empty sequence, i.e., *g*([]), results in the normal copy number profile. The process of forming a generating sequence by repeatedly applying CNAs is shown in Fig. 6b. Putting together our definitions, we define the probability for any copy number profile *P* as

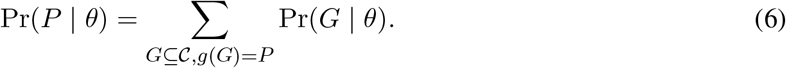

Note that any *G* ⊆ 𝒞 is a generating sequence. An example generated copy number profile is shown in Fig. 6e. Putting this probability into equation (2) then allows us to solve Problem 1 by optimizing CNA tuple probabilities.

#### 4.3.2 Copy number calling via reinforcement learning

Before describing our approach, we describe a simplified version that could work in theory but has difficulties in practice. To exactly solve Problem 1, we must maximize Pr(**R, B** | **Σ**^*R*^, **Σ**^*B*^, *θ*), which requires calculating a sum over all *P* ∈*𝒫*. The set 𝒫of all possible copy number profiles is too large for this to be feasible, so instead we must use sampling. We maximize log Pr(**R, B** | **Σ**^*R*^, **Σ**^*B*^, *θ*), which is equivalent to maximizing Pr(**R, B** | **Σ**^*R*^, **Σ**^*B*^, *θ*) while allowing for useful mathematical manipulations. To achieve this maximization, we calculate the gradient of log Pr(**R, B** | **Σ**^*R*^, **Σ**^*B*^, *θ*) with respect to *θ*. As derived in Supplementary Section A.3.3, we have the equation

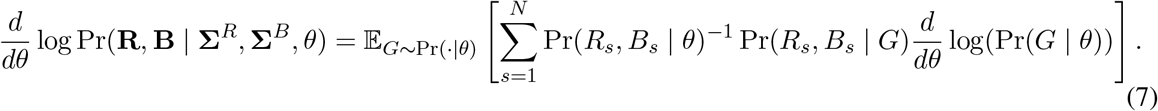

In the above equation and in the following text, we use the shorthands 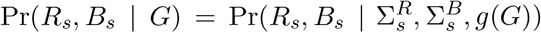 and 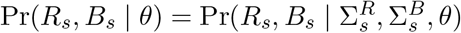. We define the reward function as

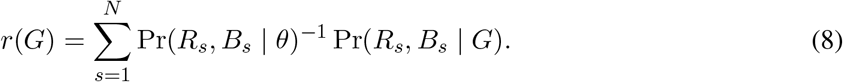

This reward function requires an estimate of Pr(*R*_*s*_, *B*_*s*_ | *θ*), which can be accomplished via sampling as described in Supplementary Section A.3.4. With this reward function, we have a standard policy learning gradient [51] (assuming a reward is only given at the last time step)

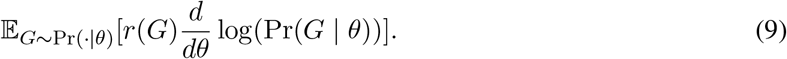

Although this all could work in theory, solving this problem from scratch requires significant computational resources and may require substantial tuning of hyperparameters on each new dataset in order to ensure successful optimization. This would be acceptable if the goal of this paper was to successfully estimate a copy number profile for a single cancer. However, the goal is instead to provide a robust and fast tool for estimating copy number profiles in general. To do so, we guide the optimization with an initial set of copy number profiles 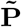 provided by NaiveCopy. A simplified version of guiding the optimization with NaiveCopy’s predicted copy number profiles is described in Supplementary Section A.3.5 and modifications to improve DeepCopy’s accuracy are described in Supplementary Section A.3.6 and A.3.7. Additional mathematical details are provided in Supplementary Section A.3.8 and A.3.9.

### 4.4 Evaluation

DeepCopy’s reinforcement learning optimization was run on a laptop with 96GB of RAM and a 3.6 GHz processor (with 12 cores), without the use of a GPU for all experiments. On 20 simulation instances we compared predicted copy number profiles with the ground truth copy number profiles. To do so, rather than looking at segments, we evaluated the *K* bins with a fixed size of 100kb. Specifically, we defined the accuracy as the average percentage of bins across cells with the unordered allele-specific copy number predicted exactly correctly. To be more precise, let

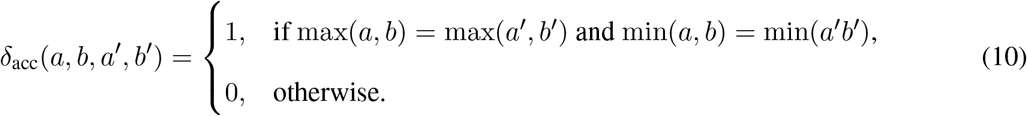

Let *P*_1_, … *P*_*N*_ be the predicted copy number profiles and 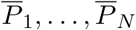 be the true copy number profiles for *N* cells across *K* fixed-size bins in some simulation. Then the accuracy is as below

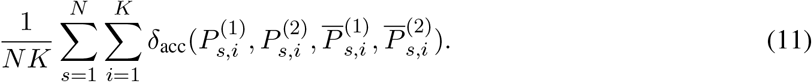

An alternative error metric is the L1 error. Let

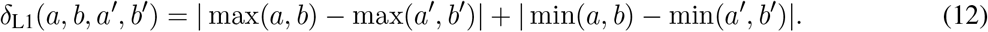

Then, the L1 error metric is defined as

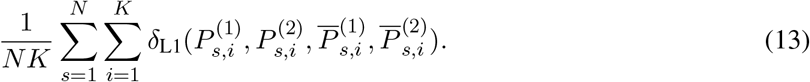

On both simulated and real data, we generated phylogenies for each method and evaluated their parsimony values. To generate these phylogenies, we applied the zero-agnostic copy number transformation (ZCNT) distance [34] to determine a distance matrix between cells and then applied the Neighbor Joining algorithm [52] to determine the phylogeny. The states of internal nodes are calculated using the Sankoff algorithm [53] to solve the small parsimony problem. Finally, parsimony values are calculated with edge lengths determined by the ZCNT distance. Additional details of these calculations are provided in Supplementary Section A.4.2.

Additionally, on real data, we performed validation by utilizing orthogonal SNV data. For ovarian cancer patient OV2295, we used the set of SNV reported in the paper [20]. For breast cancer patient S0, we ran Mutect2 [54] on two pseudobulk samples generated from the single-cell data, one composed of normal cells (4,239 cells) and one composed of tumor cells (5,963 cells), identifying 3,044 SNVs using standard filtering criteria. We restricted to likely truncal SNVs for all analyses. Specifically, for ovarian cancer patient OV2295, we restricted to SNVs that were present in at least 5 cells for all three cell lines. For breast cancer patient S0, we restricted to SNVs present in at least 5 cells for at least 4 out of the 5 sections (noting that one section consists primarily of non-tumor cells). We then generated VAF plots which showed consistency with truncal SNVs typically occurring prior to CNAs (Supplementary Section B.1). Specifically for any copy number {*X*^*A*^, *X*^*B*^} we saw peaks in the VAF around *X*^*B*^*/*(*X*^*A*^ + *X*^*B*^) and *X*^*A*^*/*(*X*^*A*^ + *X*^*B*^) rather than other integer multiples of 1*/*(*X*^*A*^ + *X*^*B*^), indicating that the SNV occurs on all copies of one of the two haplotypes. We utilized this tendency to produce a log-likelihood ratio test that measures the relative probability of an SNV’s observed (reference and variant) reads given the copy numbers predicted by DeepCopy when compared to each alternative method on all cells. We provide a brief description here with full mathematical details in Supplementary Section A.4.3. The probability of a single read being a variant read (as opposed to a reference read) for an SNV on haplotype A for copy number (*X*^*A*^, *X*^*B*^) is *X*^*A*^*/*(*X*^*A*^ + *X*^*B*^). A similar calculation gives these values for reference reads as well as for SNVs that occur on haplotype B. Utilizing these, we calculate the probability of all of the reads for a given SNV assuming it occurred on haplotype A, and the probability of all reads for that SNV assuming it occurred on haplotype B. Since we do not know which haplotype each SNV occurred on, we take the maximum of these two values to estimate the probability of the observed (variant and reference) reads of a given SNV (given the copy numbers of each cell in the position of the SNV). Taking the log of the probability of an SNV given the predictions of DeepCopy and subtracting the log of the probability of that SNV given the predictions of an alternative method gives a log-likelihood ratio for a single SNV. Summing across all truncal SNVs gives a cumulative measurement for truncal SNV support, with positive values supporting DeepCopy’s predictions and negative values supporting the method being compared against. Finally, we performed bootstrapping on the set of truncal SNVs to obtain statistical bounds.

## Data availability

Ovarian cancer patient OV2295 [20] has single-cell FASTQ files available in the European Genome-phenome Archive under accession number EGAS00001003190. Breast cancer patient S0 is available at https://support.10xgenomics.com/single-cell-dna/datasets). Breast cancer patient TN3 [22] has FASTQ files for both ACT and 10x sequencing technologies available in the NCBI Sequence Read Archive under accession number PRJNA629885. Simulated data generated in this paper are publicly available at https://github.com/elkebir-group/DeepCopy.

## Code availability

DeepCopy is implemented in Python using PyTorch and is available at https://github.com/elkebir-group/DeepCopy under the BSD 3-clause license.

## Acknowledgements

We would like to thank the SIGNALS team and in particular Dr. Williams for providing predicted copy number profiles of SIGNALS, CHISEL and Alleloscope on both breast cancer patient S0 and the ovarian cancer dataset. This work was partially supported by the National Science Foundation grant CCF-2046488. This work used resources, services, and support provided via the Greg Gulick Honorary Research Award Opportunity supported by a gift from Amazon Web Services.

## A. Supplementary methods

In Supplementary Section A.1.1 we discuss our data processing steps followed by a description of Naive-Copy (Supplementary Section A.2) and then DeepCopy (Supplementary Section A.3).

### A.1 Data processing

#### A.1.1 Processing raw read counts

We calculated read counts for bins of size *𝓁*_bin_ (set to *𝓁*_bin_ = 100kb by default). To do so, we removed duplicate reads and reads with a very low mapping quality (below 40) We then removed bins with a mappability below 0.8, which removes telomeres and centromeres. Additionally, we remove bins with extreme outlier high read counts. To do so, we calculate the mean and standard deviation of the bottom 99% of bins by read count. Then, outlier bins with read depths above 6 standard deviations above the mean are removed.

The number of reads in a bin is not only a function of the copy number of that bin but also its GC content due to biases in Illumina sequencing, and the bin’s mappability. We calculated the *GC content* of these bins, which is defined as the proportion of bases that are either guanine (G) or cytosine (C), and utilized existing estimates of mappability. We then applied LOWESS regression to correct read depths for GC bias and mappability bias. Specifically, we first predict the read depth from mappability and then divide by this value. Then for each cell, we predict the read depth from the GC content and again divide the read depth by this value. This type of bias correction utilizing LOWESS regression is common in copy number calling methods such as CHISEL [29]. After we removed outlier bins and applied GC bias correction, we obtained a read depth vector 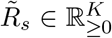 for each cell *s* where *K* is the number of 100kb bins after filtering.

Additionally for mathematical convenience, we scaled 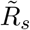 to have an average value of 1 across all bins for each cell *s*. For our real data, the number *K* of bins was 26,033 for ovarian cancer patient OV2295, 26,135 for breast cancer patient S0, 26,041 and 26,141 for breast cancer patient TN3 sequenced with ACT and 10x technologies.

#### A.1.2 Determining haplotype-specific read counts

To be able to infer allele-specific and haplotype-specific copy numbers, we determined the numbers 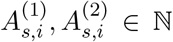 of reads in each bin *i* and cell *s* that can be mapped to the two parental haplotypes, respectively. To do so, we first pooled all of the single cell BAM files together into a pseudobulk sample. Then, we ran bcftools’ (1.9) [55] `mpileup` and `call` commands on this pseudobulk sample to obtain SNP positions and alleles. We then ran SHAPE-IT 4 [56] in combination with the 1000 genomes reference panel [57] to phase these SNPs into haplotype blocks. Next, we reapplied bcftools’ `mpileup` and `call` commands to the individual cells in the positions of these phased SNPs to determine the SNP counts for individual cells. We then converted these to cumulative counts for the A allele and B allele in haplotype blocks for each cell. Finally, we phased these haplotype blocks with a novel algorithm with several theoretical advantages including the ability to phase haplotype blocks when the average copy number for the two alleles is balanced across cells. We discuss this algorithm in the following.

##### Simplified description of the phasing algorithm

The simplest way of phasing haplotype blocks is to simply define the allele with the lower count across all cells in any haplotype block to be the B allele. However, as noted in the SIGNALS paper [30], this may be ineffective in the case where only a small subset of cells have an imbalanced copy number between the two alleles. This is because in that case, imbalances in allele counts due to noise in cells with a balanced copy number may overwhelm imbalances in the small number of cells with imbalanced copy numbers. Additionally, it is possible for the average copy number across cells to be balanced despite individual cells having an imbalanced copy number. For instance, if some cells have an amplification on one allele and other cells have an amplification on the other allele. To overcome these challenges, we begin by describing a simple algorithm that would work well assuming the copy number is constant across each chromosome (but differs across cells). Then, we will describe a modification to this algorithm that allows for accurate haplotype phasing without that assumption.

Running SHAPE-IT yields a partition of the reference genome into haplotype blocks. Let 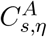 and 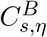 be the number of counts of the first and second allele respectively, for cell *s* and haplotype block *η* (in some fixed chromosome). Note that the first and second alleles are defined arbitrarily since the haplotype blocks are not yet phased. The probability of observing values of 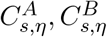 assuming there is no imbalance comes from a binomial distribution which can be approximated by a Gaussian. Specifically, 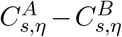 has a mean of zero, and a variance equal to the total number of counts 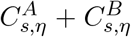. Consequently, dropping constant terms, observing the imbalance 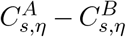 has a probability proportional to exp 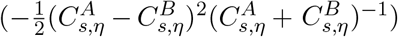 .Thus, the log probability evidence for imbalance is 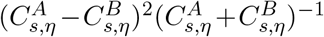 (again dropping a constant factor). Including the sign of the imbalance (which is necessary to keep track of for phasing haplotype blocks) gives 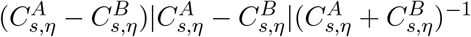 Define

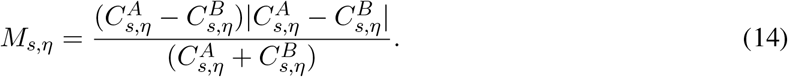

Intuitively, |*M*_*s,η*_| is proportional to the negative log probability of the absence of an imbalance towards the direction sign(*M*_*s,η*_). In other words, if |*M*_*s,η*_| is large then there is very strong evidence of imbalance toward A allele if *M*_*s,η*_ *>* 0 and the B allele if *M*_*s,η*_ *<* 0.

For any two haplotype blocks *η* and *η*′ in the same chromosome, we want to phase the haplotypes such that the evidence for imbalance in any cell *s* points in the same direction for both haplotype blocks. In other words, we want to phase the two haplotype blocks such that if cell *s* has strong evidence for an imbalance in favor of the A allele in haplotype *η* then it should also be imbalanced in favor of the A allele in haplotype block *η*′. Mathematically, we want *M*_*s,η*_ *·* 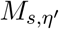to be positive for any cell *s* and haplotype blocks *η* and *η*′, especially if the magnitude of *M*_*s,η*_ and 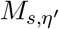are large (indicating strong evidence for imbalance). Note that swapping the phasing of haplotype block *η* simply multiplies *M*_*s,η*_ by *−*1 for all cells *s*. Define *p*_*η*_ as our final phasing, which is either 1 or *−*1 for any haplotype block *η*. We want to maximize *p*_*η*_*M*_*s,η*_ *·*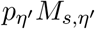for all cells *s* and for all haplotype blocks *η* and *η*′. Theoretically, if there were zero noise then *M*_*s,η*_ should simply be proportional to the actual copy number imbalance in cell *s* multiplied by the number of counts in the haplotype block *η* and perhaps multiplied by *−*1 due to incorrect phasing. Let **M** = [*M*_*s,η*_] be the matrix with entries *M*_*s,η*_ across all cells *s* and haplotype blocks *η* on some chromosome. We can remove much of the noise by replacing **M** with a rank 1 matrix approximation produced by a singular value decomposition. That is, we have **M** *≈ XY* ^*T*^ such that *M*_*s,η*_ *≈ X*_*s*_ *· Y*_*η*_ where *X*_*s*_ represents a scaled version of the actual imbalance in cell *s* and *Y*_*η*_ represents a multiplicative factor which contains information on the phasing and total count of the haplotype block *η*. Then, we are simply maximizing 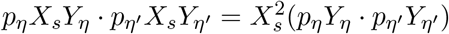. However, this is very easily accomplished by setting *p*_*η*_ = sign(*Y*_*η*_) for all haplotype blocks *η*, so that *p*_*η*_*Y*_*η*_ *· p*_*η*_*′ Y*_*η*_*′* is always positive.

##### Additional phasing algorithm details

The approach described above far works perfectly if one assumes that the copy number mostly remains constant for all cells in each chromosome, i.e., most CNAs only affect entire chromosomes. However, this is not a realistic assumption as many tumors have chromosome-arm CNA events as well as smaller, focal CNAs. The simplest modification of this approach is to apply it to individual bins where one assumes the copy number remains approximately constant in each bin. However, utilizing information across an entire chromosome to eliminate noise and determine which cells are truly imbalanced and in what pattern is very useful. Thus, it is ideal to first determine patterns of imbalances that occur in subsets of the chromosome and then utilize these patterns to phase haplotype blocks. To accomplish this, we first performed a singular value decomposition to approximate **M** = [*M*_*s,η*_] with a higher, rank 10 matrix 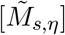. The purpose of this approach is to remove noise utilizing information across the entire chromosome to find patterns of imbalances while allowing for differences to exist in different subsets of the chromosome. Then, using groups of 100 consecutive haplotype blocks, we approximated 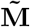 with a rank 1 matrix for each group. We phased each group of 100 haplotype blocks as previously described. To reconstruct the final phasing, we updated the phasing of these groups of haplotype blocks by setting their phase to best match an exponential moving average of previous sets of 100 haplotype blocks with a smoothing factor of 0.9. Specifically, assume the first *K*_phase_ sets of 100 haplotype blocks have already been phased. Let 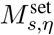 indicate the average value of 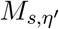for *η*′ values in the *η*th set of 100 haplotype blocks.

Assume 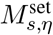 has already been phased for *η ≤ K*_phase_. Let 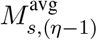 be the moving exponential moving average of 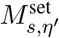 for *η*′ *< η*. Then the moving average 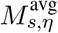 is defined as 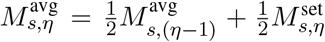 Additionally, 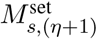 is phased by either multiplying by 1 or *−*1 depending on which phasing maximizes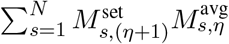

Once all of the phasing is completed, we determined the vector *p*_*η*_ such that *p*_*η*_ is either *−*1 or 1 for each haplotype block *η* depending on the phasing. Using this we defined the phased haplotype block counts as

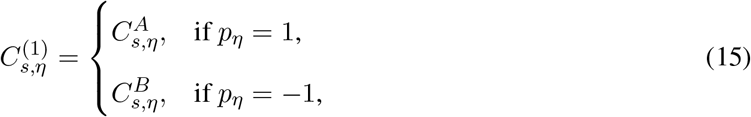

and

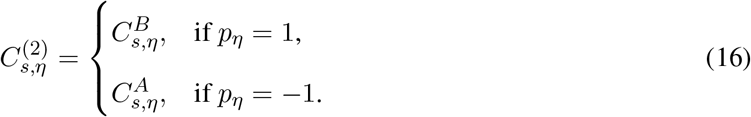

Finally, we can define the haplotype-specific count vectors 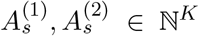. Specifically, 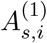 is defined as the sum of 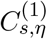 for all haplotype blocks *η* in the bin *i*. Similarly, 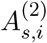 is defined as the sum of 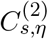 for all haplotype blocks *k* in the bin *i*.

Once 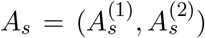 has been determined, we must correct for one additional issue. Specifically, invalid haplotype block phasing can occur when all cells have a complete loss of heterozygosity. Specifically, if all cells have a loss of heterozygosity for some bin (and no normal cells are provided) then haplotype blocks cannot be determined, and there will be near-zero counts for both haplotypes. To correct this, if some bin has less than one haplotype-specific count per cell (across all cells) then a pseudo-count of five is added to the first haplotype. Thus, regions with a complete loss of heterozygosity across all cells have a BAF near zero, rather than having a random BAF determined by noise and incorrect reads.

#### A.1.3 Estimating segments and noise levels

Given read depths 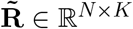and haplotype-specific counts **A**^(1)^, **A**^(2)^ ∈ *ℕ*^*N ×K*^ for *N* cells and *K* fixed-size bins, we now describe how we segmented the *K* bins into *L* segments and obtained read counts and corresponding variances **R, Σ**^*R*^ ∈ *ℝ*^*N ×L*^ and BAFs and corresponding variances **B, Σ**^*B*^ ∈ *ℝ*^*N ×L*^ for *N* cells and *L* segments. Our segmentation algorithm relies on estimates of the mean and variance of the read depth and BAF of consecutive bins to identify breakpoints. Therefore, before describing the segmentation algorithm, we will first discuss how to estimate the mean and variance of the read depth and BAF of a set Λ of consecutive fixed-size bins.

##### Estimating noise in segments

We are given a set Λ of fixed-size bins *i* for a cell, each of which has a read depth value 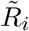 and a number 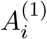, 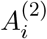 of counts of each haplotype. Additionally, the read depth vector is scaled to have an average value of 1 for mathematical convenience. We then estimate an average read depth and BAF for the set Λ of bins as well as a noise level in these estimates. The noise level refers to an estimated standard deviation of these average values. Additionally, this noise level can be squared to give estimated variances.

Calculating the average is trivial for the read depth (taking the mean value across bins) and finding the average BAF can be done by first summing haplotype counts across bins and then calculating the BAF from this sum. More specifically, the average read depth is as below

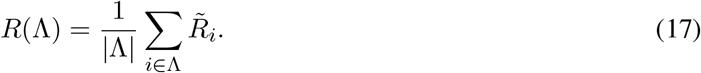

Additionally, the average BAF is given below

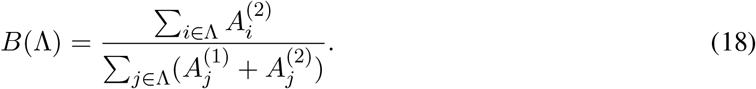

For calculating the noise in the read depth, we first start with a simple statistical estimate of the noise in the set Λ of bins. This simple estimate starts with the assumption of independent noise across the bins in Λ. Define 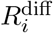 as 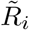 minus the mean value of 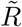 in the segment.

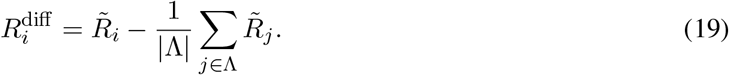

The simplest way of calculating the read depth noise is by utilizing the standard error of the Λ measurements given below

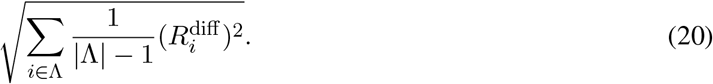

In practice, there might be locations-specific noise for bins in Λ even if they have identical copy numbers and even after GC-bias correction. Therefore, we modify the simple estimate with a heuristic in order to achieve better performance in the case where the noise across bins is correlated. Specifically, our heuristic is approximately proportional to the standard error in the case that the noise is uncorrelated across bins, but increases substantially when the noise is highly correlated across bins. To accomplish this, we use the fact that the Fourier transform of independent noise is uniform across the frequency spectrum, whereas the Fourier transform of spatially localized noise (with high auto-correlation) is disproportionately low frequency. We define *F* = [*F*_1_, …, *F*_|Λ|_]^*⊤*^ as the Fourier transform of *R*^diff^ across the bins *i* ∈ Λ. The Fourier transform preserves the sum of squares such that the squared standard error is also calculated below

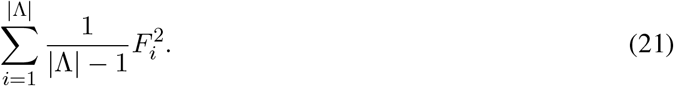

If the noise was uncorrelated across bins, we would expect an equal amplitude across the different frequencies. However, a higher amplitude of low frequency values would indicate auto-correlation in the noise values. Lower frequency values in the noise result in a higher error in the estimated average read depth. As a simple example, imagine the first |Λ|*/*2 bins all have one value, and the second |Λ|*/*2 bins all have some other value. Imagine this is due to the error in the read depth coming from some unknown bias which affects the first |Λ|*/*2 bins differently than the second |Λ|*/*2 bins. Then, the error in our estimate of the average read depth would be equivalent to the scenario in which we only had two bins that we are averaging, since we only have two independent samples of the noise. Consequently, the squared error in our average read depth measurement is proportional to the inverse frequency of the noise. Specifically, we sum the square of each component in the Fourier transform divided by |Λ| times the frequency of that component to get the square of the error. If the frequency were equal to 1*/*|Λ|, this would give the standard error, however, the error increases as the frequency of the noise decreases. Define Λ_*i*_ as |Λ| multiplied by the frequency of the *i*th component of the Fourier transform. Note that there is no zero-frequency component since *R*^diff^ has an average value of 0 by definition. Thus Λ_*i*_ is at most |Λ|. Then our heuristic estimate of the variance is the below slight modification to equation (21)

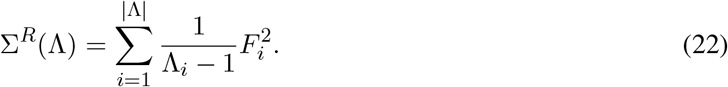

The simplest way of estimating the noise in the average BAF is to only consider the total SNP counts across all bins. Each count can be treated as its own independent random variable with a value of either 0 for the A allele or 1 the B allele. Let *C*_*a*_ be the number of counts of the A allele and *C*_*b*_ be the number of counts of the B allele. Then, the BAF is

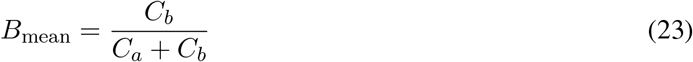

and the variance in the BAF is

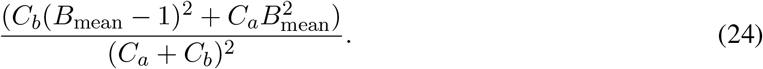

To avoid zero noise in the case where the BAF is either 0 or 1, we add one pseudocount to *C*_*a*_ and *C*_*b*_. This estimate is generally accurate, however, it assumes all counts are sampled from the same distribution across the Λ bins. This assumption may be broken due to incorrect phasing of haplotype blocks, very small focal CNAs within a segment or other errors in determining the counts of individual SNPs. To correct for this, we also note that the average BAF is equivalently calculated as the weighted average of the BAF of all Λ bins, weighted by the total SNP count in each bin. Let 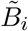 be the BAF of each of the bins *i* ∈ Λ as below

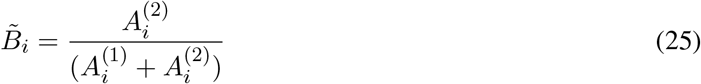

and let 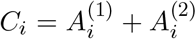 be the total count for both alleles in each bin *i*. The variance in the average BAF is then approximately

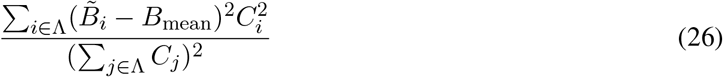

assuming the counts are spread across a large number of bins. The unbiased estimator has a slightly different denominator causing the error to go to infinity rather than 0 if there is a single bin. However, this estimate is only needed for a corrective term to add to the noise in the case where different bins appear to have SNP counts drawn from different distributions. Consequently, we achieve an effective estimate of the noise by using

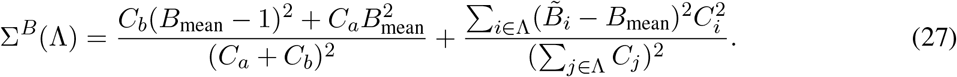

As an example, in the case where only one bin has a non-zero count, the second term disappears and we simply estimate the noise from the allele counts in that bin. However, in the case where one bin has allele counts of (100, 0) and another bin has allele counts of (0, 100) the second term prevents us from falsely assuming we have a very low noise BAF of 0.5 with variance 1*/*800, resulting in an adjusted variance of 1*/*800 + 1*/*8.

#### Estimating segments

Our system starts with *K* bins of size *𝓁*_bin_ (set to *𝓁*_bin_ = 100kb), and then merges these together into *L* segments where it is believed each cell has a constant copy number within each of these segments. To determine the boundary of these segments, one must find which positions in the genome have substantial evidence for a breakpoint in the copy number for some cells. Specifically, the evidence for a breakpoint is determined within each cell, and then this evidence is summed across cells. To determine the evidence for a breakpoint within a cell, we must determine the evidence that the last *Γ* bins prior to the breakpoint have a different read depth or BAF than the next *Γ* bins after the breakpoint. Larger values of *Γ* allow evidence to be accumulated across more bins, yet smaller values of *Γ* allow for the detection of smaller copy number aberrations. Consequently, we utilized *Γ* ∈ {10, 20, 40}. This allows for the detection of small 1Mb copy number aberrations, while also allowing for evidence to be accumulated across 4Mb regions when detecting the boundary of larger copy number aberrations that may occur in only a small subset of cells.

Let Λ_before_, Λ_after_ be the set of *Γ* bins before and after some possible breakpoint position, respectively. Then the evidence for a breakpoint at that position in that cell (from the read depth and utilizing that value of *Γ*) is

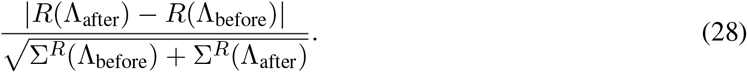

The analogous computation is true for BAF:

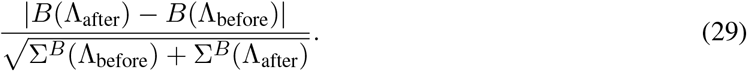

This computation is done for all possible breakpoints, for all cells, for each of the three values of *Γ*, and for both the read depth and BAF. The evidence is summed across cells, giving 6 values for the evidence of each possible breakpoint. Our segmentation algorithm works by iteratively picking breakpoints with the most evidence but not allowing for two breakpoints to be close to each other and thus relying on the same evidence. For instance, if there is a large amount of evidence for a breakpoint at position *x*, then there will likely also be a large amount of evidence at position *x*+1, since the set of bins before and after the breakpoint are very similar for position *x* and position *x* + 1. Thus, if one breakpoint uses *Γ*_1_ bins, and another uses *Γ*_2_ bins, then they must at least be a distance of min(*Γ*_1_, *Γ*_2_) bins apart. With this restriction in mind, the eligible breakpoints with the most evidence are iteratively added until no breakpoint has an evidence value of at least 3 times the number of cells. Note that this cutoff must be proportional to the number of cells since breakpoints will receive an evidence value near the number of cells just due to random chance. To avoid bins larger than 20Mb, this process is then applied again with the value of *Γ* = 200, iteratively adding breakpoints until there are no eligible breakpoints left (and thus all breakpoints are within 20Mb of some other breakpoint).

Once the set of segments is determined we set *R*_*s,i*_,*B*_*s,i*_, 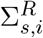and 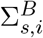 to be the values of *R*(Λ(*i*)), *B*(Λ(*i*)), Σ^*R*^(Λ(*i*)), and Σ^*B*^(Λ(*i*)) for cell *s* where Λ(*i*) is the set of fixed-size bins comprising segment *i*.

### A.2 NaiveCopy

The goal of the NaiveCopy pipeline is to produce the inputs required by DeepCopy. NaiveCopy takes as input measurements **R, B, Σ**^*R*^, **Σ**^*B*^ ∈ *ℝ* ^*N ×L*^ obtained using the previously described data processing steps and produces preliminary copy numbers 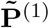,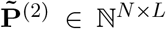 for *N* cells and *L* segments. These copy numbers help guide DeepCopy’s reinforcement learning search. In this section, we describe the process of generating these preliminary copy number profiles.

#### A.2.1 Defining measurement probabilities

In this section, we define the probability 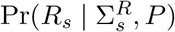 of read depth values *R*_*s*_ and the probability 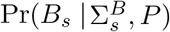 of BAF values *B*_*s*_ given a copy number profile *P* and estimates of the variances 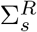, 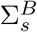 of these values of a cell *s*. This is utilized in producing copy number profiles for both NaiveCopy and DeepCopy. Given the copy number profile *P* = (*P* ^(1)^, *P* ^(2)^), we can predict the read counts to be proportional to *P* ^(1)^ + *P* ^(2)^. Thus, if we predict the correct copy number profile, we predict *c ·* (*P* ^(1)^ + *P* ^(2)^) to match *R*_*s*_ on expectation for some scaling constant *c*. Given the central limit theorem, it is reasonable to assume measurement values come from a Gaussian distribution. Then, the probability 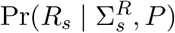 of observing *R*_*s*_ given *P* and 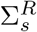 is as follows.

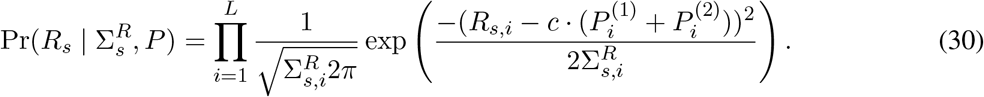

Note that this expression is minimized by minimizing the below total squared error.

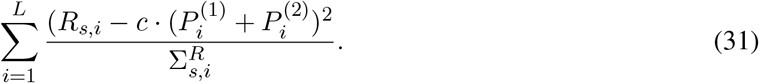

For evaluating a copy number profile *P* as used by DeepCopy, we set *c* to 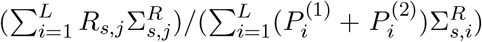 in order to minimize this error and maximize the probability. For NaiveCopy, we set *c* = *γ*^*−*1^ where *γ* is the cell-specific scaling factor discussed in Supplementary Section A.2.2.

The expected value of the B-allele frequency in bin *i* given the copy number profile *P* is 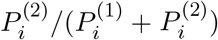. Similarly as for the read count, for the B-allele frequency, we have the below equation.

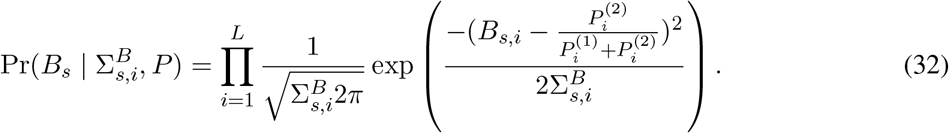

Again, maximizing this probability is equivalent to minimizing the blow total squared error

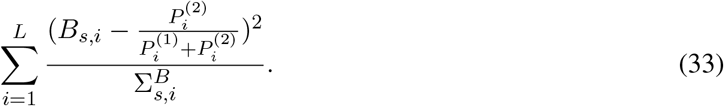

#### A.2.2 Estimating cell-specific scaling factor

Given a ground truth profile *P*, the ground-truth cell-specific scaling factor *γ*_true_ is defined as the average total copy number 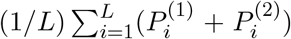. In this section, we utilize variance aware segmentation to estimate the cell-specific scaling factor given an allele counts vector *A* = [*A*^(1)^, *A*^(2)^]^*⊤*^ ∈ *ℕ*^2*×K*^ and a read depth vector 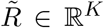. These vectors are modified to use 1Mb bins formed out of merging our original *K* bins of fixed size 100kb. These larger bins allow for a more efficient exhaustive search of all possible segments within each chromosome. Additionally, any loss of precision due to using large bins is not relevant since very small copy number aberrations are not relevant for determining the cell-specific scaling factor.

##### Determining low variance segments

Before describing our approach, we give a brief conceptual motivation. If one had perfect noise-free read depth and B-allele frequency measurements, then determining the cell-specific scaling factor would be trivial. One could simply select the smallest cell-specific scaling factor such that integer copy numbers can result in the observed read depth and allelic imbalance values. Similarly, if one had a list of highly accurate pairs of read depth and B-allele frequency measurements, one could easily find the cell-specific scaling factor that best fits the measurements. In reality, read depth and B-allele frequency measurements are very noisy. As such, our method works by first finding low-noise segments that provide highly accurate measurements of the B-allele frequency and the mean read depth within those segments. Then we evaluate if those high-quality read depth and BAF values would be possible for a given value of *γ*. Finally, we determine which value of *γ* best matches these observed measurements.

We will define 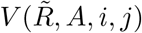 as a function that inputs the read depth vector 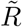, the allele-specific count vector *A*, some starting bin *i* and some ending bin *j*, and outputs a measurement of how accurately we know the mean 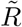 and BAF value within that segment. If 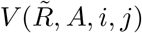 is high, then this implies an accurate measurement of a mean read depth and B-allele frequency value pair that must be possible to observe for a given cell-specific scaling factor. The goal is to split the 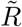 and *A* vectors into segments that maximize this value. We define 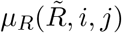 as the mean value of 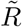 on the interval from *i* to *j*, i.e.,

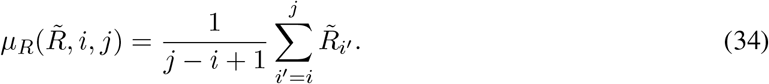

We define 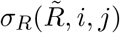 as the standard error of this mean value, i.e.,

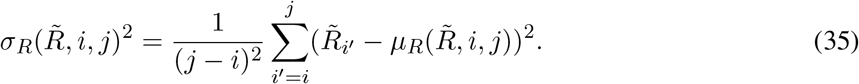

We define *μ*_*B*_(*A, i, j*) as the B-allele frequency of the total allele-specific counts in *A* on the interval *i* to *j*, i.e.,

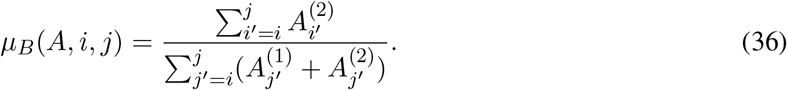

Finally, we define *σ*_*B*_(*A, i, j*) as the noise estimate for *μ*_*B*_(*A, i, j*) as described in Supplementary Section A.1.3.

Let *L*_min_ be the minimum length segment (in terms of *𝓁*_bin_ = 100kb bins) we consider acceptable for giving an estimated error in the average read depth and B-allele frequency values within that segment. We use *L*_min_ = 5 as a default setting. Taking the inverse of our error estimate gives a measurement of precision. Thus, adding together an inverse of these error values for the read depth and B-allele frequency gives

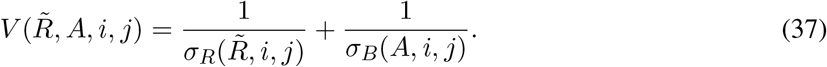

Now that we have a measurement of the accuracy of read depth and BAF values provided by some segments, we can move on to defining the problem of determining an optimal variance-aware segmentation. Define *A*_*S*_ as the set of all possible lists of non-overlapping segments of length at least *L*_min_ on *L* bins such that each segment is contained within a chromosome. More precisely, define 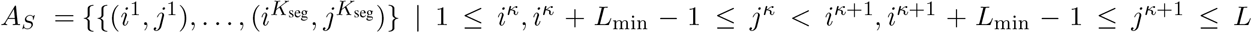 for all κ ∈ {1, …, *K*_seg_ − 1} and *i*^*κ*^ and *j*^*κ*^ are on the same chromosome for all *κ* ∈ {1, …, *K*_seg_}}. We then have the following problem of variance-aware segmentation.

###### Problem 2

(Low Variance Segments). *Given a read depth vector* 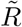 *and a allelic imbalance vector B, find the list of segments* 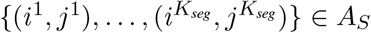 *that maximize* 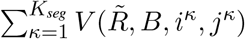.

Solving Pustive search is computationally infeasible. Therefore, we use a greedy approach as an approximate solution. Specifically, within each chromosome, we first find segment *i* to *j* of length at least *L*_min_ that maximize 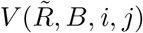. Then, iteratively, we find the segment *i* to *j* of length at least *L*_min_ that maximizes 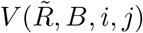 without overlapping with any existing segment. The process stops when it is not possible to add any additional intervals of length at least *L*_min_ without having overlapping segments. The values of 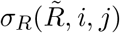 and *σ*_*B*_(*A, i, j*) can be calculated efficiently for each segment once the appropriate cumulative sums of input vectors and their squares have been stored. For instance, once the cumulative sum across the genome of 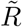 and 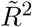 have been stored, 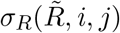 can be calculated in constant time independent of the length of the segment.

##### Determining cell-specific scaling factor from low variance segments

After finding an approximate solution to Problem 2, we have a list of segments 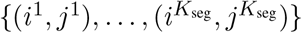. For each of these segments, we have an estimated mean read depth and allelic imbalance as well as an estimated error in that estimate. Define 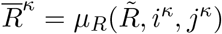, 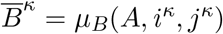, for *κ* ∈ {1, …, *K*_seg_}. Also define 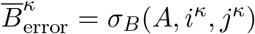, and 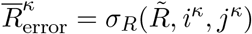.

Given a cell-specific scaling factor *γ*, possible values for the read depth and allelic imbalance are of the form (*N*_1_ + *N*_2_)*/γ* and min(*N*_1_, *N*_2_)*/*(*N*_1_ + *N*_2_) respectively where *N*_1_ and *N*_2_ are both non-negative integers representing haplotype copy numbers. Define *f*_*R*_(*N*_1_, *N*_2_, *γ*) = (*N*_1_ + *N*_2_)*/γ*, and *f*_*B*_(*N*_1_, *N*_2_) = *N*_2_*/*(*N*_1_ + *N*_2_). We define the equation for the minimum error of a read depth and allelic imbalance measurement given a cell-specific scaling factor as

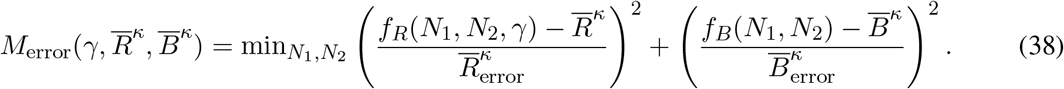

We note that the number of possible *f*_*R*_(*N*_1_, *N*_2_, *γ*) and *f*_*B*_(*N*_1_, *N*_2_) values while keeping *f*_*R*_(*N*_1_, *N*_2_, *γ*) within some range (for instance 0 to 1) is proportional to *γ*. Therefore, for random read depth and allelic imbalance values, one would expect the squared error 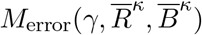 to scale proportional to *γ*^*−*2^. Consequently, our goal is to minimize 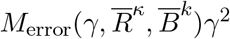 rather than simply minimizing 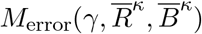. As further intuition, note that 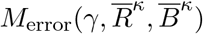 can be made arbitrarily small for any 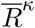,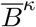 within the range of possible values by setting *γ* arbitrarily large. Putting this all together, our goal is to select *γ* to minimize the total error

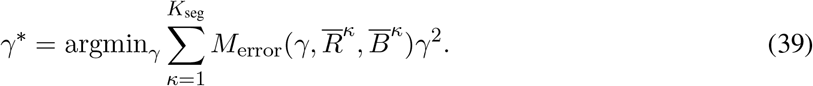

Additionally, we apply the constraint *γ ≥* 1 to avoid the solution of setting *γ* extremely low while predicting copy numbers of all zeros for all bins. This single variable minimization is accomplished in our code by a simple 1-dimensional grid search (in our implementation the grid search starts at 1 and ends at 10 with exponential increments of exp(0.02) followed by a secondary local search around the optimal solution with increments of exp(0.002)). As a slight caveat, if there exists an additional value of *γ* that approximately minimizes 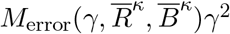 (off by at most 10), and is not within a ratio of 20% of *γ*^*∗*^, then this value of *γ* is also reported. For most cells, this does not occur, but for a few cells there exists multiple similar quality cell-specific scaling factors.

#### A.2.3 The predicted profiles of NaiveCopy

In Supplementary Section A.2.2 we gave a method for predicting cell-specific scaling factors. These xscaling factors can then be utilized to give a basic prior estimate of the copy number profiles 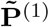, 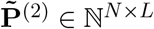 for *N* cells and *L* segments. As derived in Supplementary Section A.2.1, the log probability log 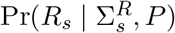 of a read depth vector *R*_*s*_ given variances 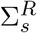 the copy number profile *P* is proportional to

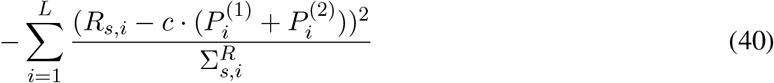

where *c* is a constant. In the case that we know the cell-specific scaling factor *γ*, this becomes the below expression.

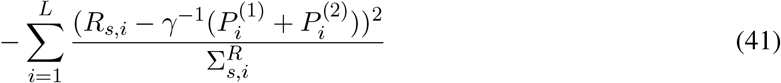

Similarly, the log probability of the BAF vector *B*_*s*_ given the copy number profile *P* is proportional to

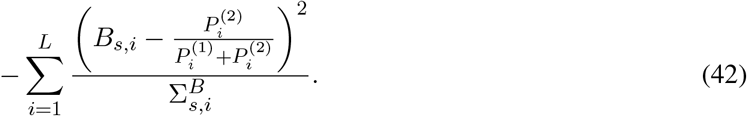

Consequently, the optimal copy number profile 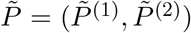 for maximizing 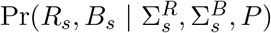 minimizes the below expression.

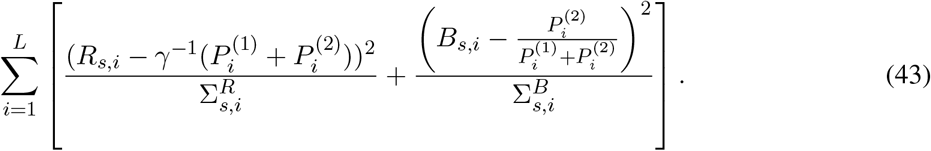

Fortunately, the error term for each bin *i* only depends on the copy numbers for that bin 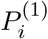 and 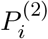 so we simply select these values to minimize the error for each bin. Applying this to the BAF vector and read depth vector for each cell (given the cell-specific scaling factor for this cell) gives NaiveCopy’s predicted copy number profiles 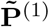, 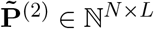 for each of the *N* cells and *L* segments.

### A.3 DeepCopy

#### A.3.1 Generating copy number profiles

We have defined *g*(*G*) as the copy number profile created by starting with the normal copy profile 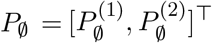 such that 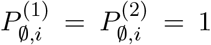 for each segment *i*, and applying the CNAs in *G*. Here, we define this precisely. First, we define a function *f*_*g*_(*P*, [*c*]), which takes as input a copy number profile *P* and a single CNA tuple *c* = (*c*_WGD_, *c*_hap_, *c*_start_, *c*_end_, *c*_value_) and outputs a new copy number profile *f*_*g*_(*P*, [*c*]) = [*Q*^(1)^, *Q*^(2)^]^*⊤*^ such that

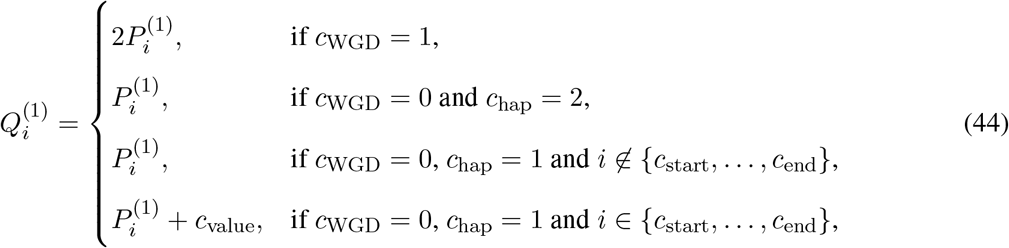

and

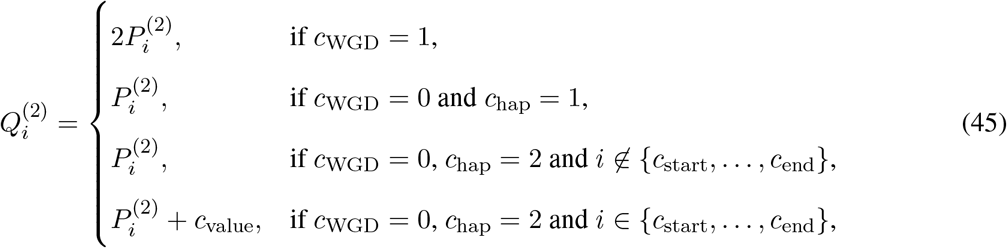

With this definition, we now inductively define

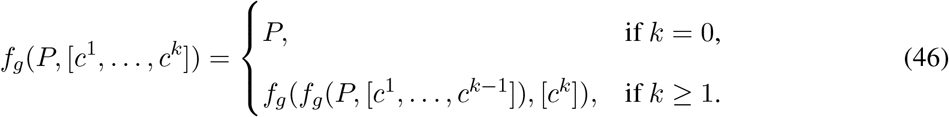

Finally, we define *g*(*G*) = *f*_*g*_(*P*_*∅*_, *G*).

#### A.3.2 Neural network architecture

Each input copy number profile *P* = [*P* ^(1)^, *P* ^(2)^]^*⊤*^ first has its values capped at a maximum value of 19.

In other words, if 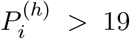, then we set 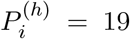. Consequently, each element in 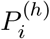 can take 20 values ({0, 1, … 19}). This value is one-hot encoded so that *P* becomes an *L ×* 2 *×* 20 tensor. This tensor is then reshaped into an *L ×* 40 tensor by concatenating the one-hot encoded copy numbers for both haplotypes for each of the *L* segments. An embedding network then converts this to a 500 dimensional vector. Specifically, the embedding network applies convolutions of size 5 across the bins. There are a total of 400 such convolutions, changing the second dimension of the copy number profile representation from 40 to 10 (going from 40 channels to 10 channels). The copy number profile representation tensor is then flattened, and a hyperbolic tangent non-linearity is applied. Finally, a fully connected layer converts this tensor to a 500 dimensional embedding of the copy number profile. The starting and ending position of each CNA are encoded as one-hot *L* dimensional vector. A fully connected layer also converts these to 500 dimensional representations. When determining the end position, the embedding of the start position is added to the embedding of the copy number profile. When determining the copy number, the embeddings of both the start and end positions are added to the embedding of the copy number profile. For predicting the start position, the end position, or copy number, a hyperbolic tangent non-linearity is applied followed by a fully connected layer. Additionally, the softmax function is applied to convert the outputs into probabilities. Predicting whether or not to terminate the generating sequence is done in the same manner as predicting the start position.

#### A.3.3 Policy learning equation derivation

In the main text, we state the equation

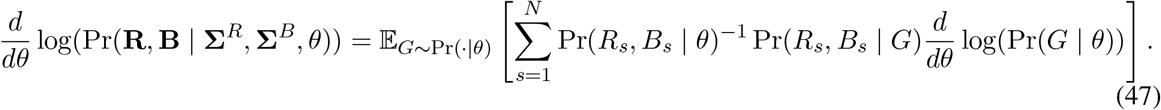

As stated in the main text, we use the shorthands 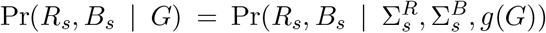and 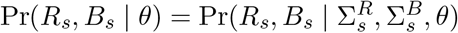. Below is the derivation of this equation.

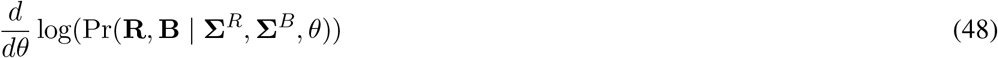

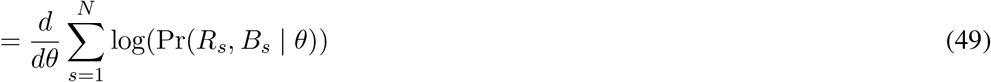

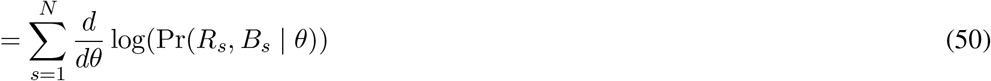

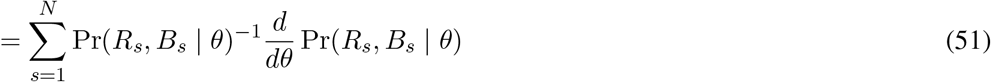

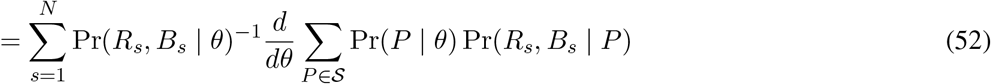

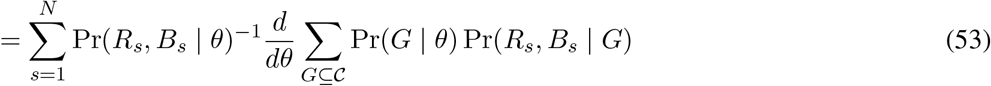

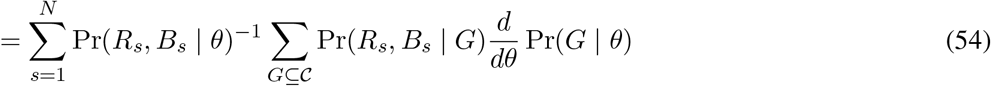

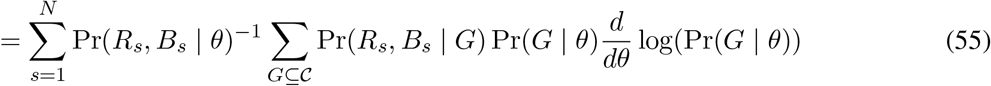

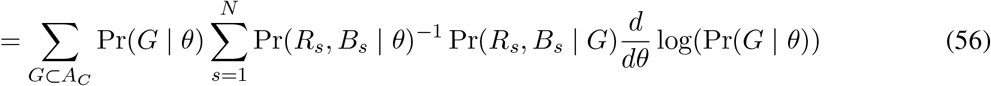

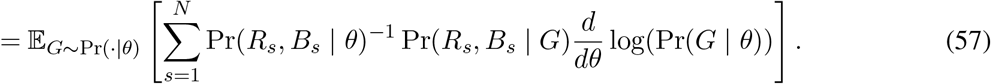

#### A.3.4 Probability estimation via sampling

In several steps of our method, we need to estimate probabilities using sampling. Specifically, we need to estimate Pr(*R*_*s*_, *B*_*s*_ | *θ*) for the reward function calculation, and we need to estimate Pr(*P* | *θ*) for copy number profile prediction. Let 𝒜(*P*) be the set of generating sequences that efficiently generate *P*, as defined in Supplementary Section A.3.8. Let *P*_1_, …, *P*_*M*_ be the copy number profiles used to guide our search as described in Supplementary Section A.3.5. Note *g*(*G*) = *P* if *G* ∈ *A*(*P*). To estimate Pr(*R*_*s*_, *B*_*s*_ | *θ*), we use

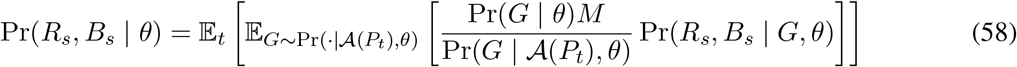

where *t* is sampled uniformly from {1, …, *M* }. Note that this value changes during training and so we re-estimate it after each gradient update to the model’s parameters *θ*. Define *f*_same_(*P, P*′) = 1 if *P* = *P*′ and 0 otherwise. To estimate Pr(*P* | *θ*), we use

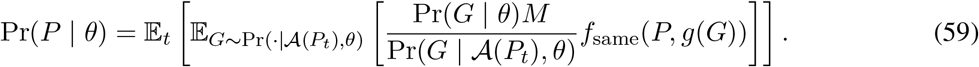

#### A.3.5 Guiding the optimization with initial copy number profiles

In order to guide the optimization, we start with a reasonable prior on generating sequences of CNA tuples. In this section, we describe a slightly simpler version of this prior, which is then improved on in Supplementary Section A.3.6 and A.3.7. We start with a set *P*_1_, …, *P*_*M*_ of plausible copy number profiles. Specifically, we utilize the unique profiles predicted by the NaiveCopy, as well as a set of additional copy number profiles determined during training as described in Supplementary Section A.3.7. As such, *M ≥ N*. Then, we restrict our generating sequences to efficiently generate copy number profiles in {*P*_1_, …, *P*_*M*_ }. We define 𝒜(*P*) as a set of generating sequences that efficiently generate *P*. The exact definition is given in Supplementary Section A.3.8, but for now, we simply state *g*(*G*) = *P* if *G* ∈ 𝒜(*P*). Define 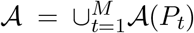. We define Pr(*G* | 𝒜(*P*), *θ*) as the probability of sampling *G* when restricting to 𝒜(*P*) which is defined precisely in Supplementary Section A.3.9. To sample when restricted to 𝒜 we first uniformly sample *t* from {1, …, *M* } and then sample *G* from Pr(*G* | 𝒜(*P*_*t*_), *θ*). Thus, 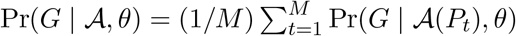.

Define *B*(*G, 𝒜*) = 1 if *G* ∈ 𝒜 and 0 otherwise. To restrict generating sequences to 𝒜, we multiply *r*(*G*) (defined in equation (8)) by *B*(*G, 𝒜*) in the original objective function. Since we are modifying our sampling procedure, we must also modify our training to compensate for that.

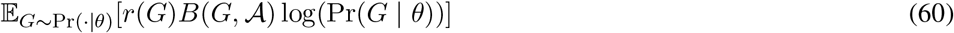

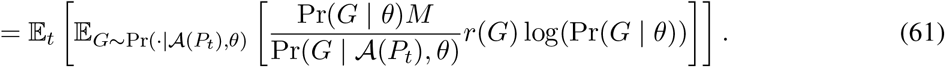

Define *r*′(*G, t*) = Pr(*G* | *θ*)*M* Pr(*G* | 𝒜(*P*_*t*_), *θ*)^*−*1^*r*(*G*), as our new reward function for our new sampling procedure. For those familiar, note, the term Pr(*G* | *θ*)*M* Pr(*G* | 𝒜(*P*_*t*_), *θ*)^*−*1^ is the adjustment used for importance sampling [58] (since Pr(*G* | 𝒜(*P*_*t*_), *θ*)*/M* is the sampling probability). We now have the objective function

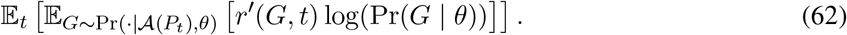

After learning the model parameters *θ*, we utilize sampling to estimate which copy number profile *P* maximizes Pr(*R*_*s*_, *B*_*s*_ | *P*) Pr(*P* | *θ*) for each cell *s* as described in Supplementary Section A.3.4.

#### A.3.6 A modification to improve the removal of spurious CNAs

The prior copy number profile estimates provided by NaiveCopy are likely to contain fake CNAs that only appear to exist due to noise. Our method described thus far should correct for this so long as *P*_1_, …, *P*_*M*_ contains most of the correct copy number profiles in addition to some profiles with fake CNAs. However, in high noise data sets it is possible for most or all of the NaiveCopy estimates to contain fake CNAs. To correct for this, our method must delete fake CNAs. To accomplish this, we allow our method to predict copy number profiles *g*([*c*^1^, …, *c*^*k*^]) for any subsequence [*c*^1^, …, *c*^*k*^] of [*c*^1^, …, *c*^*k*^, …, *c*^*k*+*r*^] ∈ 𝒜. Note, *g*([*c*^1^, …, *c*^*k*^]) is the copy number profile *g*([*c*_1_, …, *c*_*k*_, …, *c*_*k*+*r*_]) but with the last *r* CNAs removed. Thus, more intuitively but less precisely, we allow our model to predict copy number profiles which are like some profile in *P*_1_, …, *P*_*M*_ but with some CNAs removed. The details of this modified sampling procedure are described in Supplementary Section A.3.9. Additional modifications to the search space to allow for other copy number profiles is described in Supplementary Section A.3.7.

#### A.3.7 Modifying the search space of copy number profiles

To guide our reinforcement learning, our model is restricted to generating copy number profiles in some set of given copy number profiles (in addition to modifications of those copy number profiles by removing CNAs). NaiveCopy provides this set of initial copy number profiles. However, for the sake of improving the search, we add additional copy number profiles to the set of profiles that are allowed to be generated. During any iteration, for each cell *s*, the copy number profile that maximizes Pr(*P* | *θ*) Pr(*R*_*s*_, *B*_*s*_ | *P*) is calculated. We refer to this as the best-fit copy number profile for cell *s* in iteration *t*. For the next iteration, this copy number profile is included in the list of copy number profiles to be generated. Specifically, this changes the list of copy number profiles to be generated if this best-fit profile *P* was originally generated by removing CNAs from some copy number profiles originally in the list of profiles to be generated (described in Supplementary Section A.3.6). Additionally, for each best-fit copy number profile, a new copy number profile is generated by randomly changing the copy numbers for one chromosome to the copy numbers of some other best-fit copy number profile. These copy number profiles are also added to the list of profiles to be generated in the next iteration. This increases exploration during training. Additionally, another set of modified copy number profiles is added in order to avoid local minima in the cases of certain bins having very low haplotype-specific read counts. Specifically, the set of best fit profiles for each cell is copied, and then their haplotype-specific copy numbers are modified to better fit their haplotype-specific counts. To do so, for each bin and each copy number, the algorithm identifies all cells with that given copy number in that bin. The haplotype-specific read count of all of those cells in that bin is then calculated giving values *C*^*A*^, *C*^*B*^. Then, cell specific copy number (*X*^*A*^, *X*^*B*^) is modified to best fit *C*^*A*^, *C*^*B*^ while retaining the same total copy number. Let *B*^tweak^ = 0.9(*X*^*B*^*/*(*X*^*A*^ + *X*^*B*^)) + 0.05 be the BAF implied by the copy numbers (*X*^*A*^, *X*^*B*^) with a slight adjustment to avoid 0 or 1 probabilities. The algorithm then selects haplotype specific copy numbers and maximizes the log probability *C*^*A*^ log(1 *− B*^tweak^) + *C*^*B*^ log(*B*^tweak^). For many bins, this procedure will likely not modify the copy number profile. However, in cases with extremely low haplotype-specific read counts, the profiles may be improved due to this procedure of pooling counts together across cells. Note that these modified profiles are simply added to the set of possible profiles during training and are not required to be used by DeepCopy.

#### A.3.8 Efficient generative sequences

We previously mentioned the concept of sets of generative sequences that efficiently generate a copy number profile. This concept is defined precisely here. First, we define *B*_*E*_(*P*′, *P, c*) as a function that inputs two copy number profiles *P*′, *P*, and one CNA tuple *c*, and outputs either 0 or 1. We will define *B*_*E*_(*P*′, *P, c*) to output 1 if *c* efficiently modifies *P*′ towards *P*, and 0 otherwise. At first, we will ignore the existence of whole genome duplications, and then describe the modification to allow whole genome duplications.

Intuitively, define 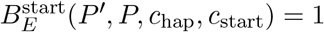 if *c*_start_ is the start position of a region of constant copy number in 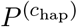 and 0 otherwise. Mathematically, we have

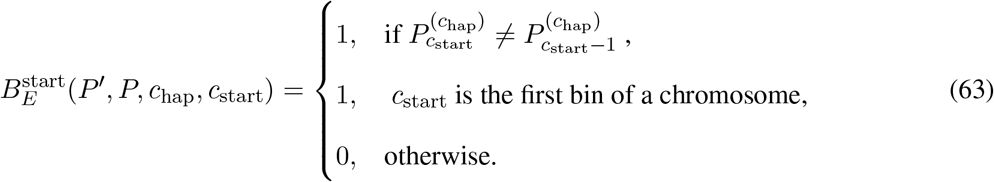

Similarly, we define 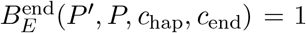 if *c*_end_ is the end of a region of constant copy number in 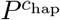 and 0 otherwise. Mathematically, we have

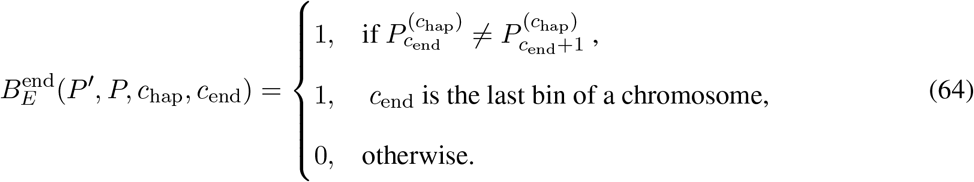

These two functions help ensure that our CNA tuple has a *c*_start_ at the start and *c*_end_ at the end for the CNA in *P*. Intuitively, let 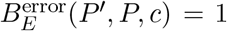 if the copy number aberration *c* results in *P*′ being more similar to *P*. Define Dist(*P, P*′) as the L1 distance between any two copy number profiles *P* and *P*′. As defined in Supplementary Section A.3.1, let *f*_*g*_(*P, G*) be a function that inputs a copy number profile *P* and a list *G* of CNA tuples and applies those CNAs to the profile. Mathematically, we have

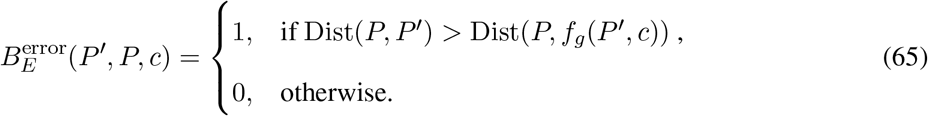

This function ensures that adding the CNA *c* to *P*′ results in a copy number profile that is closer to *P*. Utilizing these we have the below equation

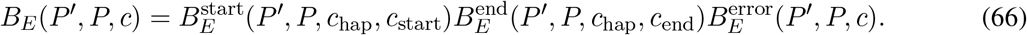

If the median total copy number in *P* is 4 or more then we ensure efficient generating sequences include a whole genome duplication (WGD). Specifically, if a whole genome duplication has not occurred yet, we replace *B*_*E*_(*P*′, *P, c*) with *B*_*E*_(*P*′, *P/*2, *c*), anticipating the duplication which will occur in *P*′. Additionally, we allow a whole genome duplication to occur once per cell in the generative process. Once the whole genome duplication has already occurred, then *B*_*E*_(*P*′, *P, c*) is utilized as normal (since the doubling of *P*′ has already occurred). Note that while we do not allow a WGD to occur multiple times in a cell’s evolutionary history (as modeled by *G*), we do allow distinct cells to undergo distinct WGDs in their evolutionary histories. Effectively, this supports the existence of potentially, multiple subclonal WGDs as well as a single truncal WGD.

Given our function *B*_*E*_(*P*′, *P, c*), we have the following constructive definition of the set *𝒜*^*∗*^(*P*) of generative sequences that efficiently generate *P* and its intermediates. Initially, let *𝒜*^*∗*^(*P*) contain the empty set {}. Inductively, define [*c*^1^, …, *c*^*k*^] ∈ *𝒜*^*∗*^(*P*) if [*c*^1^, …, *c*^*k−*1^] ∈ *𝒜*^*∗*^(*P*) and *B*_*E*_(*g*([*c*^1^, …, *c*^*k−*1^]), *P, c*^*k*^) = 1. Finally, define the set *𝒜*(*P*) of efficient generative sequences that generate exactly *P* as *𝒜*(*P*) = {*G* ∈ *𝒜*^*∗*^(*P*) | *g*(*G*) = *P* }.

#### A.3.9 Restricted sampling probabilities

In this section, we describe sampling restricted to *𝒜*(*P*). Let *𝒜*^*∗*^(*P*) be as defined in Supplementary Section A.3.8. First, define *B*(*P*, [*c*^1^, …, *c*^*k*^]) = {*c*^*k*+1^ | [*c*^1^, …, *c*^*k*+1^] ∈ 𝒜^*∗*^(*P*)}. In other words ℬ(*P*, [*c*^1^, …, *c*^*k*^]) defines the set of subsequent CNA tuples that can occur after the application of [*c*^1^, …, *c*^*k*^] to the normal profile *P*_*∅*_ in the process of efficiently generating *P*. This allows us to define the sampling probability of each new CNA tuple given existing CNA tuples as

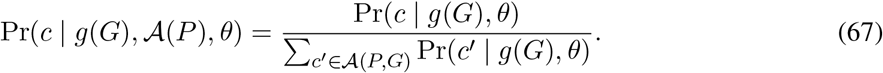

We recursively define a probability for any generating sequence [*c*^1^, …, *c*^*k*^] ∈ *𝒜*^*∗*^(*P*) as

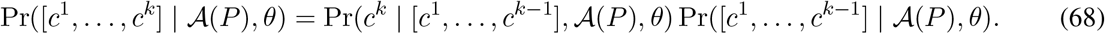

If *G* = [*c*^1^, …, *c*^*k*^] ∈ *𝒜*(*P*), then Pr(*G* | *𝒜*(*P*), *θ*) is the sampling probability of *G* (prior to the modification for removing fake CNAs).

In Section A.3.6 we state that the sampling procedure is modified to help remove fake CNAs. Specifically, we allow the model to predict any [*c*^1^, …, *c*^*k*^] with [*c*^1^, …, *c*^*k*^, …, *c*^*k*+*r*^] ∈ *𝒜*, where 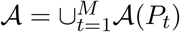 for preliminary copy number profiles *P*_1_, …, *P*_*M*_ guiding the optimization. In practice, our implementation first samples some [*c*^1^, …, *c*^*r*^] ∈ *𝒜* (P) (for some *P* in our set of allowed copy number profiles) and then automatically samples all [*c*^1^, …, *c*^*k*^] with *k ≤ r*. The probability of such a generating sequence [*c*^1^, …, *c*^*k*^] is still proportional to Pr(*c*^1^, …, *c*^*k*^ | *𝒜* (*P*), *θ*) (but with some normalization constant).

### A.4 Post-processing and validation

#### A.4.1 Determining clone sizes when not all cells have values for all bins

The results of SIGNALS on both datasets and CHISEL on the ovarian cancer dataset do not contain copy number values for some cells in each bin. Instead, some bins may have no copy number value for some cells due to having no SNPs. As suggested by the SIGNALS team, for most of our comparisons we fill in these missing values using neighboring bins. However, for analyzing clone sizes as well as the number of unique copy number profiles, we allow two cells to be in the same clone as long as they agree on all copy numbers for all bins for which values are provided. Specifically, to determine the set of clones, we initialize the set of clones as empty and the set of cells to be put into clones as the full set of cells. Then, iteratively, if some cell does not match any existing clone (in terms of the copy numbers on bins for which there exist copy number values), a new clone is formed by that cell. If some cell does match an existing clone (in terms of the copy numbers on bins which have copy number values for both the cell and clone), then that cell is added to the clone. When a cell is added to a clone, the clone then has a value for any bin for which either the clone originally had a value, or the new cell being added had a value. This process is repeated until all cells are within clones. The number of unique copy number profiles is then equal to the number of clones. For SIGNALS predictions on both datasets, there exists no pair of cells that agree on the values of all bins for which both cells have values. Consequently, for SIGNALS, there exist no clones containing more than 1 cell, and each cell has a unique copy number profile.

#### A.4.2 Determining CNA trees

To calculate a tree on the set of cells, we calculate a CNA distance between the copy number profiles of pairs of cells and then apply neighbor-joining. Specifically, we slightly modify the zero-agnostic copy number transformation (ZCNT) distance to include whole genome duplication [34]. The original copy number transformation (CNT) distance from a haploid copy number profile *P* ∈ *ℕ*^*L*^ to copy number profile *P*′ ∈ *ℕ*^*L*^ is the minimum number of CNA events required to transform *P* into *P*′ [59]. The ZCNT provides a fast approximation on the CNT distance by allowing zero copy number regions to be amplified into non-zero copy number regions. Although this assumption is not realistic, the ZCNT distance closely approximates the CNT distance while being quick enough to calculate in order to easily calculate distances between millions of pairs of cells [34]. Additionally, the ZCNT distance is symmetric, such that the distance from *P* to *P*′ is equal to the distance from *P*′ to *P*, i.e., ZCNT(*P, P*′) = ZCNT(*P*′, *P*). To compute the distance between two diploid copy number profiles *P* = [*P* ^(1)^, *P* ^(2)^]^*⊤*^ and *Q* = [*Q*^(1)^, *Q*^(2)^]^*⊤*^, we use

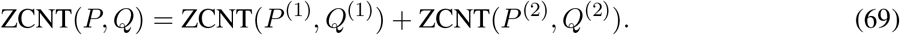

To allow for whole genome duplication, we define the distance 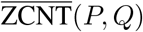 between copy number profiles *P* and *Q* as

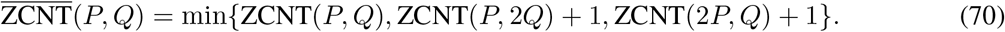

When calculating the distance matrix *D*, we also include one normal cell in addition to the original *N* cells in the dataset. We use the Neighbor Joining algorithm [52] to compute an unrooted tree from distance matrix *D*. We then root this tree at this normal cell, prior to removing the normal cell from the tree. After calculating the rooted tree *T*, we must determine CNA events that occur on the tree. To do so, we first apply the Sankoff algorithm [53] to solve the small parsimony problem, finding the copy number states of the internal nodes of the tree, one bin at a time. Note that although applying the Sankoff algorithm assumes independence of bins, CNA events should typically occur together within individual internal nodes due to the tree being constructed using ZCNT distances. Once all internal node copy number profiles are determined, we can calculate the parsimony score of the tree by again applying ZCNT distances. Specifically, the length of each edge is equal to the ZCNT distance, as defined in equation (69), between the copy number profiles of the nodes connected by the edge. Then, the parsimony score is the sum of distances on all edges.

##### Simplified visualization

While full trees for each method are shown in Figs. S16, S6, and S20, in the main text, we also showed a simplified visualization of DeepCopy’s tree on large clades that allows for labeling CNAs. To obtain this visualization, we set a minimum clade size to not be merged as min_merge_, and a minimum clade size to not be removed if unable to be merged as min_remove_. Then we initialize the set of *visualized clades* as the set of leaves. In some iteration, if two visualized clades are siblings, and either clade has below min_merge_ cells, they are merged into one visualized clade. This process is applied until there are no sibling visualized clades with a size below min_merge_ cells. However, there may be some extremely small visualized clades that are not siblings with visualized clades. For instance, a single-cell leaf may be siblings with a large part of the tree that contains many visualized clades. To address this, visualized clades with a size below min_remove_ are simply removed from the tree. In practice, this results in only a small portion of cells being removed from the tree while allowing for an effective visualization. This process results in an easily visualized tree in which are nodes are visualized clades with at least min(min_merge_, min_remove_) cells. In practice, we set min_remove_ *<* min_remove_ to try to minimize the number of cells removed. Additionally, in practice, most clades are at least as large as min_merge_. On the breast cancer patient S0, we set min_remove_ = 2, min_merge_ = 25, resulting in 781 out of 785 cells being kept in the simplified tree. On the ovarian cancer dataset [20], we set min_remove_ = 10, min_merge_ = 25, resulting in 573 out of 617 cells being kept in the simplified tree.

#### A.4.3 Calculating the probability of SNV counts given copy numbers

The goal of this section is to describe a method for orthogonally validating CNAs using single-nucleotide variants (SNVs) that were present in the *most recent common ancestor* (MRCA) of all tumor cells. Such SNVs are also known as *truncal*, as they occur on the trunk of the tumor phylogeny transforming a normal cell into the MRCA. As stated in the main text, we determine an SNV to be truncal if it occurred on at least 4 of 5 sections on breast cancer patient S0, or if it occurred on all three samples of the ovarian cancer patient.

We distinguish two types of truncal SNVs. The first type is a truncal SNV that occurred prior to any CNA affecting its genomic locus. In other words, such a truncal SNV was introduced in a cell with copy number (1, 1) at the SNV’s locus. Consequently, if subsequent CNAs occurred at that locus resulting in a final copy number of (*X*^(1)^, *X*^(2)^) in the MRCA then one expects the probability of variant reads and reference reads for that SNV to either be *X*^(1)^*/*(*X*^(1)^ + *X*^(2)^) or *X*^(2)^*/*(*X*^(1)^ + *X*^(2)^), depending on which allele the SNV occurred on. The second type is a truncal SNV that occurred after the introduction of CNAs affecting its genomic locus. Let (*X*^(1)^, *X*^(2)^) be the final copy number of the SNV’s locus in the MRCA. Then, the probability of variant reads and reference reads for that SNV is *x/*(*X*^(1)^ + *X*^(2)^) where *x* ∈ {1, …, *X*^(1)^ + *X*^(2)^}. As discussed in the main text, and shown in Supplementary Section B.1, the majority of truncal SNVs are of the first type in our data.

Let an SNV occur on some segment, and let the allele-specific copy number of that segment for cell *s* Be (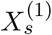, 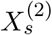) for the first and second allele, respectively. If one knew the SNV occurred on the first allele then one could estimate the probability of variant reads to be 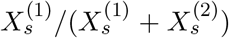, and the probability of reference reads to be 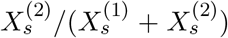. Let *v*_s_ be the number of variant reads for cell *s* and *r*_s_ be the number of reference reads for cell *s* observed at the SNV locus. Then, the probability of observing these reference reads and variant reads for cell *s* is the below equation assuming the SNV is on the A-allele

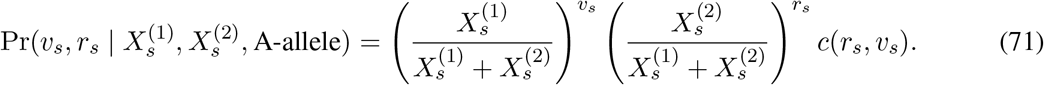

Here 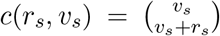 is the binomial coefficient only dependent on *r*_*s*_ and *v*_*s*_, but not the copy numbers of the cells. The probability of a set of variant reads and reference reads for all *N* cells for a given *SNV* then becomes the following

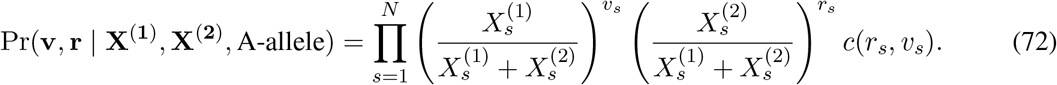

Similarly, if we assume the SNV occurred on the B-allele we have the following.

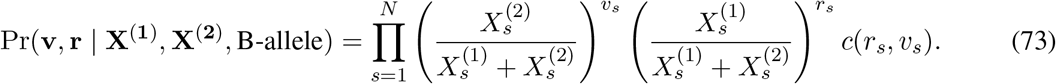

Since we do not know which allele the SNV occurred on, we take the maximum as below.

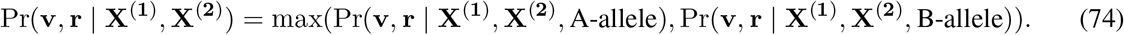

Taking the log probability ratio, i.e.,

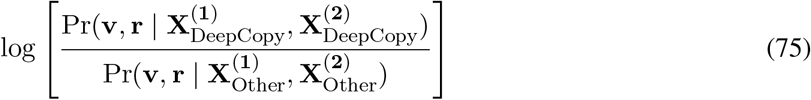

between our method and another method determines how much an SNV supports our method vs another method. Note that the binomial coefficients *c*(*r*_*s*_, *v*_*s*_) cancel out and do not need to be computed. Summing the log probability ratios across all truncal SNVs gives the total evidence for either method over the other, such that positive values indicate stronger support for our method versus the other method. We additionally perform bootstrapping on the set of SNVs to be summed, allowing us to obtain confidence intervals and quantify statistical significance.

However, one complication is that if a single read is said to come from an allele with copy number 0, i.e., either *X*^(1)^ = 0 or *X*^(2)^ = 0. In that case, all the probabilities go to 0 and the log probability ratio becomes infinite. To address this, we set a minimum probability of any reference of variant read to be 0.05 (and consequently the maximum probability to be 0.95) even if there is LOH.

#### A.4.4 Simulation set-up

The noise levels in the simulation are based on breast cancer patient S0. Consequently, the total number of 100kb bins and the number of 100kb bins in each chromosome is set to the same number *K* = 27,283 as in the real data. For each simulation, we define *p*_fit_ as the probability that each new clone will have increased fitness. Let *P*_1_ be the normal copy number profile with 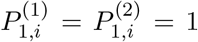 for all bins *i*. Alternatively in simulations with whole genome duplication, let 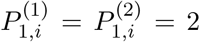 for all *i*. Define the fitness of this starting clone as 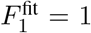. We then iterative define our procedure of adding CNAs to clones. Given the existing clones with profiles *P*_1_, …, *P*_*k*_ and fitness values 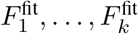, we first select the clone in which the CNA occurs. Specifically, we select clone *s* with probability 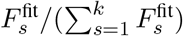. Let *a*_*k*+1_ be the index number of the selected clone. Since the copy (0, 0) occurs very rarely in cancer, we set the fitness 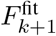 of the new clone to exp(*−*1000) if the new clone contains the copy number (0, 0). Otherwise, we randomly set 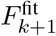 to either 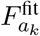 with probability 1 *− p*_fit_ or 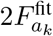 with probability *p*_fit_. The higher the probability *p*_fit_, the more likely the simulated tumor is to be dominated by a small set of large high fitness clones. With low values of *p*_fit_, most cells tend to come from distinct small clones. We varied *p*_fit_ ∈ {1*/*6, 1*/*8, 1*/*10, 1*/*12, 1*/*14, 1*/*16, 1*/*20, 1*/*50, 1*/*100, 1*/*1000} in our simulations, with lower values of *p*_fit_ resulting in more clonal expansion and thus larger clones and fewer unique copy number profiles. We initialize 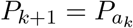 prior to applying the CNA. The chromosome that the CNA occurs on is selected with a probability proportional to the size of the chromosome. The parental haplotype on which the CNA occurs is selected uniformly at random. In breast cancer patient S0, it is common for CNAs to start or end at the start or end of a chromosome. Consequently, in our simulation, the start position has a 50% probability of being automatically set to the beginning of the chromosome. If the starting position is not automatically set to the beginning of the chromosome, the start position is instead selected uniformly at random within the chromosome. The end position has a 50% probability of being automatically set at the end of the chromosome. If the end position is not automatically set to the end of the chromosome, it is randomly selected from bins within the chromosome at or after the start position bin.

Let *c*_hap_, *c*_start_, *c*_end_ be the haplotype number selected, the starting position, and the ending position, respectively. The CNA is randomly set to be either a deletion or an amplification, each with 50% probability. If it is an amplification, we set 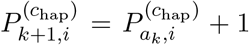 for all *i* with *c*_start_ *≤ i ≤ c*_end_. Let ReLU be the ReLU function, such that ReLU(*x*) = *x* for *x ≥* 0 and ReLU(*x*) = 0 for *x <* 0. If it is a deletion, set 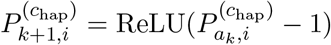 for all *i* with *c*_start_ *≤ i ≤ c*_end_ to prevent negative copy numbers. With this, we have generated 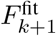 and *P*_*k*+1_ from *P*_1_, …, *P*_*k*_ and 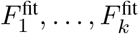. We continue this procedure for 4000 CNAs and thus generate *P*_1_, …, *P*_4001_ and 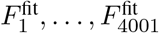.

After the copy number profiles have been generated, we generate *n* = 1000 cells and their read depth and BAF measurements. Specifically, each cell is assigned to clone *s* with probability 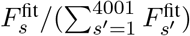. The noise-free read depth for a cell with copy number profile (*P* ^(1)^, *P* ^(2)^) is then defined below where it is scaled to have an average value of 1.

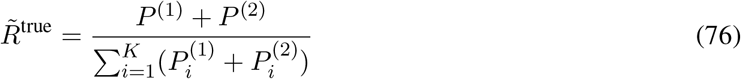

The noise-free BAF is defined as 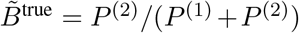, where the division is calculated element wise.

To generate the measured read depth and BAF, we must add noise. We do this based on the noise observed in breast cancer patient S0. For the BAF, the number 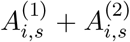 of observed haplotype specific reads for each bin *i* in each cell *s* is set to the average number of reads in the corresponding bin *i* in breast cancer patient S0. Specifically, the number of reads for each of the two alleles is drawn from a binomial distribution, with the probability of B-allele reads set to the true 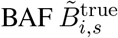, and the total number of binomial trials set to the total number 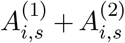 of haplotype specific reads. Consequently, some bins have a higher level of noise as a result of having fewer haplotype specific reads, which occurs in real data due to differing numbers of SNPs in different bins of the genome.

Simulating noise in the read depth is made slightly more complex by the fact that there is known to be overdispersion relative to what one would expect given a simple Poisson distribution. Specifically, if one assumes the reads are drawn from a Poisson distribution with the number of reads proportional to the total copy number then the simulated noise levels would be extremely low and unrealistic. Instead, we model the noise level based on breast cancer patient S0 noise levels directly. However, properly estimating which variations in read depth are due to noise also requires removing variations in read depth due to changes in copy number. To avoid this complexity, we estimate noise levels utilizing normal cells. Specifically, section A in breast cancer patient S0 consists primarily of normal cells. After filtering out 10% of cells (221 cells) with read depths that appear to be non-normal, we are left with 1970 normal cells. As previously mentioned, 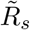 is scaled to have an average value of 1. Consequently, for normal cells 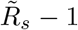 isolates changes in read depth due to noise. We calculate 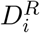 as the standard deviation in bin *i* as below.

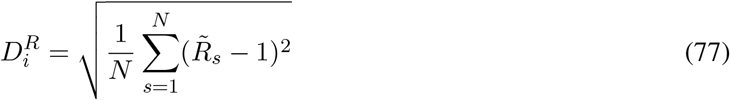

The average per bin per cell read count for the breast cancer patient S0 *R*_avg_ is then calculated. Then, for each bin *i* and each cell *s*, we use a negative binomial distribution with mean 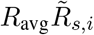and standard deviation 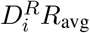. Since the number of successes parameter is an integer, we technically can not exactly match a given mean and standard deviation, so instead, we exactly match the mean and pick the parameters that closest match the desired standard deviation. This distribution then allows us to generate measured read depths 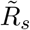 for each cell.

## B Supplementary results

### B.1 VAF analysis on additional copy numbers

As stated in the main text, we use the set notation {*X*^*A*^, *X*^*B*^} to refer to either of the copy numbers (*X*^*A*^, *X*^*B*^) or (*X*^*B*^, *X*^*A*^). In the main text, we plotted the variant allele frequency (VAF) of truncal SNVs that occur on cells with the copy number {2, 1} inferred by DeepCopy and Alleloscope for ovarian cancer patient OV2295 (Fig. 3f-g). Here, we provide similar VAF plots for other copy numbers inferred by Deep-Copy, SIGNALS, CHISEL, and Alleloscope. Note for VAF plots, to avoid excessive noise, we restrict to SNVs with at least 20 total reference and variant reads. For ovarian cancer patient OV2295, Fig. S7 and Fig. S10 shows VAF plots for DeepCopy, SIGNALS, CHISEL, and Alleloscope on several copy numbers. Specifically, all copy numbers with at least 300 SNVs for all methods are shown, in addition to the copy number {2, 2} which has at least 1,000 SNVs for all methods other than Alleloscope. For copy number {1, 2}, Alleloscope’s VAFs form a unimodal distribution concentrated around 1*/*2, in contrast to all other methods predicting the expected bimodal distribution with peaks around 1*/*3 and 2*/*3. Additionally, for {1, 1} and {1, 2}, Alleloscope’s VAFs have a large spike at 1.0, indicating LOH incorrectly being predicted as {1, 1} and {1, 2}. CHISEL also has a much smaller spike in the VAF at 1.0 for {1, 2}, whereas DeepCopy and SIGNALS do not have this issue. For the copy number {2, 2}, DeepCopy, SIGNALS, and CHISEL all have a unimodal distribution centered at 1*/*2, rather than having peaks at 1*/*4 or 3*/*4. This indicates SNVs occurring prior to CNAs or WGD. Additionally, for all methods, copy number {1, 2} is predicted to have allelic mirroring in which some cells have copy number (1, 2) whereas other cells have copy number (2, 1) for the same genomic region. For any SNV on haplotype A, the VAFs on copy numbers (1, 2) and (2, 1) are expected to be near 1*/*3 and 2*/*3 respectively as shown in Fig. S8. Similarly, for any SNV on haplotype B, the VAFs on copy numbers (1, 2) and (2, 1) are expected to be near 2*/*3 and 1*/*3 respectively. However, in either case, the VAFs on copy numbers (1, 2) and (2, 1) should not be both near 1*/*3 nor should they be both near 2*/*3. Fig. S9 shows the VAFs of SNVs (with at least 10 reads) on (1, 2) and (2, 1) for DeepCopy, SIGNALS, CHISEL, and Alleloscope, demonstrating agreement with SNVs for DeepCopy, SIGNALS, and CHISEL, but not Alleloscope. Specifically, the Pearson correlation between the VAF for (1, 2) and (2, 1) is *−*0.65, *−*0.72, *−*0.67, and *−*0.05 for DeepCopy, SIGNALS, CHISEL, and Alleloscope respectively, where a strong negative correlation validates predicted allelic mirroring.

In addition to these common copy numbers, we investigate a wide variety of additional copy numbers predicted by DeepCopy. In order to maximize the number of observed SNVs, we use all *n* = 890 cells, rather than the subset of *n* = 617 cells for which all methods have predictions. Fig. S10 shows VAFs for the top-11 copy numbers with the most SNVs (or equivalently, all copy numbers with at least 140 SNVs occurring on that copy number), as well as the copy number {3, 3} (which has 64 SNVs but is still very visually clear due to the simple unimodal VAF structure). For all copy numbers {*X*^*A*^, *X*^*B*^} there are peaks at *X*^*A*^*/*(*X*^*A*^ + *X*^*B*^) and *X*^*B*^*/*(*X*^*A*^ + *X*^*B*^) as expected.

On breast cancer patient S0, we show VAF plots for the *n* = 3,540 cells for which DeepCopy, SIGNALS, and CHISEL give predictions. We exclude Alleloscope since further restricting to *n* = 785 cells in one section would substantially reduce the SNV read counts making VAF plots much less visually clear. Fig. S17 shows VAF histograms for copy numbers whose corresponding segments contain at least 28 truncal SNVs for DeepCopy, SIGNALS, and CHISEL. For all methods and all copy numbers {*X*^*A*^, *X*^*B*^}, there are peaks at *X*^*A*^*/*(*X*^*A*^ + *X*^*B*^) and *X*^*B*^*/*(*X*^*A*^ + *X*^*B*^) as expected. For copy number 1, 2, CHISEL seems to have a less clear bimodal distribution than DeepCopy and SIGNALS, as a result of having more VAFs near 1*/*2 and more VAFs near 1.0. Additionally, DeepCopy, SIGNALS, and CHISEL all predict allelic mirroring for copy number {1, 2}. As previously described, SNV validation of allelic mirroring would result in the VAFs for (1, 2) and (2, 1) containing one value near 1*/*3 and one value near 2*/*3 (but not both values near 1*/*3 nor both values near 2*/*3). This pattern validating the predicted allelic mirroring is clear for DeepCopy and SIGNALS but not CHISEL as shown in Fig. S18. Specifically, the Pearson correlation between the VAF on (1, 2) and (2, 1) is *−*0.89, *−*0.89, and *−*0.20 for DeepCopy, SIGNALS, and CHISEL respectively, where a strong negative correlation demonstrates SNV validation of allelic mirroring. This result is in line with our statistical analysis on SNVs showing that CHISEL matches SNV data worse than DeepCopy and SIGNALS (Section 2.4).

### B.2 Detection of small CNAs

In Section 2.2, we analyzed the general accuracy of DeepCopy and NaiveCopy on simulated data. However, here we specifically analyze the accuracy of these methods in detecting small focal CNAs. These small CNAs can easily be overwhelmed by noise, causing copy number calling to be difficult to accomplish. Let 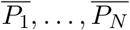 be the ground truth copy number profiles. We define the length of a CNA segment as the number of 100kb bins for which the copy number remains constant. Mathematically, let 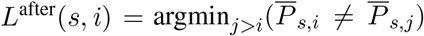. Let 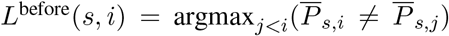. The length of the copy number segment containing *i* in cell *s* is *L*^length^(*s, i*) = *L*^after^(*s, i*) *− L*^before^(*s, i*) *−* 1.

We now define the set of bins in small CNAs, medium CNAs, and large CNAs. For simulation number *ζ ζ*, let 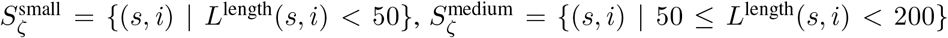, and 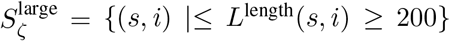. Then 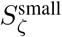 contains bins in CNAs smaller than 5Mb, 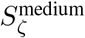 contains bins in CNAs between 5Mb and 20Mb, and 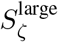 contains bins in CNAs at least 20Mb. As defined in Section 4.4, *δ*_L1_(*a, b, a*′, *b*′) = | max(*a, b*) *−* max(*a*′, *b*′)| + | min(*a, b*) *−* min(*a*′, *b*′)|. Then, pooling across all 20 simulations gives the following unordered allele-specific error measurement on small CNAs.

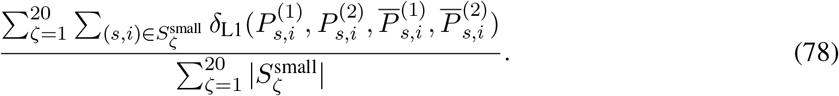

Identical equations but with “small” replaced by “medium” and “large” give the error on medium and large CNAs respectively.

As shown in Fig. S3a, DeepCopy achieves an unordered allele-specific copy number error of 0.61, 0.27, and 0.084 on small, medium, and large CNAs respectively. In contrast, SIGNALS has a much higher error of 1.25, 0.97, and 0.42 on small, medium and large CNAs respectively. CHISEL has a larger error than DeepCopy on medium and large CNAs (0.54 and 0.24 respectively) but a similar error on small CNAs (0.62).

Haplotype-specific error measurements are calculated as follows.

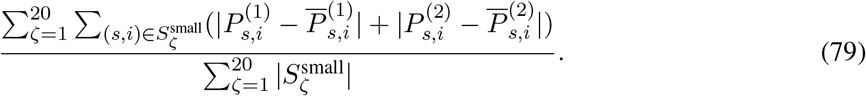

However, since the existing methods SIGNALS and CHISEL modify the phasing of their inputs, evaluating them directly on this haplotype-specific error is unfair. Consequently, we post-process the outputs of both of these existing methods to match the ground truth phasing before comparisons. This gives a lower bound of their haplotype-specific error. Specifically, to accomplish this, for each bin *i* we swap the values of 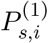and 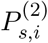 for all cells *s* if that reduces the L1 error. DeepCopy’s predictions do not receive this modification or any information from the ground truth copy number profiles. As shown in Fig. S3b, DeepCopy achieves the lowest errors for small, medium, and large CNAs (0.62, 0.27, and 0.084 respectively), whereas CHISEL has larger errors (0.68, 0.59 and 0.28 respectively) and SIGNALS has the highest errors (1.27, 0.98 and 0.43 respectively).

**Figure S1:**
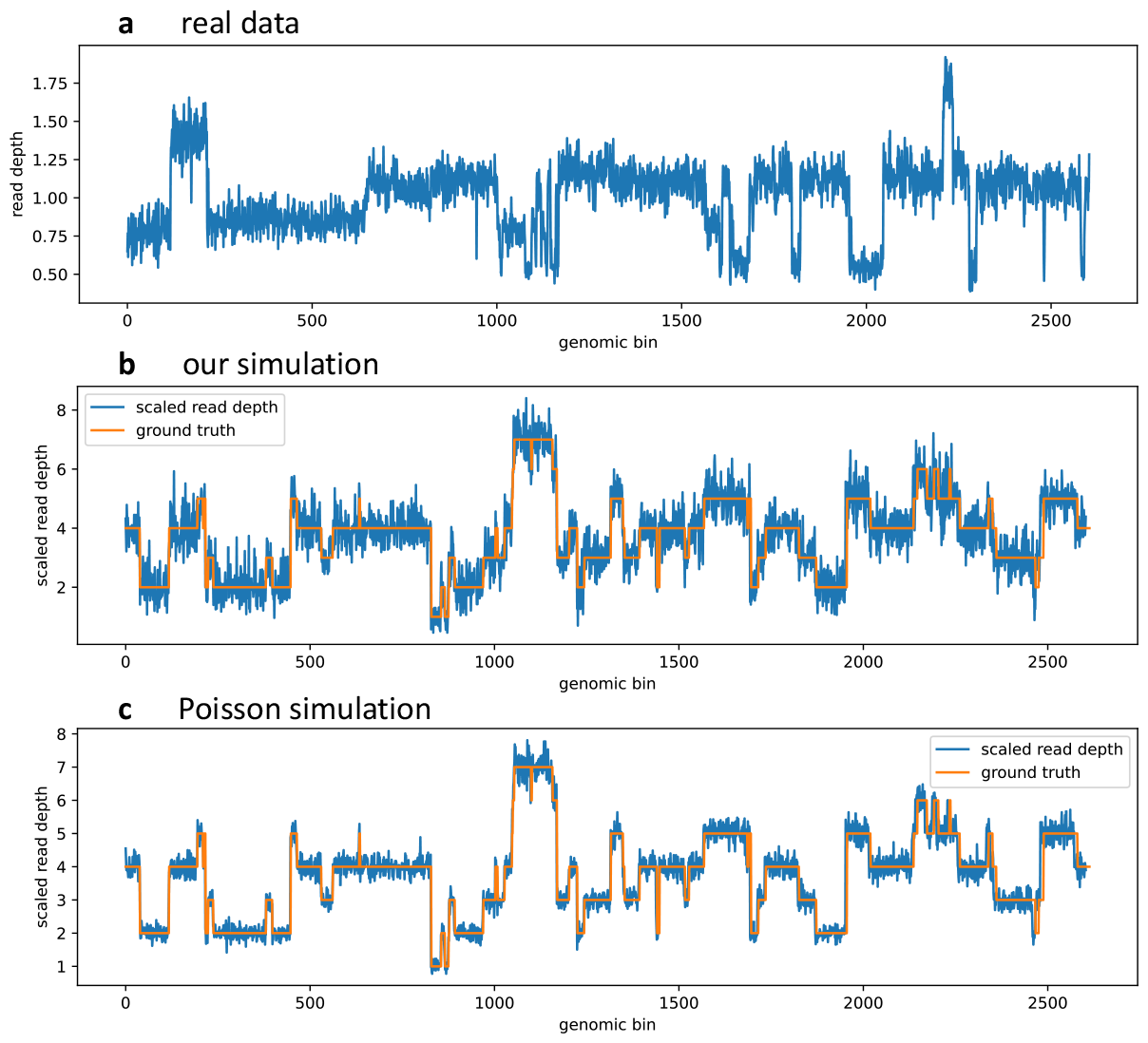
A comparison of breast cancer patient S0 read depths and our simulated read depths. Read depths on real and simulated data are shown. Bins of size 1Mb are used for the sake of visual clarity. **a**, Read depths for a cell in breast cancer patient S0 are shown. These read depths are scaled to have an average value of 1. **b**, Read depths are shown for one cell in our simulation instance. The read depths are scaled such that the average read depth equals the average copy number to allow an easy comparison with the ground truth. **c**, A Poission noise-based read depth plot on the same simulated cell is generated. The number of reads used is equal to the average number of reads per cell breast cancer patient S0 (989,469 reads). These reads are then mapped to bins with a probability proportional to the total copy number of the bin. The read depths are scaled such that the average read depth equals the average copy number to allow an easy comparison with the ground truth. Despite using the same number of reads as in breast cancer patient S0, the noise level is still clearly far below the noise level of the measured read depths from breast cancer patient S0.

**Figure S2:**
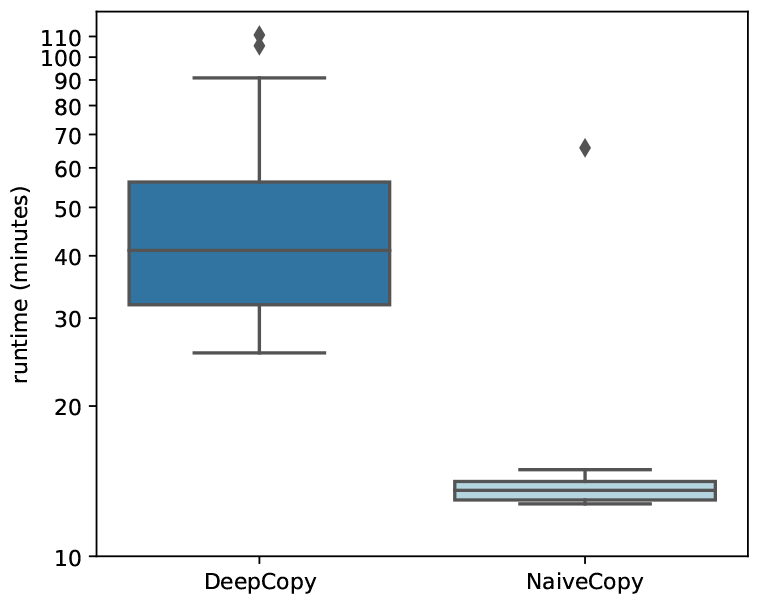
Runtimes on 20 simulation instances for DeepCopy and NaiveCopy. NaiveCopy steps starting from read depths 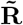 and haplotype-specific counts **A** (provided by the simulation) run fairly quickly. The runtime of DeepCopy’s optimization ranges from 26 minutes to 111 minutes, whereas NaiveCopy’s runtime ranges from 13 minutes to 66 minutes. DeepCopy (and NaiveCopy) were run on a laptop with 96GB of RAM and a 3.6 GHz processor (with 12 cores), without the use of a GPU for all experiments.

**Figure S3:**
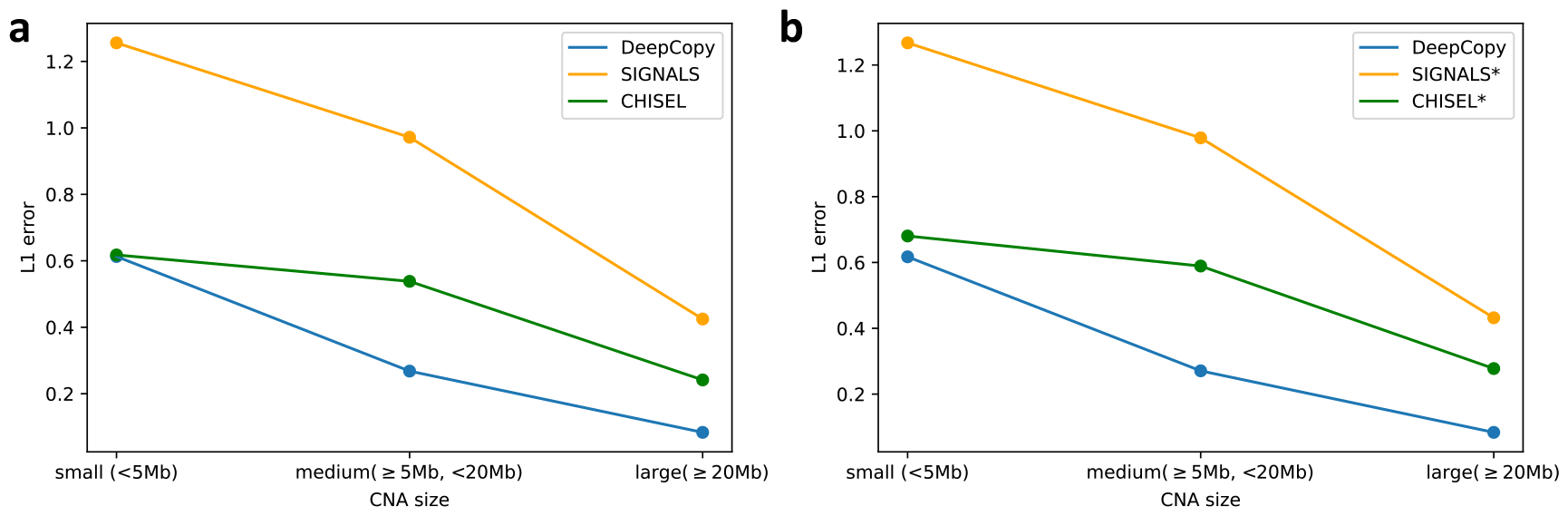
The error of DeepCopy and NaiveCopy on small, medium, and large CNAs on simulated data. **a**, Unordered allele-specific L1 error (equation (79)). **b**, Haplotype-specific L1 error. SIGNALS and CHISEL are given ground truth phasing.

**Figure S4:**
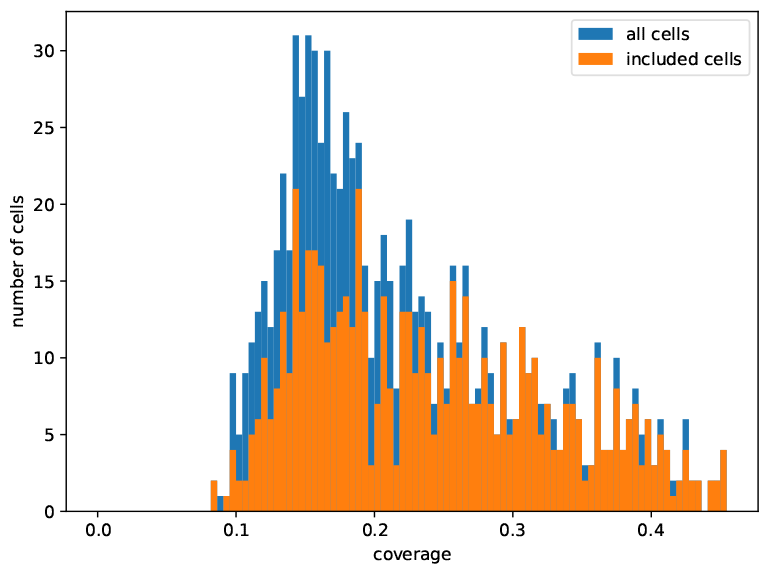
The coverage of cells of ovarian cancer patient OV2295 [20]. A histogram of the coverage of cells. In blue is the set of all 890 available cells. In orange are the *n* = 617 cells for which all methods were run on and comparisons were performed.

**Figure S5:**
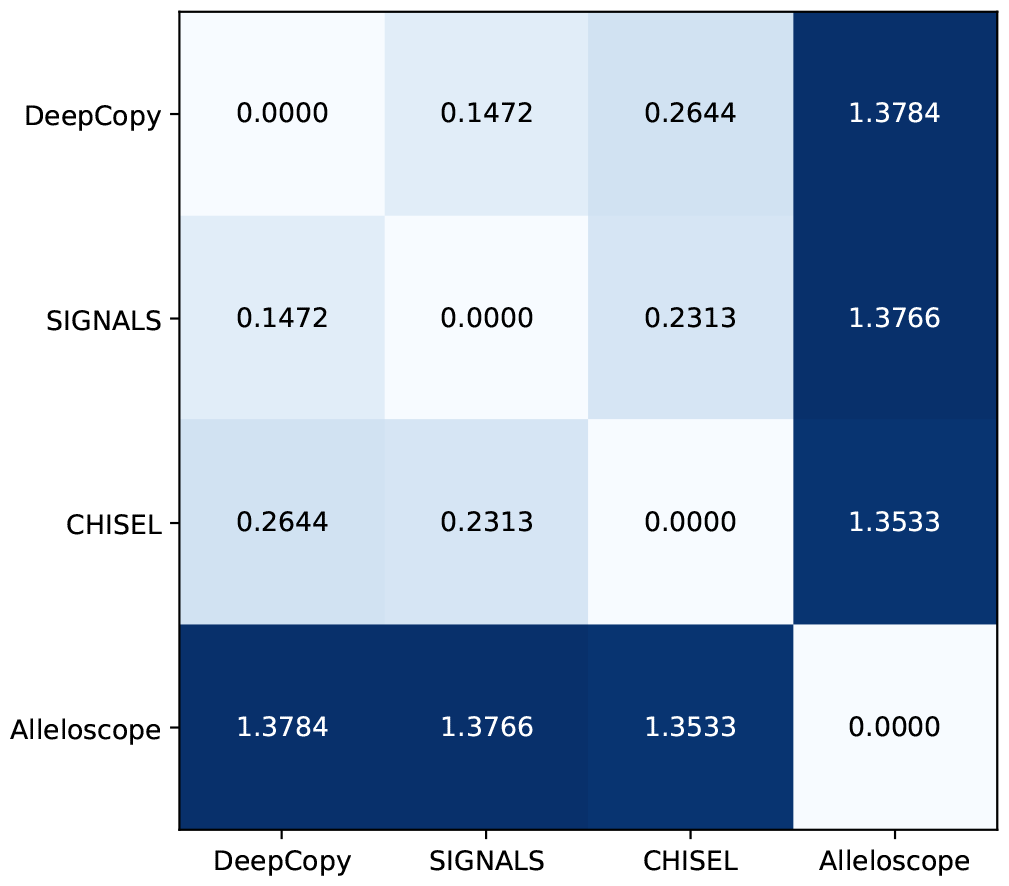
L1 differences between copy number profiles predicted by DeepCopy, SIGNALS, CHISEL, and Alleloscope on ovarian cancer patient OV2295 [20]. Haplotype-specific copy number profiles are compared for DeepCopy, SIGNALS, CHISEL and Alleloscope on *n* = 617 cells from ovarian cancer patient OV2295 [20]. Distances are calculated as in equation 79, with each bin phased to minimize errors between methods (as was done when comparing SIGNALS and CHISEL to ground truth copy numbers in Supplementary Section B.2). Alleloscope is by far the most distant from other methods, with a distance over 1.35 to any other method while all other methods’ predictions are within a distance of 0.265 of each other. This is due to the lack of whole genome duplication in Alleloscopes predictions.

**Figure S6:**
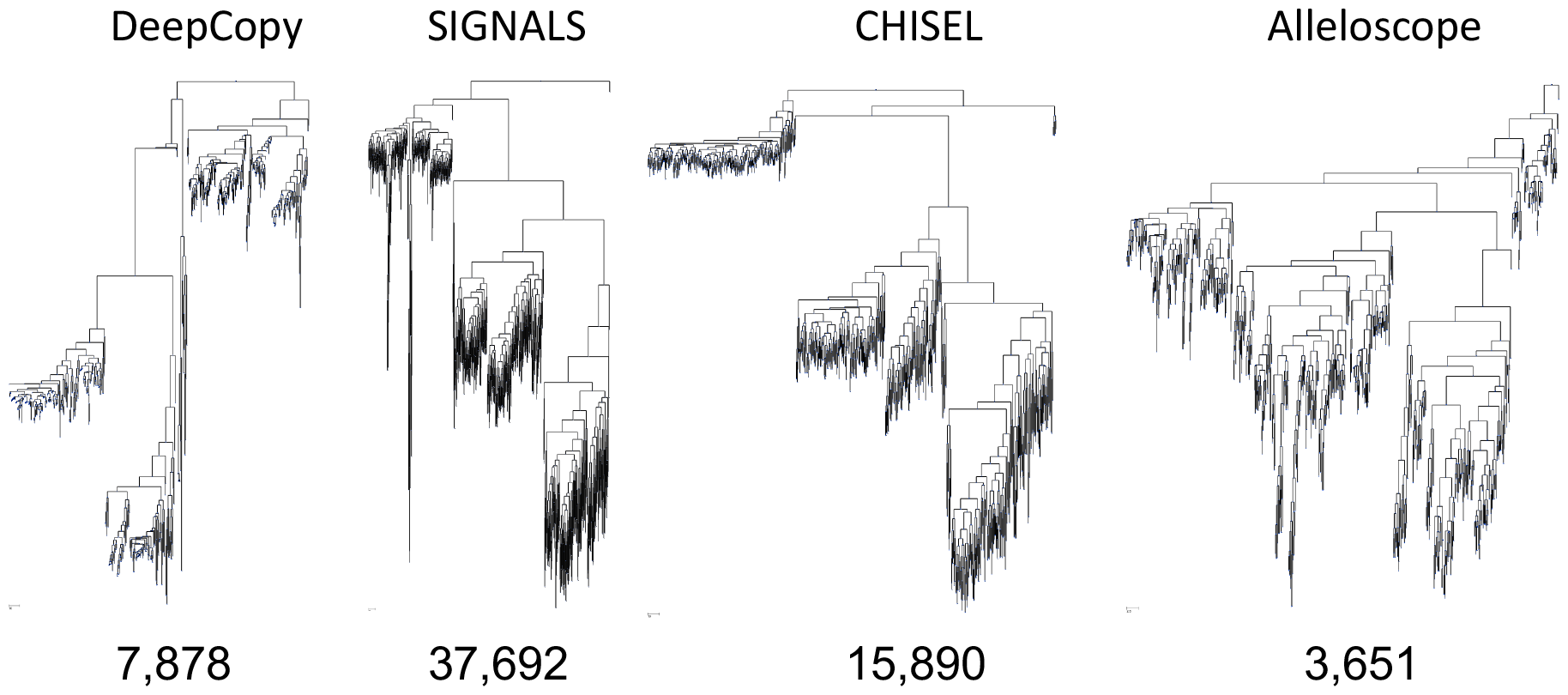
Full trees of each method for ovarian cancer patient OV2295 [20]. Full trees on the set of unique copy number profiles (clones) for DeepCopy, SIGNALS, CHISEL, and Alleloscope. These trees are scaled to the same height for plotting, however, the relative length of branches in each plot is proportional to the number of CNA events on the branch (estimated by the ZCNT distance). Parsimony values are labeled below each tree. Branches near the leaves are the shortest for DeepCopy and Alleloscope, the longest for SIGNALS, and have intermediate lengths for CHISEL. Inferring spurious CNAs results in CNAs that do not follow an evolutionary tree and consequently occur near leaves (possibly with homoplasy). Additionally, Alleloscope missing legitimate differences in whole genome duplication results in a very low parsimony score.

**Figure S7:**
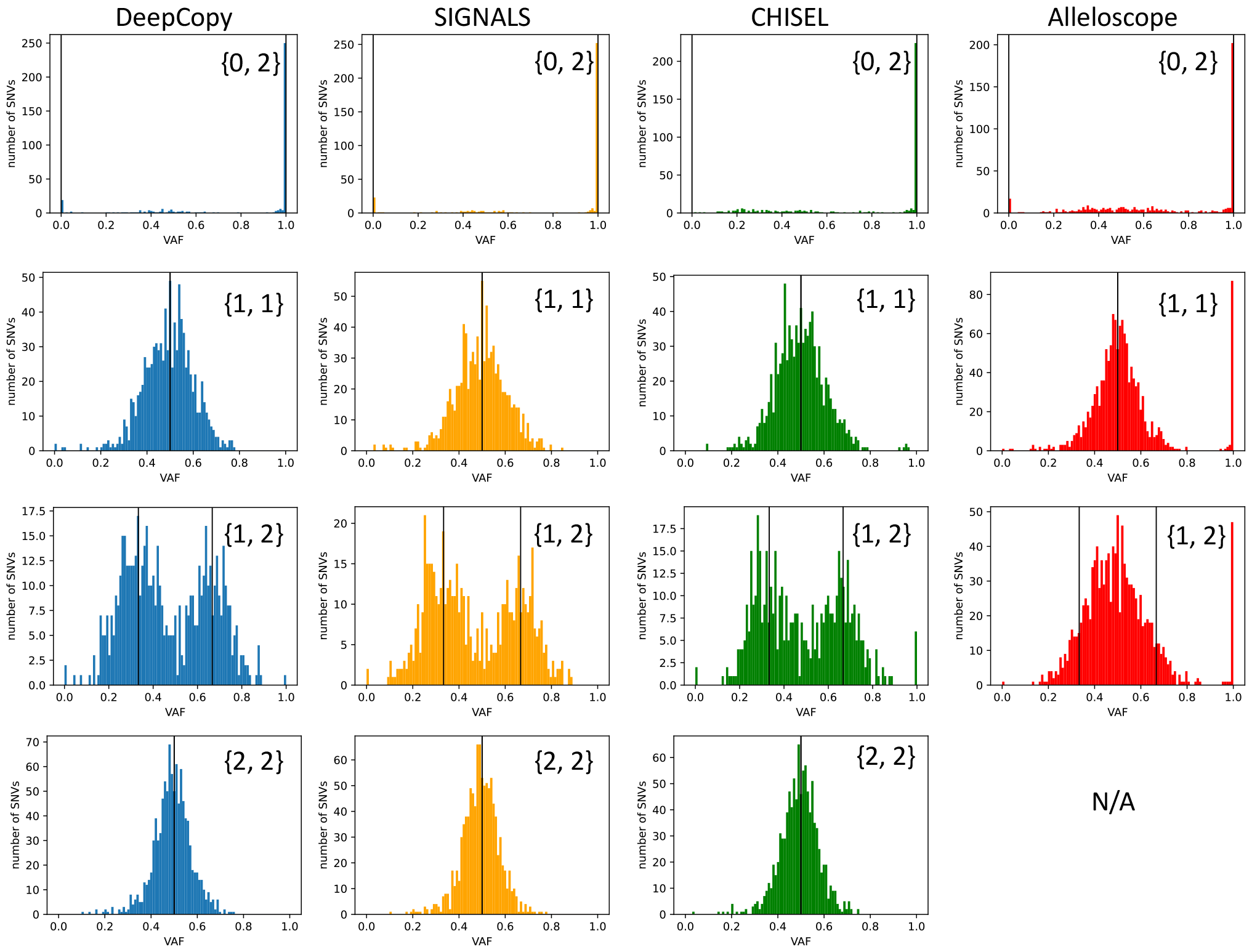
VAF values for common copy numbers for ovarian cancer patient OV2295 [20]. The common copy numbers {0, 2}, {1, 1}, and {1, 2} are shown for all DeepCopy, SIGNALS, CHISEL, and Alleloscope. The copy number {2, 2} is shown for DeepCopy, SIGNALS, and CHISEL, but not Alleloscope (due to the rarity of this copy number according to Alleloscope’s predictions). With the exception of Alleloscope, all VAF plots have peaks around the theoretically expected values as indicated with black vertical lines.

**Figure S8:**
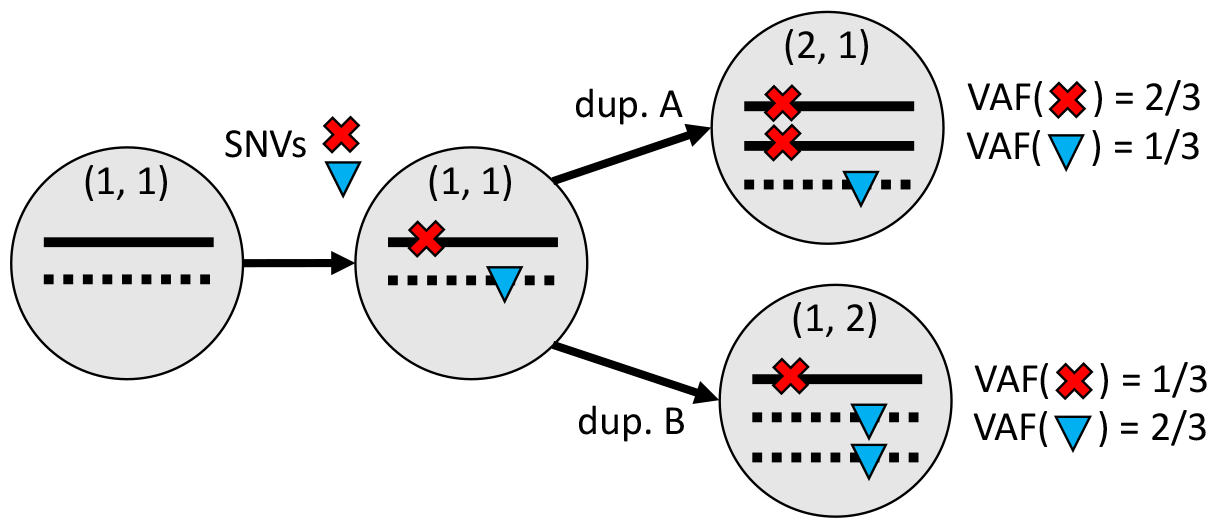
A diagram of how allelic mirroring affects VAFs. The impact of allelic mirroring on the VAF of truncal SNVs is shown. Specifically, the red SNV occurs on haplotype A (solid) and thus has an expected VAF of 1*/*3 on copy number (1, 2) and an expected VAF of 2*/*3 on copy number (2, 1). Similarly, the blue SNV occurs on haplotype B (dashed) and thus has an expected VAF of 2*/*3 on copy number (1, 2) and an expected VAF of 1*/*3 on copy number (2, 1).

**Figure S9:**
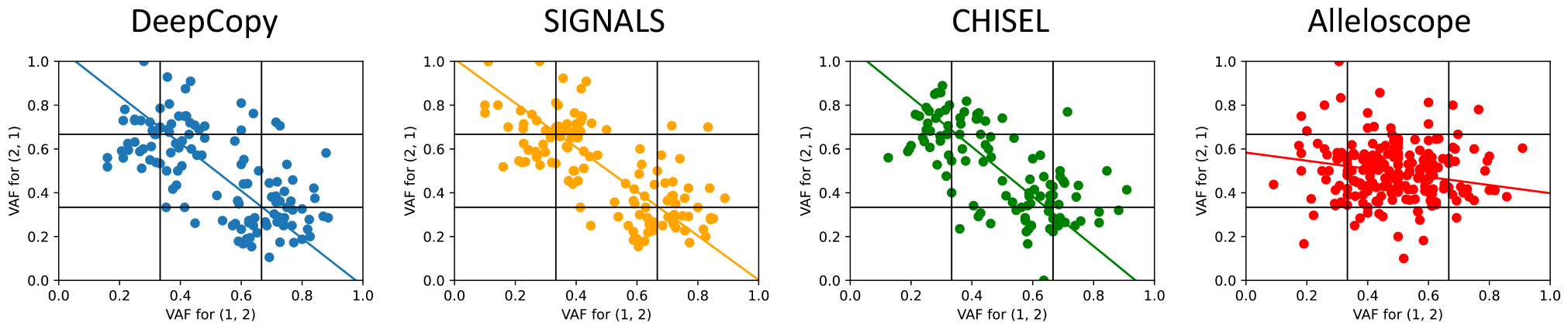
VAFs demonstrating allelic mirroring for copy number {2, 1} on ovarian cancer patient OV2295 [20]. VAFs are shown for SNVs that occur on both copy number (1, 2) and (2, 1) due to allelic mirroring. A best fit line for each method is shown where orthogonal distance regression is used due to their being noise in both *x* and *y* variable measurements. A strong negative Pearson correlation between VAFs on (1, 2) and (2, 1) demonstrates SNV support of predicted allelic mirroring for DeepCopy (*r* = *−*0.65), SIGNALS (*r* = *−*0.72) and CHISEL (*r* = *−*0.67) but not Alleloscope (*r* = *−*0.05).

**Figure S10:**
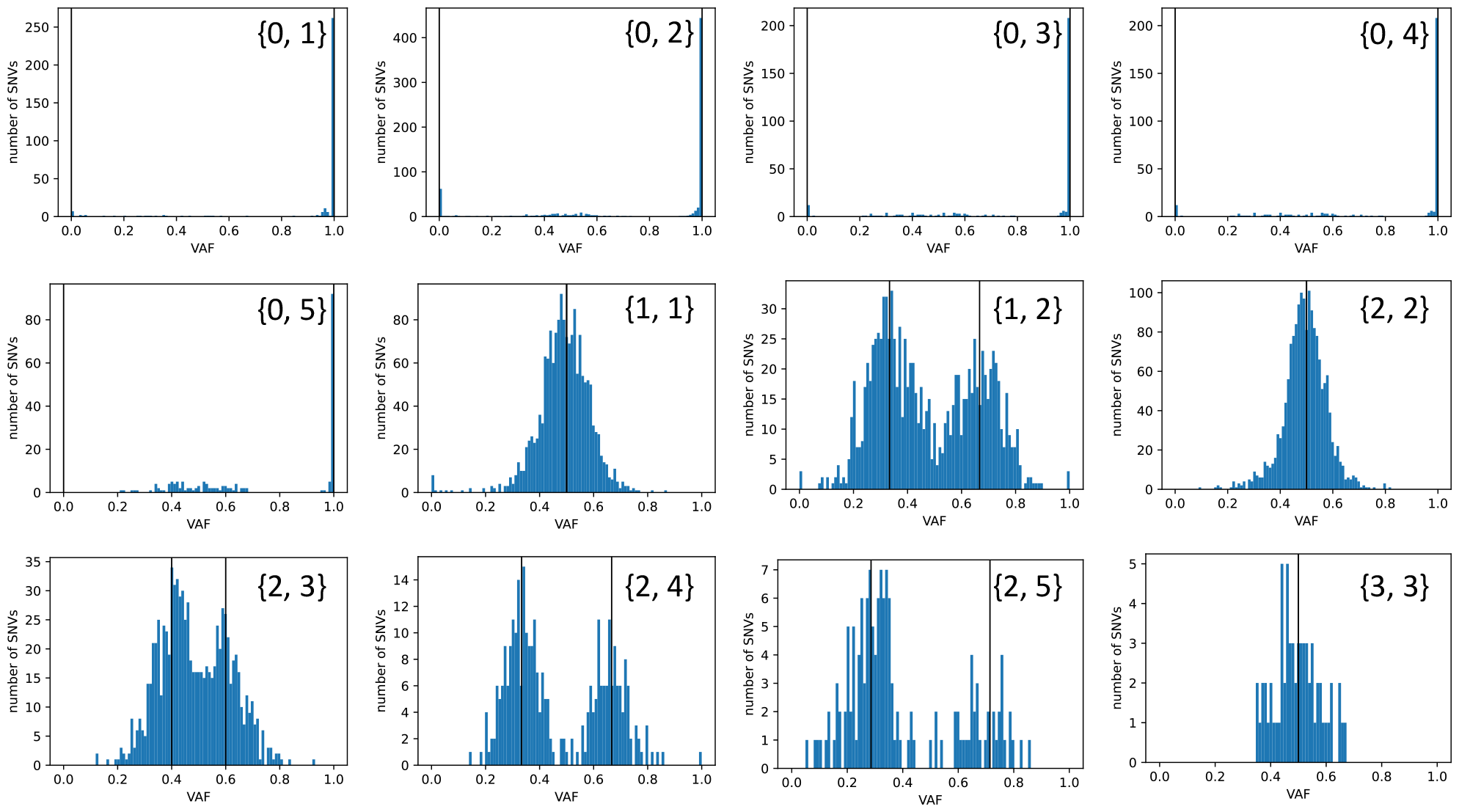
VAFs of truncal SNVs overlapping with various copy numbers inferred by DeepCopy on ovarian cancer patient OV2295 [20]. VAFs are shown for all the top-11 copy numbers with the most SNVs (or equivalently, all copy numbers with at least 140 SNVs occurring on that copy number), as well as the copy number {3, 3} (which has 64 SNVs but is still very visually clear due to the simple unimodal VAF structure). The allele-specific copy number of each plot is indicated on the plot. Black vertical lines indicate the expected VAF of SNVs either occurring on the major or minor allele. On all plots, the VAFs are concentrated around these expected numbers.

**Figure S11:**
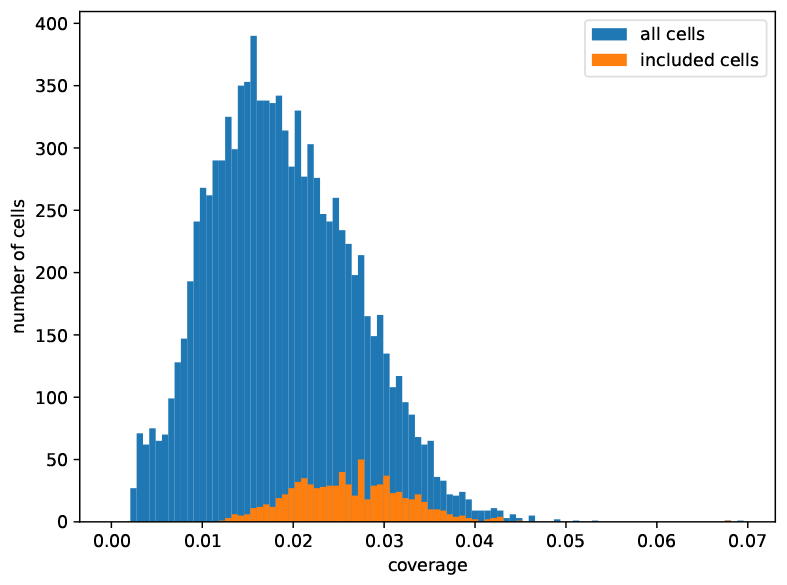
The coverage of cells of breast cancer patient S0. A histogram of the coverage of cells. In blue is the set of all 10,202 available cells. In orange are the *n* = 785 cells for which all methods were run on and comparisons were performed. Note that many of the excluded cells were normal (non-cancerous) cells, which will have a lower coverage due to the absence of amplifications and whole genome duplications.

**Figure S12:**
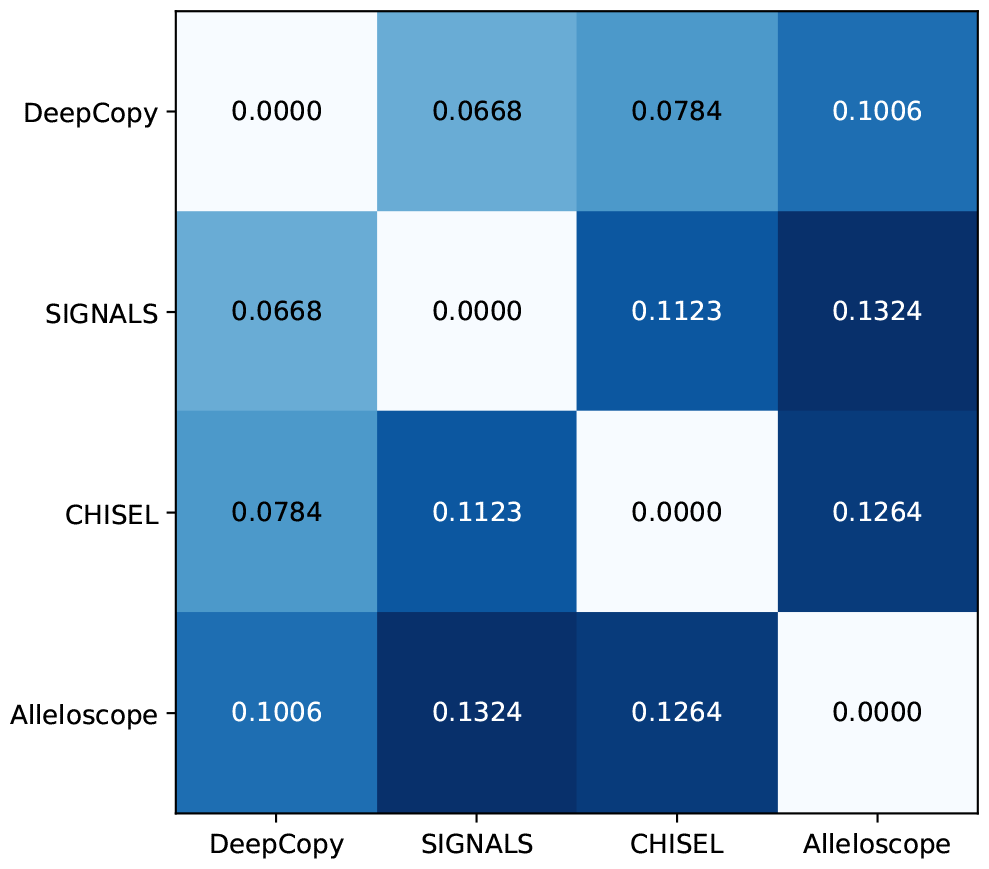
L1 differences between copy number profiles predicted by DeepCopy, SIGNALS, CHISEL, and Alleloscope on breast cancer patient S0. Haplotype-specific copy number profiles are compared for DeepCopy, SIGNALS, CHISEL, and Alleloscope on *n* = 785 cells from breast cancer patient S0. Distances are calculated as in equation (79), with each bin phased to minimize errors between methods (as was done when comparing SIGNALS and CHISEL to ground truth copy numbers in Supplementary Section B.2). For all methods, the minimum distance alternative method prediction is DeepCopy, indicating that DeepCopy best fits the consensus of other methods.

**Figure S13:**
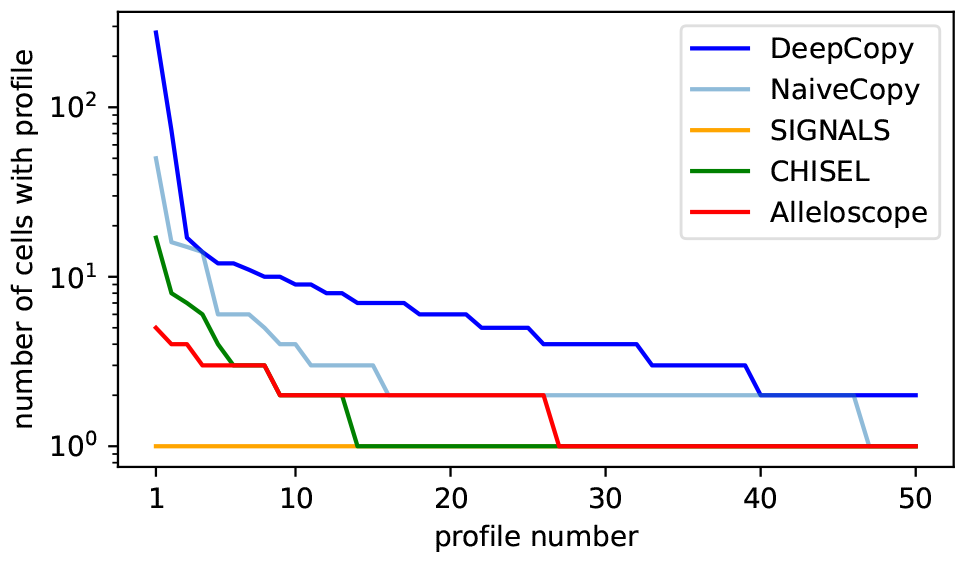
The number of cells for NaiveCopy for common copy number profiles on breast cancer patient S0. The number of cells for the top 50 most common copy number profiles on breast cancer patient S0 is shown for DeepCopy, NaiveCopy, SIGNALS, CHISEL, and Allelscope. Specifically, NaiveCopy is now included to show the specific advantage of DeepCopy’s evolutionary model. The number of cells in the most common copy number profile is 276, 50, 1, 17, and 5 for DeepCopy, NaiveCopy, SIGNALS, CHISEL, and Alleloscope, respectively. The number of unique copy number profiles is 206, 628 785, 737, and 747 for DeepCopy, NaiveCopy, SIGNALS, CHISEL, and Alleloscope, respectively.

**Figure S14:**
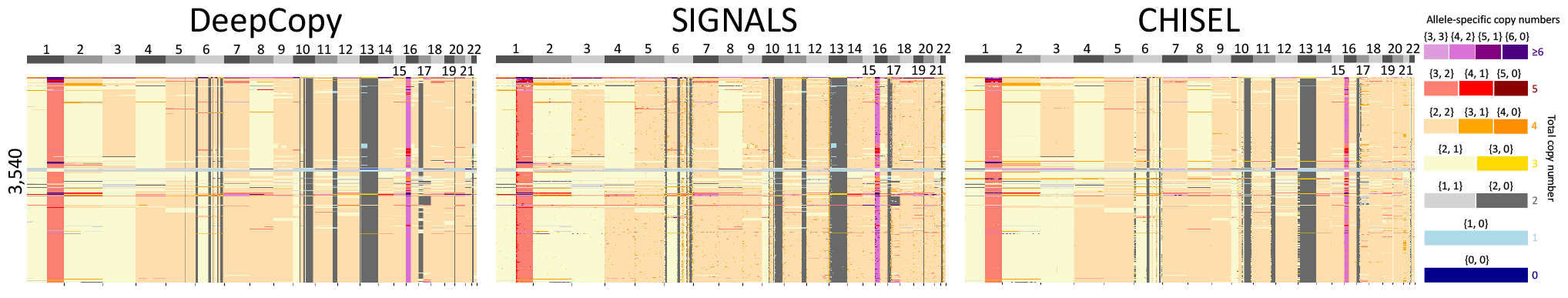
A heatmap of allele-specific copy number predictions on 3,540 cells from breast cancer patient S0. Here we show allele-specific copy number predictions on the full set of 3,540 cells for which DeepCopy, SIGNALS, and CHISEL have predicted profiles.

**Figure S15:**
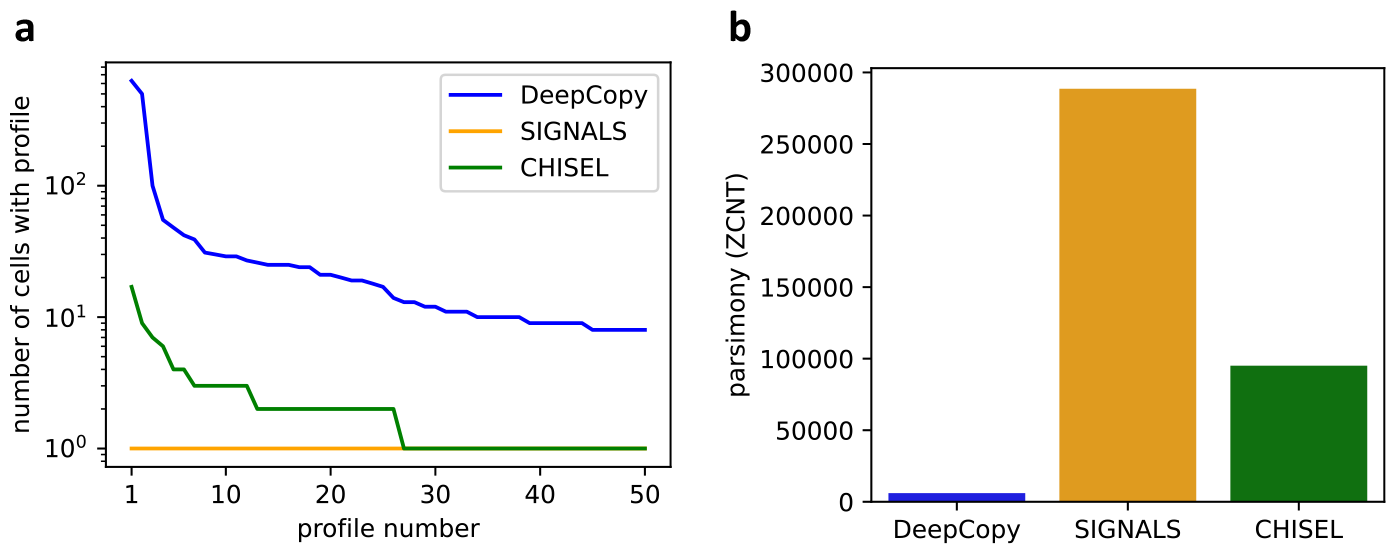
Clone sizes and parsimony on on 3,540 cells from breast cancer patient S0. **a**, A comparison of clone sizes for DeepCopy, CHISEL, and SIGNALS. The most common unique copy number profiles have 629, 1, and 17 cells for DeepCopy, CHISEL, and SIGNALS respectively. **b**, A comparison of parsimony scores for DeepCopy, CHISEL, and SIGNALS, showing parsimony scores of 6,014, 288,568, and 95,084, respectively.

**Figure S16:**
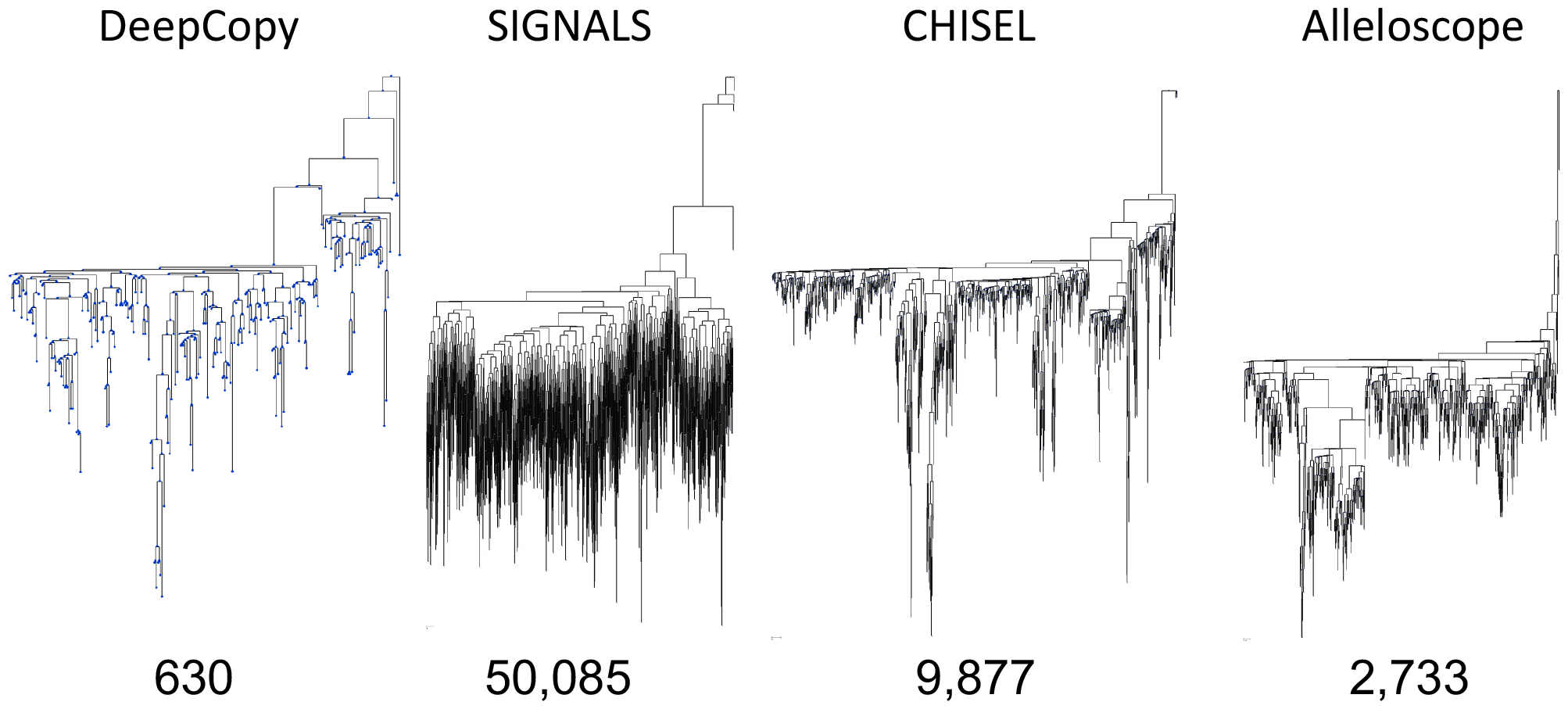
Full trees of each method on breast cancer patient S0. Full trees on the set of unique copy number profiles (clones) for DeepCopy, SIGNALS, CHISEL, and NaiveCopy. All trees are scaled to the same height for plotting, however, the relative length of branches in each plot is proportional to the number of CNA events on the branch (estimated by the ZCNT distance). Parsimony values are labeled next to each tree. Branches near the leaves are the shortest for DeepCopy, longer for CHISEL and Alleloscope, and extremely long for SIGNALS. Inferring spurious CNAs results in CNAs that do not follow an evolutionary tree and consequently occur near leaves (possibly with homoplasy), resulting in very long branches near the leaves.

**Figure S17:**
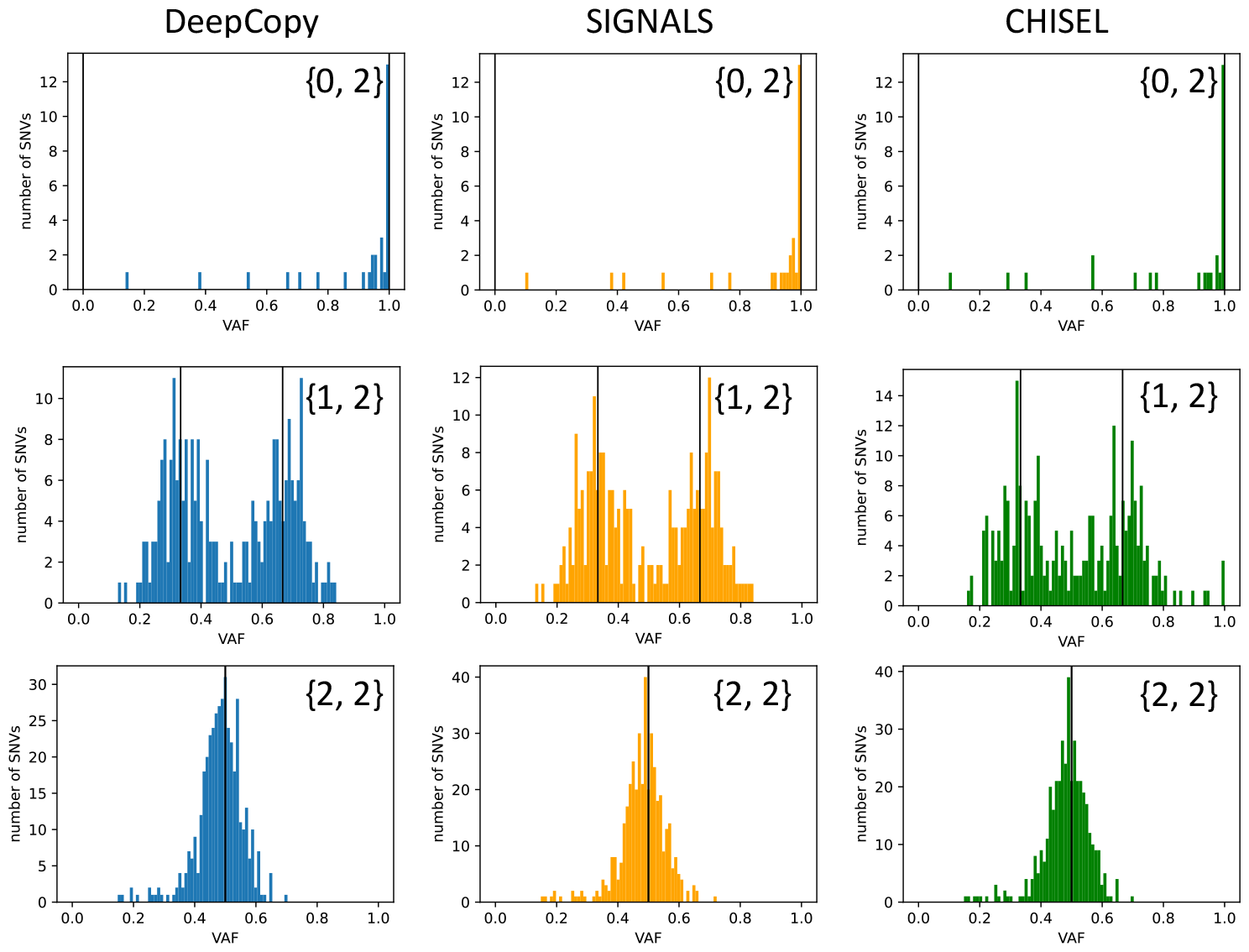
VAFs of truncal SNVs overlapping with three common copy numbers for DeepCopy SIGNALS and CHISEL on breast cancer patient S0. VAFs are shown for the top-3 copy numbers with the most SNVs (or equivalently, all copy numbers with at least 28 SNVs occurring on that copy number for DeepCopy, CHISEL, and SIGNALS). The allele-specific copy number of each plot is indicated on the plot. Black vertical lines indicate the expected VAF of SNVs either occurring on the major or minor allele. The VAFs are concentrated around these expected numbers.

**Figure S18:**
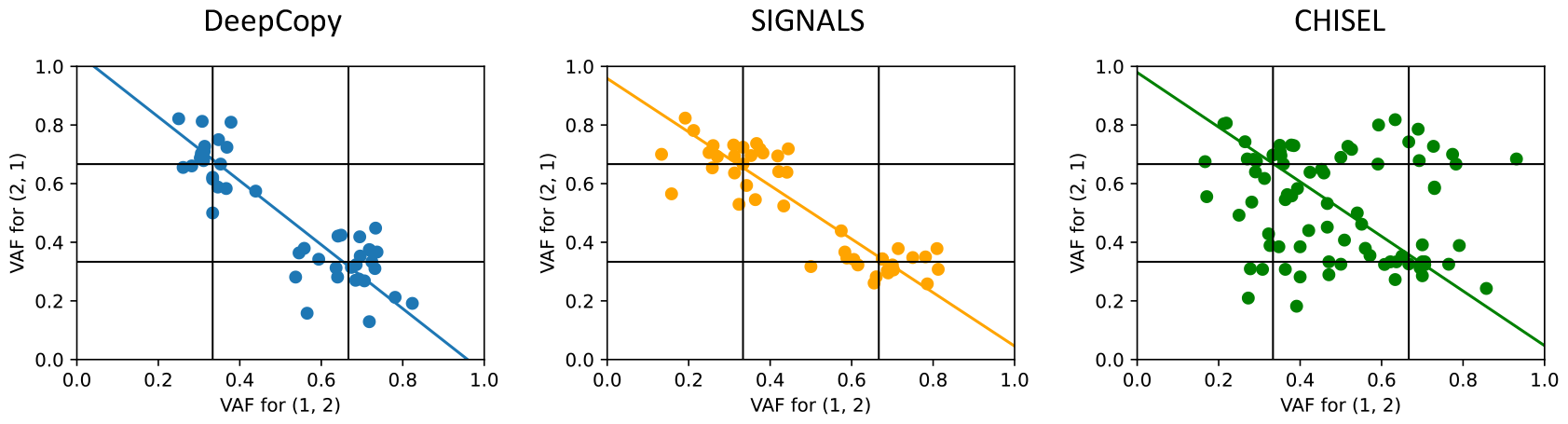
VAFs demonstrating allelic mirroring for copy number {2, 1} on breast cancer patient S0. VAFs are shown for SNVs that occur on both copy number (1, 2) and (2, 1) due to allelic mirroring. A best fit line for each method is shown where orthogonal distance regression is used due to their being noise in both *x* and *y* variable measurements. A strong negative Pearson correlation between VAFs on (1, 2) and (2, 1) demonstrates SNV support of predicted allelic mirroring for DeepCopy (*r* = *−*0.89) and SIGNALS (*r* = *−*0.89), but not CHISEL (*r* = *−*0.20).

**Figure S19:**
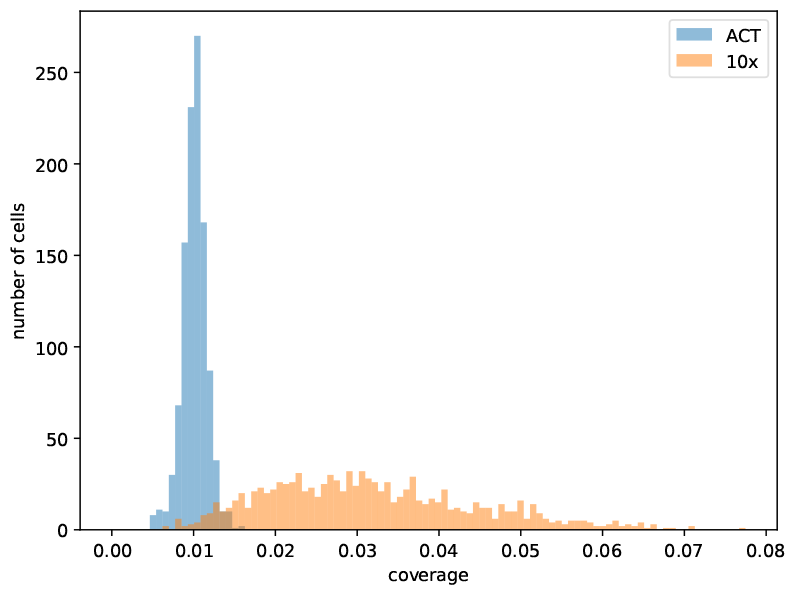
The coverage of cells of breast cancer patient TN3 [22]. The coverage of breast cancer patient TN3 cells, with blue representing cells sequenced with ACT technology and orange representing cells sequenced with 10x technology.

**Figure S20:**
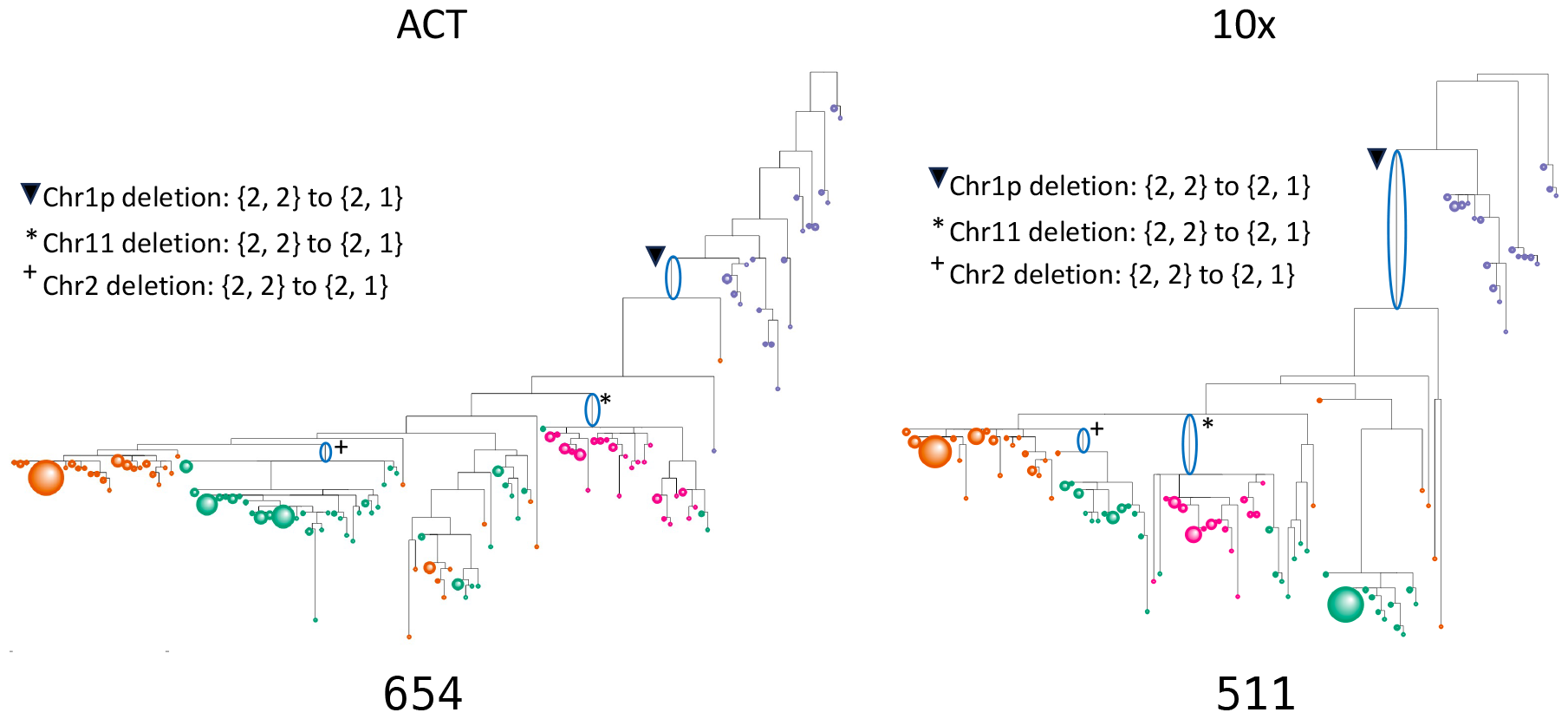
Phylogenies for breast cancer patient TN3 [22] for both sequencing technologies. Phylogenies are shown for patient TN3, with a node for each unique copy number profile. The sizes (area) of the nodes are proportional to the number of cells with that profile, and the color of the nodes matches the UMAP clusters (shown in Fig. 5a). As shown below the trees, the parsimony score is 654 on the cells sequenced with ACT technology and the parsimony is 511 on the trees sequenced with 10x technology. Note that parsimony scores can be strongly affected by focal CNAs on small numbers of cells, resulting in different parsimony values despite their being highly similar predicted copy number profiles. Three major events on the tree are indicated, showing deletions of entire chromosomes or chromosome arms.

## References

[1] Peter C Nowell. The clonal evolution of tumor cell populations: Acquired genetic lability permits stepwise selection of variant sublines and underlies tumor progression. Science, 194(4260):23–28, 1976.

[2] Thomas BK Watkins, Emilia L Lim, Marina Petkovic, Sergi Elizalde, Nicolai J Birkbak, Gareth A Wilson, David A Moore, Eva Grönroos, Andrew Rowan, Sally M Dewhurst, et al. Pervasive chromosomal instability and karyotype order in tumour evolution. Nature, 587(7832):126–132, 2020.

[3] Stefan C Dentro, Ignaty Leshchiner, Kerstin Haase, Maxime Tarabichi, Jeff Wintersinger, Amit G Deshwar, Kaixian Yu, Yulia Rubanova, Geoff Macintyre, Jonas Demeulemeester, et al. Characterizing genetic intra-tumor heterogeneity across 2,658 human cancer genomes. Cell, 184(8):2239–2254, 2021.

[4] Pan-cancer analysis of whole genomes. Nature, 578(7793):82–93, 2020.

[5] Yael Cohen-Sharir, James M McFarland, Mai Abdusamad, Carolyn Marquis, Sara V Bernhard, Mariya Kazachkova, Helen Tang, Marica R Ippolito, Kathrin Laue, Johanna Zerbib, et al. Aneuploidy renders cancer cells vulnerable to mitotic checkpoint inhibition. Nature, 590(7846):486–491, 2021.

[6] Ryan J Quinton, Amanda DiDomizio, Marc A Vittoria, Kristyna Kotynková, Carlos J Ticas, Sheena Patel, Yusuke Koga, Jasmine Vakhshoorzadeh, Nicole Hermance, Taruho S Kuroda, et al. Wholegenome doubling confers unique genetic vulnerabilities on tumour cells. Nature, 590(7846):492–497, 2021.

[7] Danish Memon, Michael B Gill, Evangelia K Papachristou, David Ochoa, Clive S D’Santos, Martin L Miller, and Pedro Beltrao. Copy number aberrations drive kinase rewiring, leading to genetic vulnerabilities in cancer. Cell reports, 35(7), 2021.

[8] Peter Van Loo, Silje H Nordgard, Ole Christian Lingjærde, Hege G Russnes, Inga H Rye, Wei Sun, Victor J Weigman, Peter Marynen, Anders Zetterberg, Bjørn Naume, et al. Allele-specific copy number analysis of tumors. Proceedings of the National Academy of Sciences, 107(39):16910–16915, 2010.

[9] Ruibin Xi, Angela G Hadjipanayis, Lovelace J Luquette, Tae-Min Kim, Eunjung Lee, Jianhua Zhang, Mark D Johnson, Donna M Muzny, David A Wheeler, Richard A Gibbs, et al. Copy number variation detection in whole-genome sequencing data using the bayesian information criterion. Proceedings of the National Academy of Sciences, 108(46):E1128–E1136, 2011.

[10] Valentina Boeva, Tatiana Popova, Kevin Bleakley, Pierre Chiche, Julie Cappo, Gudrun Schleiermacher, Isabelle Janoueix-Lerosey, Olivier Delattre, and Emmanuel Barillot. Control-FREEC: a tool for assessing copy number and allelic content using next-generation sequencing data. Bioinformatics, 28(3):423–425, 2012.

[11] Gavin Ha, Andrew Roth, Jaswinder Khattra, Julie Ho, Damian Yap, Leah M Prentice, Nataliya Melnyk, Andrew McPherson, Ali Bashashati, Emma Laks, et al. TITAN: inference of copy number architectures in clonal cell populations from tumor whole-genome sequence data. Genome research, 24(11):1881–1893, 2014.

[12] Ronglai Shen and Venkatraman E Seshan. FACETS: allele-specific copy number and clonal heterogeneity analysis tool for high-throughput DNA sequencing. Nucleic acids research, 44(16):e131–e131, 2016.

[13] Faiyaz Notta, Michelle Chan-Seng-Yue, Mathieu Lemire, Yilong Li, Gavin W Wilson, Ashton A Connor, Robert E Denroche, Sheng-Ben Liang, Andrew MK Brown, Jaeseung C Kim, et al. A renewed model of pancreatic cancer evolution based on genomic rearrangement patterns. Nature, 538(7625):378–382, 2016.

[14] Simone Zaccaria and Benjamin J Raphael. Accurate quantification of copy-number aberrations and whole-genome duplications in multi-sample tumor sequencing data. Nature communications, 11(1):1–13, 2020.

[15] Teng Gao, Ruslan Soldatov, Hirak Sarkar, Adam Kurkiewicz, Evan Biederstedt, Po-Ru Loh, and Peter V Kharchenko. Haplotype-aware analysis of somatic copy number variations from single-cell transcriptomes. Nature Biotechnology, 41(3):417–426, 2023.

[16] Ruli Gao, Shanshan Bai, Ying C Henderson, Yiyun Lin, Aislyn Schalck, Yun Yan, Tapsi Kumar, Min Hu, Emi Sei, Alexander Davis, et al. Delineating copy number and clonal substructure in human tumors from single-cell transcriptomes. Nature biotechnology, 39(5):599–608, 2021.

[17] Jean Fan, Hae-Ock Lee, Soohyun Lee, Da-eun Ryu, Semin Lee, Catherine Xue, Seok Jin Kim, Kihyun Kim, Nikolaos Barkas, Peter J Park, et al. Linking transcriptional and genetic tumor heterogeneity through allele analysis of single-cell rna-seq data. Genome research, 28(8):1217–1227, 2018.

[18] Anoop P Patel, Itay Tirosh, John J Trombetta, Alex K Shalek, Shawn M Gillespie, Hiroaki Wakimoto, Daniel P Cahill, Brian V Nahed, William T Curry, Robert L Martuza, et al. Single-cell rna-seq highlights intratumoral heterogeneity in primary glioblastoma. Science, 344(6190):1396–1401, 2014.

[19] Noemi Andor, Billy T Lau, Claudia Catalanotti, Anuja Sathe, Matthew Kubit, Jiamin Chen, Cristina Blaj, Athena Cherry, Charles D Bangs, Susan M Grimes, Carlos J Suarez, and Hanlee P Ji. Joint single cell dna-seq and rna-seq of gastric cancer cell lines reveals rules of in vitro evolution. NAR Genomics and Bioinformatics, 2(2):qaa016, June 2020.

[20] Emma Laks, Andrew McPherson, Hans Zahn, Daniel Lai, Adi Steif, Jazmine Brimhall, Justina Biele, Beixi Wang, Tehmina Masud, Jerome Ting, et al. Clonal decomposition and DNA replication states defined by scaled single-cell genome sequencing. Cell, 179(5):1207–1221, 2019.

[21] Hans Zahn, Adi Steif, Emma Laks, Peter Eirew, Michael VanInsberghe, Sohrab P Shah, Samuel Aparicio, and Carl L Hansen. Scalable whole-genome single-cell library preparation without preamplification. Nature methods, 14(2):167–173, 2017.

[22] Darlan C Minussi, Michael D Nicholson, Hanghui Ye, Alexander Davis, Kaile Wang, Toby Baker, Maxime Tarabichi, Emi Sei, Haowei Du, Mashiat Rabbani, et al. Breast tumours maintain a reservoir of subclonal diversity during expansion. Nature, 592(7853):302–308, 2021.

[23] Nicholas Navin, Jude Kendall, Jennifer Troge, Peter Andrews, Linda Rodgers, Jeanne McIndoo, Kerry Cook, Asya Stepansky, Dan Levy, Diane Esposito, Lakshmi Muthuswamy, Alex Krasnitz, W. Richard McCombie, James Hicks, and Michael Wigler. Tumour evolution inferred by single-cell sequencing. Nature, 472(7341):90–94, April 2011.

[24] Sohrab P Shah, Xiang Xuan, Ron J DeLeeuw, Mehrnoush Khojasteh, Wan L Lam, Raymond Ng, and Kevin P Murphy. Integrating copy number polymorphisms into array cgh analysis using a robust hmm. Bioinformatics, 22(14):e431–e439, 2006.

[25] Tyler Garvin, Robert Aboukhalil, Jude Kendall, Timour Baslan, Gurinder S Atwal, James Hicks, Michael Wigler, and Michael C Schatz. Interactive analysis and assessment of single-cell copy-number variations. Nature methods, 12(11):1058–1060, 2015.

[26] Jack Kuipers, Mustafa Anil Tuncel, Pedro Ferreira, Katharina Jahn, and Niko Beerenwinkel. Single-cell copy number calling and event history reconstruction. BioRxiv, pages 2020–04, 2020.

[27] Xuefeng Wang, Hao Chen, and Nancy R Zhang. Dna copy number profiling using single-cell sequencing. Briefings in Bioinformatics, 19(5):731–736, September 2018.

[28] Rujin Wang, Dan-Yu Lin, and Yuchao Jiang. Scope: a normalization and copy-number estimation method for single-cell dna sequencing. Cell systems, 10(5):445–452, 2020.

[29] Simone Zaccaria and Benjamin J Raphael. Characterizing allele-and haplotype-specific copy numbers in single cells with CHISEL. Nature biotechnology, 39(2):207–214, 2021.

[30] Tyler Funnell, Ciara H O’Flanagan, Marc J Williams, Andrew McPherson, Steven McKinney, Farhia Kabeer, Hakwoo Lee, Sohrab Salehi, Ignacio Vázquez-García, Hongyu Shi, et al. Single-cell genomic variation induced by mutational processes in cancer. Nature, 612(7938):106–115, 2022.

[31] Chi-Yun Wu, Billy T Lau, Heon Seok Kim, Anuja Sathe, Susan M Grimes, Hanlee P Ji, and Nancy R Zhang. Integrative single-cell analysis of allele-specific copy number alterations and chromatin accessibility in cancer. Nature biotechnology, 39(10):1259–1269, 2021.

[32] Xian F Mallory, Mohammadamin Edrisi, Nicholas Navin, and Luay Nakhleh. Methods for copy number aberration detection from single-cell dna-sequencing data. Genome biology, 21(1):1–22, 2020.

[33] Jochen Singer, Jack Kuipers, Katharina Jahn, and Niko Beerenwinkel. Single-cell mutation identification via phylogenetic inference. Nature communications, 9(1):5144, 2018.

[34] Henri Schmidt, Palash Sashittal, and Benjamin J Raphael. A zero-agnostic model for copy number evolution in cancer. bioRxiv, pages 2023–04, 2023.

[35] Ann-Marie Patch, Elizabeth L Christie, Dariush Etemadmoghadam, Dale W Garsed, Joshy George, Sian Fereday, Katia Nones, Prue Cowin, Kathryn Alsop, Peter J Bailey, et al. Whole–genome characterization of chemoresistant ovarian cancer. Nature, 521(7553):489–494, 2015.

[36] Yilong Li, Nicola D Roberts, Jeremiah A Wala, Ofer Shapira, Steven E Schumacher, Kiran Kumar, Ekta Khurana, Sebastian Waszak, Jan O Korbel, James E Haber, et al. Patterns of somatic structural variation in human cancer genomes. Nature, 578(7793):112–121, 2020.

[37] Ruben M Drews, Barbara Hernando, Maxime Tarabichi, Kerstin Haase, Tom Lesluyes, Philip S Smith, Lena Morrill Gavarró, Dominique-Laurent Couturier, Lydia Liu, Michael Schneider, et al. A pancancer compendium of chromosomal instability. Nature, 606(7916):976–983, 2022.

[38] Yi Kan Wang, Ali Bashashati, Michael S Anglesio, Dawn R Cochrane, Diljot S Grewal, Gavin Ha, Andrew McPherson, Hugo M Horlings, Janine Senz, Leah M Prentice, et al. Genomic consequences of aberrant dna repair mechanisms stratify ovarian cancer histotypes. Nature genetics, 49(6):856–865, 2017.

[39] Ruli Gao, Alexander Davis, Thomas O McDonald, Emi Sei, Xiuqing Shi, Yong Wang, Pei-Ching Tsai, Anna Casasent, Jill Waters, Hong Zhang, Funda Meric-Bernstam, Franziska Michor, and Nicholas E Navin. Punctuated copy number evolution and clonal stasis in triple-negative breast cancer. Nature Genetics, 48(10):1119–1130, October 2016.

[40] Stefan Ivanovic and Mohammed El-Kebir. Modeling and predicting cancer clonal evolution with reinforcement learning. Genome Research, pages gr–277672, 2023.

[41] Zubair Lalani, Gillian Chu, Silas Hsu, Shaw Kagawa, Michael Xiang, Simone Zaccaria, and Mohammed El-Kebir. CNAViz: An interactive webtool for user-guided segmentation of tumor DNA sequencing data. PLoS computational biology, 18(10):e1010614, 2022.

[42] Jacob Househam, William CH Cross, and Giulio Caravagna. A fully automated approach for quality control of cancer mutations in the era of high-resolution whole genome sequencing. bioRxiv, 2021.

[43] Leah L. Weber, Chuanyi Zhang, Idoia Ochoa, and Mohammed El-Kebir. Phertilizer: Growing a clonal tree from ultra-low coverage single-cell dna sequencing of tumors. PLOS Computational Biology, 19(10):e1011544, October 2023.

[44] Sarah Christensen, Juho Kim, Nicholas Chia, Oluwasanmi Koyejo, and Mohammed El-Kebir. Detecting evolutionary patterns of cancers using consensus trees. Bioinformatics, 36(Supplement 2):i684–i691, Dec 2020.

[45] Giulio Caravagna, Ylenia Giarratano, Daniele Ramazzotti, Ian Tomlinson, Trevor A Graham, Guido Sanguinetti, and Andrea Sottoriva. Detecting repeated cancer evolution from multi-region tumor sequencing data. Nature Methods, 15:707–714, 2018.

[46] Sahand Khakabimamaghani, Salem Malikic, Jeffrey Tang, Dujian Ding, Ryan Morin, Leonid Chindelevitch, and Martin Ester. Collaborative intra-tumor heterogeneity detection. Bioinformatics, 35(14):i379–i388, 2019.

[47] Ermin Hodzic, Raunak Shrestha, Salem Malikic, Colin C Collins, Kevin Litchfield, Samra Turajlic, and S Cenk Sahinalp. Identification of conserved evolutionary trajectories in tumors. Bioinformatics, 36(Supplement 1):i427–i435, Jul 2020.

[48] Xiang Ge Luo, Jack Kuipers, and Niko Beerenwinkel. Joint inference of exclusivity patterns and recurrent trajectories from tumor mutation trees. Nature Communications, 14(1):3676, 2023.

[49] Heng Li. A statistical framework for snp calling, mutation discovery, association mapping and population genetical parameter estimation from sequencing data. Bioinformatics, 27(21):2987–2993, 2011.

[50] O Delaneau, JF Zagury, MR Robinson, JL Marchini, and ET Dermitzakis. Accurate, scalable and integrative haplotype estimation. nat commun 10: 5436, 2019.

[51] Zihan Ding, Yanhua Huang, Hang Yuan, and Hao Dong. Introduction to reinforcement learning. Deep Reinforcement Learning: Fundamentals, Research and Applications, pages 47–123, 2020.

[52] Saitou Naruya and Masatoshi Nei. The neighbor-joining method: a new method for reconstructing phylogenetic trees. Molecular Biology and Evolution, 4(4):406–425, July 1987.

[53] David Sankoff. Minimal mutation trees of sequences. SIAM Journal on Applied Mathematics, 28(1):35–42, January 1975.

[54] David Benjamin, Takuto Sato, Kristian Cibulskis, Gad Getz, Chip Stewart, and Lee Lichtenstein. Calling somatic snvs and indels with mutect2. BioRxiv, page 861054, 2019.

[55] Petr Danecek, James K Bonfield, Jennifer Liddle, John Marshall, Valeriu Ohan, Martin O Pollard, Andrew Whitwham, Thomas Keane, Shane A McCarthy, Robert M Davies, and Heng Li. Twelve years of samtools and bcftools. GigaScience, 10(2):giab008, January 2021.

[56] Olivier Delaneau, Jean-François Zagury, Matthew R. Robinson, Jonathan L. Marchini, and Emmanouil T. Dermitzakis. Accurate, scalable and integrative haplotype estimation. Nature Communications, 10(1):5436, November 2019.

[57] Adam Auton, Gonçalo Abecasis, Richard Gibbs, Eric Boerwinkle, Harsha Doddapaneni, Yi Han, Viktoriya Korchina, Christie Kovar, Sandra Lee, Donna Muzny, Jeffrey G. Reid, Yiming Zhu, Eric S. Lander, David M. Altshuler, Stacey B. Gabriel, et al. A global reference for human genetic variation. Nature, 526(7571):68–74, 2015.

[58] Josiah P Hanna, Scott Niekum, and Peter Stone. Importance sampling in reinforcement learning with an estimated behavior policy. Machine Learning, 110(6):1267–1317, 2021.

[59] Roland F. Schwarz, Anne Trinh, Botond Sipos, James D. Brenton, Nick Goldman, and Florian Markowetz. Phylogenetic quantification of intra-tumour heterogeneity. PLoS Computational Biology, 10(4):e1003535, April 2014.

